# Beyond the mean: genetic control of gene expression fidelity and dispersion

**DOI:** 10.64898/2026.04.01.715853

**Authors:** Brendan Jamison, Alexander Chen, Kenneth Barr, Erik McIntire, Xin He, Yoav Gilad

**Author notes:** These authors contributed equally to this work (Brendan Jamison, Alexander Chen). Correspondence should be addressed to Y.G.

## Abstract

For decades, molecular biologists have interpreted gene regulation through measurements of mean gene expression, because they could not resolve regulatory variation among individual cells. The advent of single-cell genomics has now made that variation measurable, revealing pervasive differences in gene expression among apparently similar cells. Whether this variation mainly reflects stochastic noise or an informative regulatory property remains unclear. Here we show that mean-corrected gene expression dispersion is a reproducible and biologically structured feature of gene regulation that reflects regulatory fidelity. In heterogeneous differentiated cardiac cultures, genes with low dispersion are shared across cell types, enriched for housekeeping functions, depleted for expression quantitative trait loci, and more highly connected in transcriptional and protein interaction networks. In a comparative single-cell system spanning human, chimpanzee, and allotetraploid cells, a substantial subset of interspecies differences in regulatory dispersion persists in a shared *trans* environment, indicating that gene expression fidelity is often regulated in *cis*. Our findings establish gene expression dispersion as a genetically encoded dimension of gene regulation that is distinct from mean expression, and places dispersion along a fidelity-plasticity axis with implications for development, disease, and threshold-dependent cellular phenotypes.

## Introduction

Gene expression studies have always sought to explain how biological systems differ from one another, between tissues, across conditions, and among individuals^1–4^. During most of this time, expression measurements were obtained from bulk samples. Every estimate of gene expression, therefore, was necessarily an average across many individual cells, and the only relevant quantity that could be compared across samples was the mean. For decades, our research and our understanding of molecular biology, genetics, and genomics have been built on comparisons of mean expression levels, treating the mean as sufficiently informative to model biology.

The emergence of single-cell transcriptomics changed the scale of measurement, allowing researchers to quantify gene expression in individual cells^5^. This change in measurement resolution enabled a wave of studies focused on the mechanistic basis of transcription itself, including the characterization of transcriptional bursting, stochastic promoter switching, and the dynamics of RNA production and degradation at single-cell resolution^6,7^. These studies established that gene expression is inherently dynamic and stochastic at the molecular level, and that cell-to-cell variability can arise directly from fundamental properties of transcriptional regulation^8^.

Yet, despite this technological advance and the new scale of measurement, most analyses aimed at comparing gene expression between systems have continued to rely on the same mean-based conceptual framework. Even with single-cell data, the standard approach is to aggregate measurements across what are presumed to be homogeneous groups of cells, producing a “pseudobulk” profile that again represents the average expression across many cells^9–11^. In that sense, the advantage of single-cell data largely became the ability to better identify homogeneous groups of cells^12^, enabling comparisons between more precisely defined cell populations. But even at this higher resolution, the analysis still relied on means, derived from pseudobulk profiles rather than from the full distribution of single-cell gene expression levels^13–15^.

At the same time, when we measure gene expression at single-cell resolution, we find that even among apparently homogeneous cell populations, individual cells do not all express each gene at the same level^16–18^. This variability, which we refer to as dispersion of gene expression, has been studied in certain biological contexts. During development and differentiation, for example, changes in cell-to-cell variability have been shown to accompany fate transitions^19,20^. At the mechanistic level, dispersion of gene expression has been linked to transcriptional bursting and RNA turnover dynamics^21–23^.

Still, most analyses aimed at comparing gene expression across biological systems continue to treat cell-to-cell variability primarily as technical or stochastic noise. Consequently, the rich literature on transcriptional dynamics has not been fully integrated into how gene expression is analyzed as a phenotype. In large part, this disconnect reflects the fact that dispersion has been primarily interpreted through a mechanistic lens, as an emergent consequence of transcriptional dynamics such as promoter-specific bursting behavior, rather than being examined as a property of regulatory systems with potential functional significance.

Indeed, most prior studies that considered the functional consequences of regulatory dispersion have examined differences between individuals rather than between individual cells^24,25^. For example, after an immune challenge, the variance in gene expression among individuals often decreases even when the mean across the population does not change, because all individuals mount a similar response^26–28^. Such analyses describe how the distribution of mean expression shifts among people or conditions, but they do not reveal how expression varies among cells within a single sample. In parallel, studies that directly examined cell-to-cell variability have primarily focused on the mechanistic origins of dispersion, including properties of transcriptional machinery^29,30^, promoter architecture^31^, enhancer regulation^32^, and RNA kinetic processes^7^. While these studies have provided important insight into how dispersion arises, they have not explicitly examined regulatory dispersion as a functional property of gene regulation.

The role and functional importance of regulatory dispersion thus remain unclear. Understanding whether dispersion of gene expression represents a biological signal is a fundamental question, because the implications extend to many aspects of biology that are governed by threshold effects. Many genetic and physiological processes operate near thresholds, where a shift in expression can move some cells beyond a critical boundary while others remain below it^33,34^. Traditionally, such threshold-dependent outcomes have been interpreted as consequences of changes in mean expression between individuals or conditions. However, differences in expression dispersion could produce similar effects. Distinguishing whether threshold effects arise from differences in mean or differences in dispersion will fundamentally change how we interpret the biology of thresholds and how we think about mechanisms of variation in health and disease. Although our study focuses specifically on determining whether dispersion is biological rather than technical, the answer to this question carries broad implications for understanding and eventually manipulating biological variability.

## Results

### Estimating mean-corrected dispersion in single-cell gene expression data

We characterized gene expression dispersion in single cell data from cardiac heterogeneous differentiated cultures (cardiac HDCs^35^) of three unrelated individuals. The cardiac HDCs were generated in a single batch and maintained for 14 days. Following differentiation, cells from the three individuals were dissociated and combined in equal proportions prior to single-cell RNA sequencing (**Figure 1A**). After genotype-based demultiplexing and quality control filtering (Methods; **Supplementary Figure S1**), we retained data from 152,223 high-quality cells, with a mean sequencing depth of 87,329 reads per cell.

**Figure 1.**
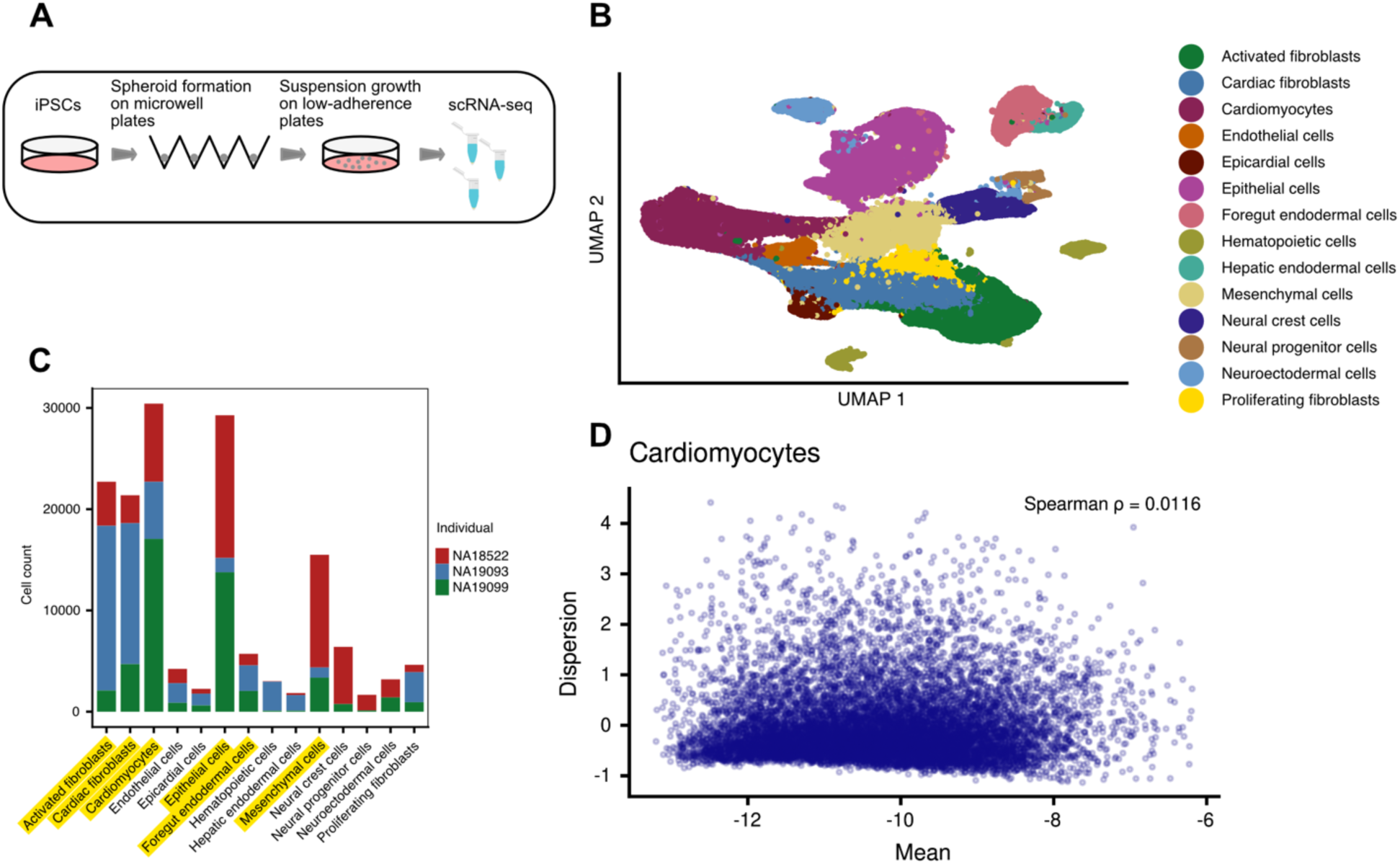
Experimental model and mean-independent dispersion. **A.** Overview of experimental procedure. iPSCs were aggregated into spheroids on microwell plates. After one day, cardiac HDCs were transferred to low-adherence plates and cultured for 12 days. **B.** UMAP projection of all single cells after quality control filtering and reference map-based annotation. **C.** The number of cells per cell type for each individual. Cell types above the threshold of 1000 cells per individual are highlighted in yellow. **D.** Representative sample demonstrating the independence of dispersion from mean expression (ρ = 0.0116). Dispersion and expression estimates are natural log transformed.

Cell type annotation was performed using a label-transfer approach based on a previously characterized cardiac HDC reference dataset containing 14 annotated cell types^35^. Across the entire dataset, 96% of cells were labeled (confidence score greater than 0.8)^36^, and we detected all 14 expected cell types (**Figure 1B**). To have adequate statistical power for estimating gene expression dispersion, we restricted downstream analyses to the six cell types represented by at least 1,000 cells in each individual (**Figure 1C**).

We used Memento^37^ to estimate gene expression dispersion in each combination of individual and cell type (3 individuals, 6 cell types, for a total of 18 samples). Memento estimates the mean and variance of gene expression from the first and second moments of the observed count distribution. These estimates are corrected for technical sampling noise using a hypergeometric sampling model. Because the variance of gene expression also depends on mean expression, Memento fits a polynomial regression of log-variance on log-mean across all expressed genes. The residual from this regression reflects whether expression of a gene is more or less variable than expected given its mean expression. We refer to this mean-adjusted measure of expression variability as dispersion. Across the 18 samples, we estimated dispersion for 11,847 genes with a mean raw UMI of 0.07 or above (the default UMI threshold set by the authors of Memento).

Memento’s estimates of dispersion are not completely independent of mean expression. Genes expressed at very high levels tend to have lower dispersion, whereas dispersion is not associated with the mean across the remainder of the expression distribution. We confirmed that this relationship does not affect any of our subsequent results by performing alternative analyses with gene sets matched for mean expression levels (see below). This relationship might also be biologically valid, as high expression levels may be associated with functional importance and hence low variability across cells. Nevertheless, we excluded from all analyses highly expressed ribosomal genes, and genes with log-mean expression larger than −6 (**Supplementary Tables S1 and S2)**. After this exclusion, dispersion and mean expression are uncorrelated (**Figure 1D; Supplementary Figures S2-S6**).

Dispersion estimates are reproducible across similar cell types collected from different individuals (**Supplementary Figure S7**), indicating that they are unlikely to be dominated by technical noise. We further confirmed reproducibility of the dispersion estimates using a dataset previously collected from cardiac HDCs derived from 46 individuals^35^, where we found that dispersion profiles for similar cell types are more strongly correlated than dispersion profiles for different cell types (*P* < 2 × 10^-16^ for all cell type comparisons; **Supplementary Figures S8-S9**).

### Dispersion exhibits structured patterns across cell types

If dispersion of single cell gene expression is a structured regulated process, genes that perform general cellular functions rather than cell type–specific roles may exhibit similar levels of dispersion across cell types. To examine this, we ranked genes by their median dispersion (across individuals) within each cell type and compared rankings across cell types. We identified 1,033 genes that consistently fell within the lowest quintile of dispersion across all six cell types. Under a model in which dispersion rankings are independent across cell types, fewer than one gene would be expected to overlap across all six cell types.

We found that shared low-dispersion genes tend to have relatively high mean expression levels, possibly consistent with their broad functional importance (recall that in the entire dataset, dispersion and mean are uncorrelated; **Methods; Supplementary Figure S10A**). Gene ontology enrichment analysis of the shared low-dispersion gene set revealed strong enrichment for housekeeping processes, including mRNA processing, protein RNA complex organization, and oxidative phosphorylation (*P* < 10^-10^; Methods; **Figure 2A**). These functional enrichments were robust to alternative clustering resolutions of the data (**Supplementary Figure S11; Supplementary Table S3**).

**Figure 2.**
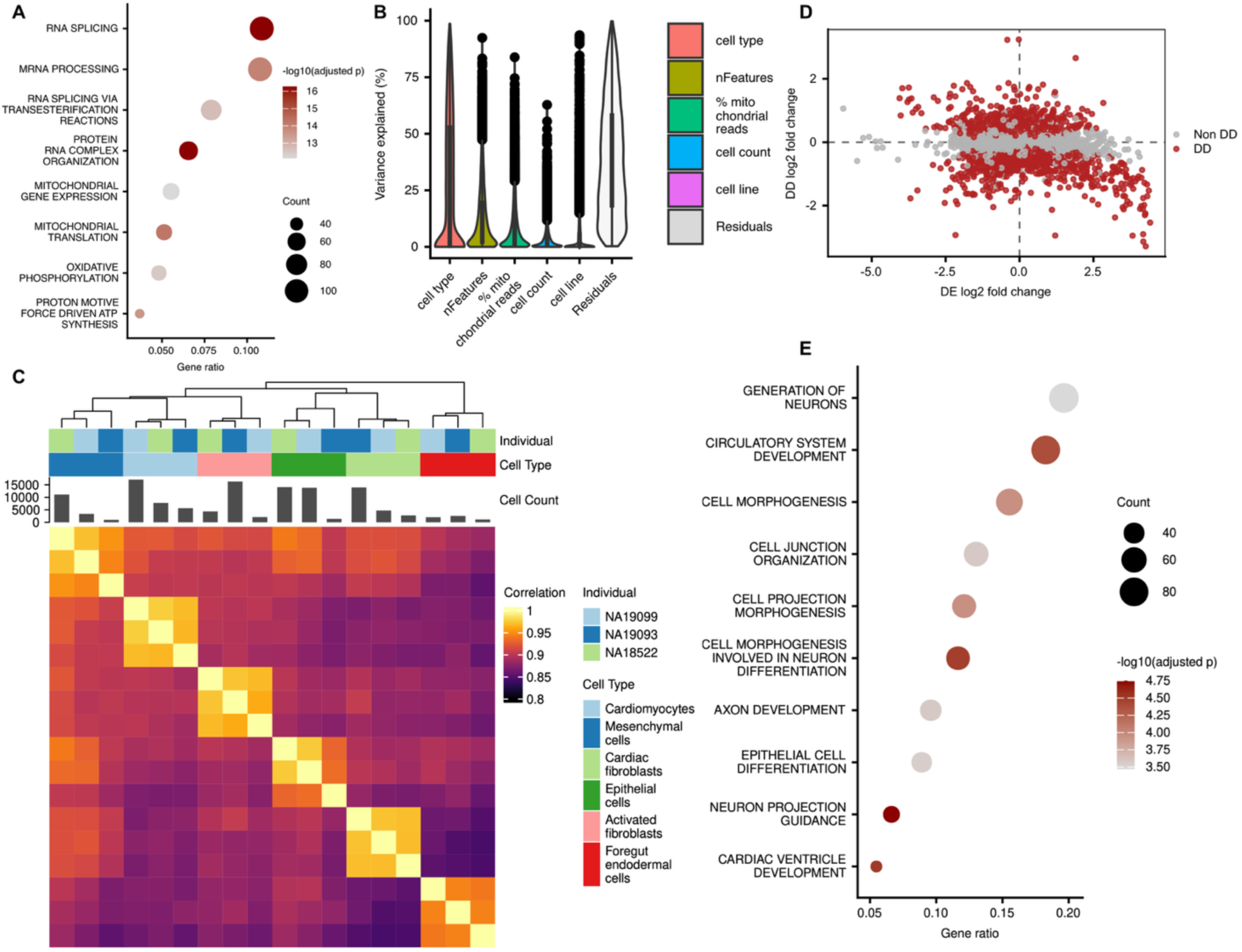
Dispersion patterns are structured across cell types. **A.** Gene Ontology (GO) enrichment analysis for genes ranked in the lowest quintile of dispersion in every cell type. **B.** Variance partition analysis showing the percent of variance in dispersion estimates across samples that is explained by biological and technical variables. **C.** Correlation structure of dispersion profiles across samples. **D.** Relationship between the differential dispersion (DD) and differential mean expression (DE) effect sizes for each gene in the differential tests between cardiomyocytes and foregut endodermal cells. Genes with significant differential dispersion (FDR < 0.05) are colored in red. **E.** GO enrichment analysis of genes that are high DE and low DD in cardiomyocytes relative to foregut endodermal cells. Like terms were clustered together by semantic similarity with GOSemSim^59^, and the term with the lowest BH-adjusted p-value was used as the representative cluster name.

We next focused on genes whose expression dispersion differs across cell types. Using variance partition analysis (**Methods**), we found that cell type accounts for the largest proportion of variance (median of 31.8%) in dispersion estimates in our data (**Figure 2B**). Clustering based on dispersion estimates yields grouping by cell type, similar to clustering based on mean expression levels, though dispersion and mean expression themselves are uncorrelated (**Figure 2C; Supplementary Figures S2-S6 & S12**).

We identified genes with different expression dispersion across cell types (we refer to these as Differentially Dispersed or DD genes) by fitting a linear model with cell type as a fixed effect (Methods). Compared to previous datasets that typically collected ∼10K UMI per cell, our dataset, with roughly ∼35K UMI per cell, is well powered to identify DD genes across a range of cluster sizes (**Supplementary Figure S13**).

We used permutation of cell type labels to obtain a null distribution, calculate empirical *P* values, and estimate false discovery rate (FDR; **Supplementary Figure S14**). At an empirical FDR < 0.05, we identified 1,415 to 3,708 DD genes between pairs of cell types (**Supplementary Table S4**). To further examine the relationship between mean expression levels and dispersion, we also fitted an analogous linear model to mean expression estimates, identifying 2,710 to 6,940 differentially expressed (DE) genes between cell types (**Supplementary Table S4**).

We examined the DD genes between cell types. We focused on the subset of DE/DD genes that show high expression but low dispersion in one cell type compared to the other. This pattern may indicate that, in a particular cell type, these highly expressed genes need to be regulated with high fidelity. Indeed, we found that such genes are enriched with cell-type specific functions (**Supplementary Table S3**). For example, in the pairwise comparison between cardiomyocytes and foregut endodermal cells, genes with higher mean expression and lower dispersion in cardiomyocytes were enriched for circulatory system development, and cardiac ventricle development (*P* < 10^-4^; **Methods**; **Figure 2D-E**).

### Dispersion is associated with regulatory fidelity and functional pleiotropy

Having established that dispersion of single cell gene expression is structured, we next examined its relationship to other properties of gene regulation to assess whether it reflects regulatory fidelity. We first considered the relationship of dispersion with expression quantitative trait loci (eQTLs). If low dispersion reflects high regulatory fidelity, such genes may be less tolerant of regulatory mutations and therefore less likely to be associated with eQTLs.

To test this, we used eQTLs mapped in cardiac-HDC single-cell RNA sequencing data from a sample of 46 individuals (Methods). These data allowed us to map eQTLs in the same cell types in which we estimated regulatory dispersion. We ranked genes by dispersion within each cell type and calculated, along the ranking (using a sliding-window approach; Methods), the proportion of genes associated with eQTLs (eGenes) identified in that cell type. Across cell types, low-dispersion genes were depleted for eGenes, whereas high-dispersion genes were enriched. This pattern was robust across analytical approaches: it was observed when dispersion rankings were analyzed separately within each cell type (**Supplementary Figure S15**), when the dispersion data from all cell types were combined (*P* < 0.05; Methods; **Figure 3A**), and even when eGenes were defined using multivariate adaptive shrinkage (MASH)^38^ to increase power for detecting eQTLs with small effect sizes (Methods**; Supplementary Figure S16**)^39^.

**Figure 3.**
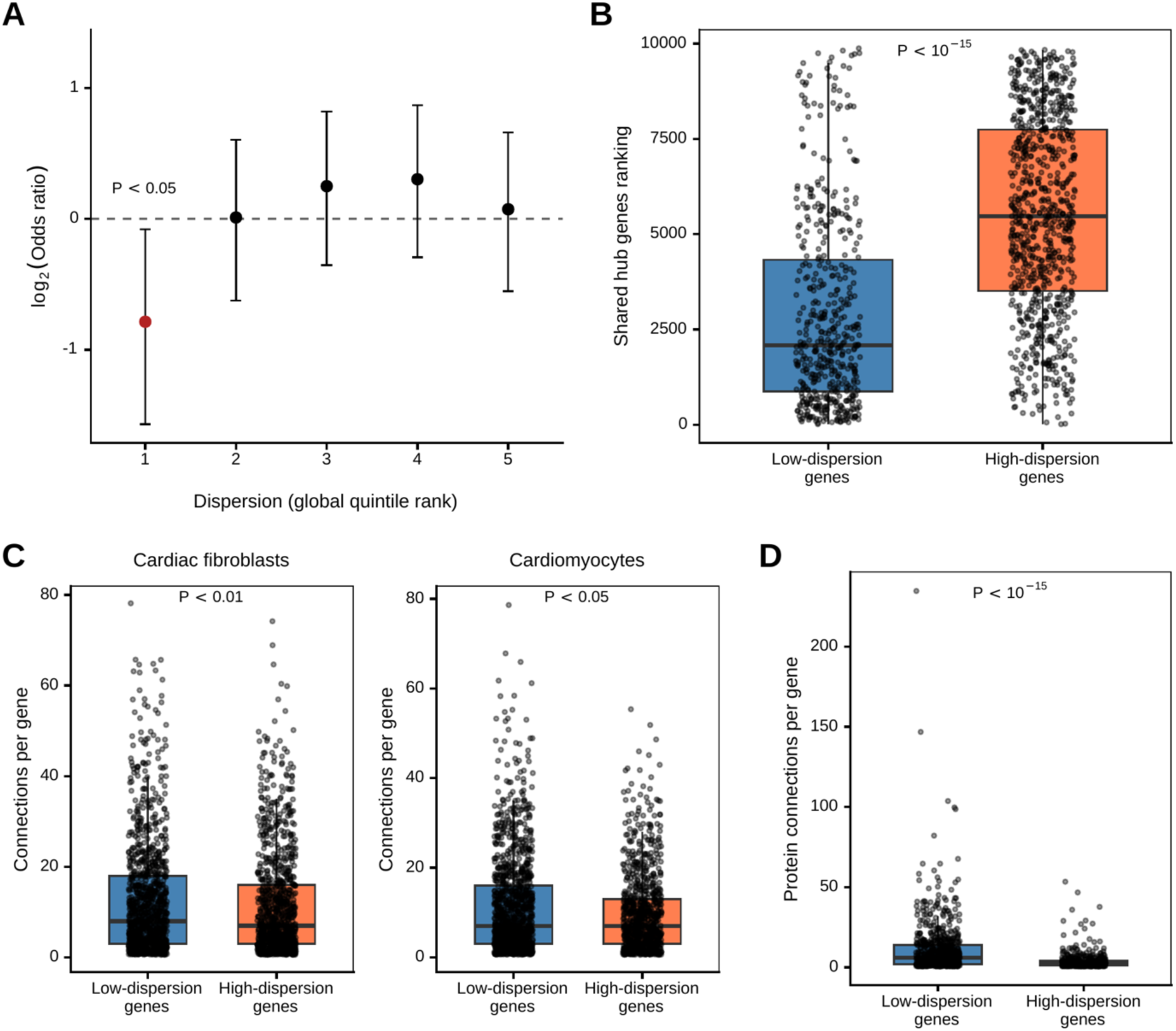
Low dispersion is associated with greater regulatory constraint and pleiotropy. **A.** Enrichment of eGenes in every quintile of the global dispersion ranking. Genes in the lowest quintile are depleted of eGenes (OR = 0.57, 95% CI: 0.33 – 0.92, *P* < 0.05). **B.** Comparison between low-dispersion and high-dispersion genes of hub gene rankings from the shared transcriptional networks of heart and skeletal muscle tissue (*P* < 10^-15^). **C.** Comparisons between low-dispersion and high-dispersion genes of per-gene connections in cell type-specific transcriptional networks. Left panel: cardiac fibroblasts (*P* < 0.01); Right panel: cardiomyocytes (*P* < 0.05). **D.** Comparison between low-dispersion and high-dispersion genes of per-gene interactions at the protein level (*P* < 10^-15^).

We next considered measures of functional pleiotropy. Pleiotropy has been shown to correlate with regulatory constraint. Connectivity within transcriptional and protein interaction networks serves as a proxy for functional pleiotropy. If low dispersion reflects higher regulatory fidelity and constraint, low-dispersion genes would be expected to exhibit greater connectivity.

To examine this, we ranked all genes by their median dispersion across cell types (Methods). Genes at the top of this ranking represent those with consistently low dispersion across the studied cell types. We then considered three sources of data related to gene connectivity: (***i***) transcriptional co-expression networks for heart and skeletal muscle tissues derived from GTEx data^40^, (***ii***) cell type–specific co-expression networks for cardiomyocytes and cardiac fibroblasts derived from the same data set, and (***iii***) protein–protein interaction networks derived from data from the Human Protein Atlas (Methods).

Across all three datasets, we found that low dispersion genes are significantly more connected than high dispersion genes (**Figures 3B-D and Supplementary Figure S17;** *P* < 10^-15^ for comparisons of low and high dispersion genes in transcriptional co-expression networks for heart and skeletal muscle tissues, *P* < 0.05 for comparison in transcriptional co-expression cardiomyocytes and cardiac fibroblasts, and *P* < 10^-15^ for comparisons in protein-protein networks).

We also repeated these analyses after selecting subsets of genes with low and high dispersion whose mean gene expression levels are matched (**Methods**, **Supplementary Figures S18A-D**), to exclude the possible explanation that differences in expression level rather than dispersion drive the observed patterns.

### Regulatory architecture differs between low– and high-dispersion genes

Given the associations between dispersion and measures consistent with regulatory fidelity, we next examined regulatory features that may underlie differences in gene expression dispersion. We first examined regulatory plasticity. Here, we define plasticity as variation in gene expression across tissues, which represent distinct regulatory environments. Under the hypothesis that lower regulatory fidelity is associated with increased plasticity^41,42^, genes with higher dispersion would be expected to show greater variation in expression across tissues.

We used bulk estimates based on the single cell data to identify differentially expressed genes between the cell types (Methods; **Supplementary Table S4**). Across the entire dataset, high-dispersion genes (top quintile) are classified as differentially expressed between cell types (at FDR < 0.05) more often than low-dispersion genes (bottom quintile; *P* < 10^-15^; **Figure 4A**). This observation is robust with respect to the statistical cutoff used to classify genes as differentially expressed, and the specific cell type comparisons that are being considered.

**Figure 4.**
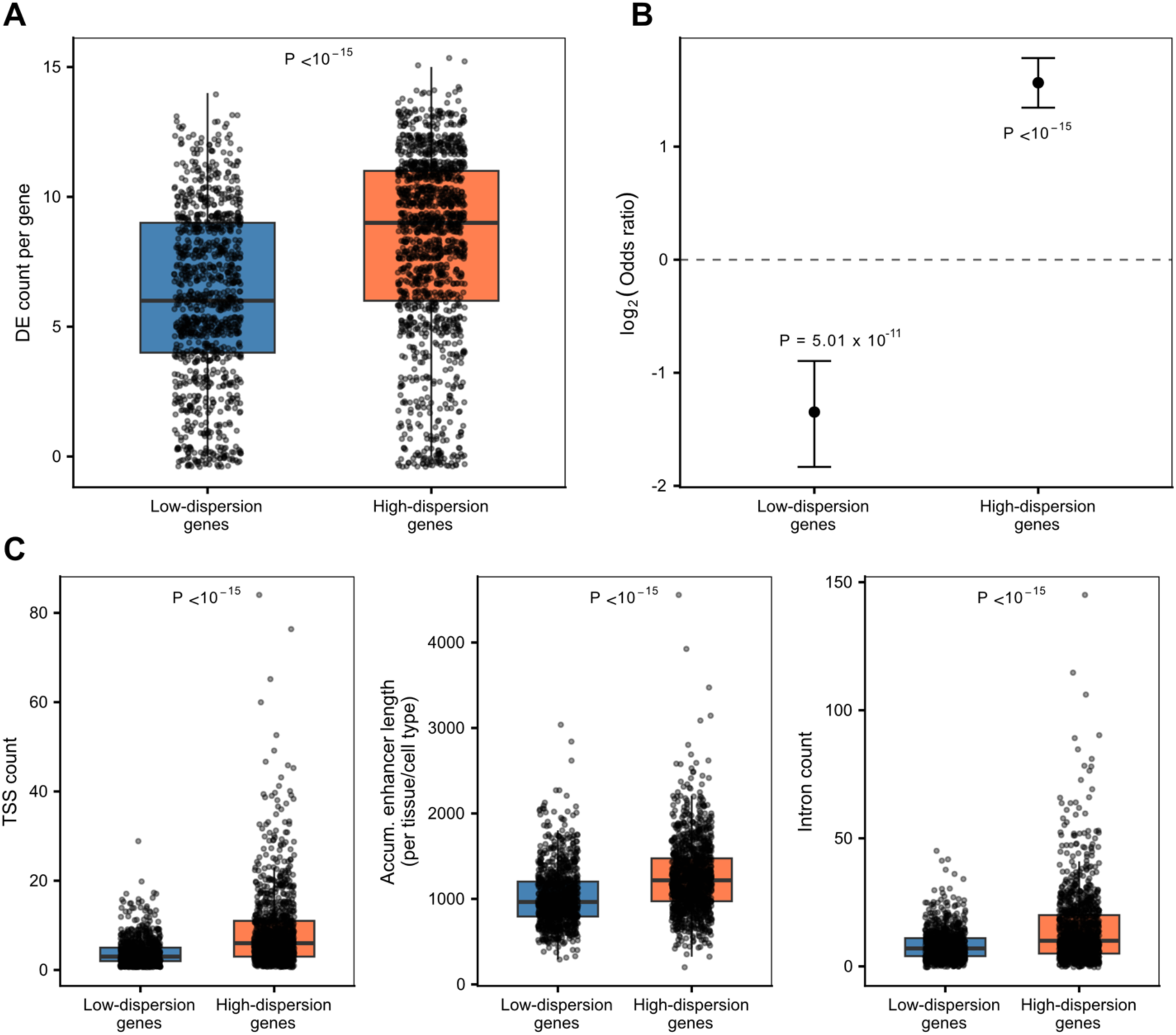
Regulatory architectures differ between low-dispersion and high-dispersion genes. **A.** Comparison of the number of pairwise tests in which low-dispersion and high-dispersion genes are differentially expressed (*P* < 10^-15^). **B.** Enrichment tests of TATA-containing promoters in low-dispersion and high-dispersion genes (low-dispersion genes: OR = 0.39, 95% CI: 0.28 – 0.54, *P* = 4.8 × 10^-11^; high-dispersion genes: OR = 2.99, 95% CI: 2.56 – 3.47; *P* < 10^-15^). **C.** Comparison of the number of transcription start sites (left panel), cumulative enhancer lengths (middle panel), and number of introns (right panel), between low-dispersion and high-dispersion genes.

We next considered different aspects of regulatory architecture. We found that high-dispersion genes are more likely to be associated with TATA-containing promoters (OR = 2.99, 95% CI: 2.56 – 3.47) than low-dispersion genes (OR = 0.39, 95% CI: 0.28 – 0.54; *P* < 0.001; **Figure 4B**). This observation is consistent with previous reports^22,43^. We further found that high-dispersion genes are associated with a higher number of transcription start sites per gene (*P* < 10^-15^; **Figure 4C**; Methods), and a longer cumulative associated enhancer length (*P* < 10^-15^; **Figure 4C**; Methods). Finally, we compared intron counts per gene based on gene annotations (Methods) and found that high-dispersion genes have significantly more introns than low-dispersion genes (*P* < 10^-15^; **Figure 4C**).

Again, we repeated these analyses with subsets of low– and high-dispersion genes matched for mean expression, to confirm robustness of the inference (**Supplementary Figure S18E-I).**

### Robustness of the results with respect to cell state heterogeneity

In all the analyses we have reported to this point, unresolved cell-state heterogeneity presents a particular challenge for interpreting high dispersion. If a nominal cell type contains cells occupying different biological states, genes that distinguish those states will appear highly dispersed even if their expression is tightly regulated within each state. This issue is much less problematic for low-dispersion genes, because unrecognized heterogeneity should generally increase, rather than decrease, dispersion.

To address this, we first confirmed that our observations and classifications of low and high dispersion genes are robust with respect to the choice of cluster resolutions (from 0.5 to 2.0; **Supplementary Figures S19-S20)**, as well as when we consider smaller random samples of the most common cell type in the cardiac HDC dataset (**Supplementary Figure S21)**. We then confirmed that our results are also robust with respect to the exclusion of genes known to be associated with known cell states (such as cell cycle, chamber specificity, and cell type maturation; Methods**; Supplementary Figure S22**).

Finally, we used co-expression analysis to identify and exclude genes that are likely to be associated with both known and unknown cell states. Because a biological cell state should generally be represented by a coordinated transcriptional program rather than by the behavior of a single gene, we reasoned that genes marking unrecognized states should tend to be co-expressed across individual cells. We therefore inferred co-expression modules independently within each cell type and combined them into a consensus set of modules (Methods**; Supplementary Figure S23A–C**). We then ranked modules by the median dispersion of their constituent genes, reasoning that modules representing unresolved cell states should be enriched for high-dispersion genes. Consistent with this expectation, modules corresponding to cell-cycle programs, for example, were consistently among the higher-dispersion modules (**Supplementary Figure S24**).

Rather than defining an arbitrary threshold for classifying modules as cell-state programs, we progressively removed all genes belonging to modules from the high-dispersion end of the distribution and repeated our analyses after each exclusion. High-dispersion genes remained distributed throughout all the ranked modules (**Supplementary Figure S24**). Importantly, the properties distinguishing low– and high-dispersion genes, including TATA-box enrichment, network connectivity, and other regulatory and genomic features, remained evident even after increasingly stringent exclusion of modules most likely to reflect cell-state heterogeneity (**Supplementary Figure S25**). Thus, the major features associated with low and high dispersion cannot be explained simply by unresolved cell-state heterogeneity.

### Gene expression dispersion is often shaped by *cis*-regulatory architecture

We next asked whether gene expression dispersion is regulated by *cis*– or *trans*-acting elements. We addressed this question using gene expression data that were collected in a comparative system in which interspecies regulatory differences can be partitioned into *cis* and *trans* components.

We collected and analyzed single-cell RNA-seq data from unguided HDCs from seven humans, seven chimpanzees, and one human–chimpanzee allotetraploid line (Methods). In the allotetraploid line, the human and chimpanzee genomes are present in the same nucleus. The human and the chimpanzee alleles, therefore, share the same *trans* environment. Using this study design, differences identified between the diploid human and chimpanzee lines that persist between the human and chimpanzee alleles in the allotetraploid line, indicate that the underlying regulatory divergence is encoded in *cis*, reflecting allele-specific effects. In contrast, differences observed between species in the diploid lines that are not recapitulated in the allotetraploid line might indicate that the regulatory divergence is mediated in *trans*, arising from factors that operate through the shared cellular environment.

We estimated mean-corrected dispersion for each gene within each species and cell type, as described above. For each cell type, we identified genes with significant differences in dispersion between human and chimpanzee diploid lines (differentially dispersed or DD genes; Methods). Specifically, for each gene within a cell type, we fit a linear model with species as a fixed effect. In the allotetraploid setting, we used Memento to perform differential dispersion analysis between species, as the experimental setup is optimized for a two-sample comparison. To increase power to detect DD genes, we then used the resulting DD effect sizes and standard errors as inputs for MASH^38^, which leverages shared patterns of effects across conditions to improve statistical power.

Across cell types, we identified between 728 and 2,774 interspecies DD genes per cell type (at LFSR < 0.05; median 1,659 DD genes per cell type; **Supplementary Tables S4-S5**). Across cell types, 5% to 29% of interspecies DD genes identified in the diploid comparison were recapitulated as DD between the human and chimpanzee alleles in the allotetraploid line (**Supplementary Figure S26A; Supplementary Table S5**). The recapitulation rate increases with cell number but plateaus at approximately 25-30% even for the most common cell types (**Supplementary Figure S26; Supplementary Table S6**). Thus, for an appreciable subset of genes with interspecies differences in dispersion, the difference persists in a shared *trans* environment. These observations indicate that interspecies differences in gene expression dispersion are often encoded by *cis*-regulatory sequence divergence between the two species.

We highlight two patterns of DD genes of particular interest that deviate from the expected mean-variance relationship in scRNA-seq data^44,45^. For instance, in stromal cells, *SETD2*, a chromatin modifier that methylates H3K36 during transcription^46^, shows no differential expression between humans and chimpanzees (*P* > 0.8) but is differentially dispersed (*P* < 0.01; **Figure 5A**). Similarly, in neuroblast cells, *SESN3*, which encodes a stress-induced protein that functions as a redox regulator^47^, exhibits comparable expression between species despite significant differences in dispersion (mean expression: *P* > 0.5; dispersion: *P* < 10^-4^; **Figure 5B**). In ENS glia cells, *MAML2*, a transcriptional coactivator that amplifies Notch target gene expression^48^, shows higher mean expression (*P* < 10^-6^) but lower dispersion in chimpanzees compared to humans (*P* < 10^-4^; **Figure 5C**). Together, these patterns indicate that gene expression dispersion reflects a regulatory feature that is distinct from changes in mean expression.

**Figure 5.**
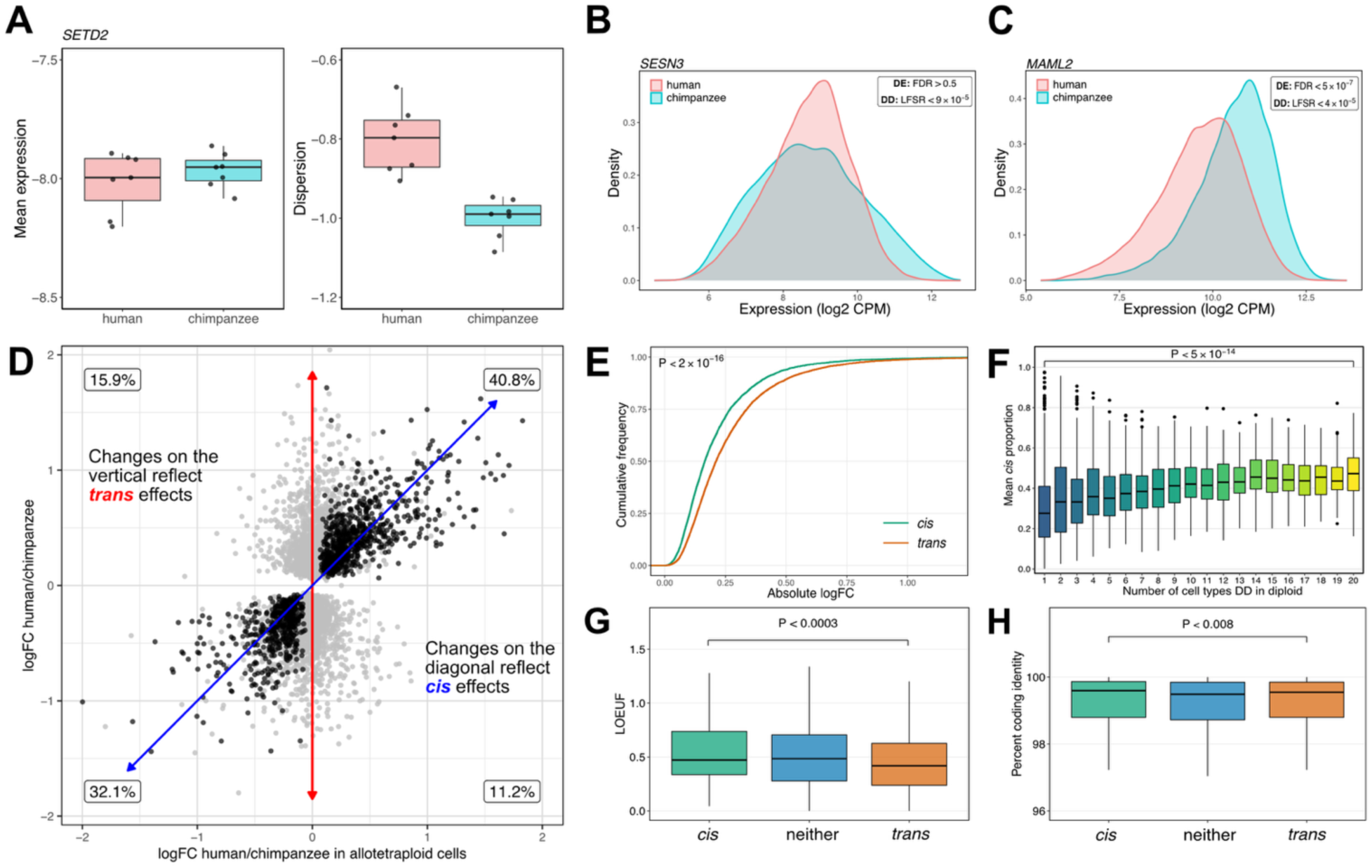
Interspecies differences in regulatory dispersion. **A.** Boxplots show (left) log-mean expression and (right) log-dispersion estimates for *SETD2* in human and chimpanzee stromal cells. Each point represents an individual. **B.** Example of a DD gene that is not differentially expressed (DE) between the species but is differentially dispersed. **C.** Example of a DD gene with higher expression but lower dispersion between species. **D.** DD effect sizes for all cell types between humans and chimpanzees in diploid cells (y-axis) versus allotetraploid cells (x-axis) (LFSR < 0.001). In each quadrant we note the percentage of DD genes. Genes with concordant effect size sign in diploid and allotetraploid cells are shown in black (though this definition underestimates potentially concordant *trans* effects). **E.** Empirical cumulative distributions of effect size for all cell types in diploid cells for tests with high *cis* proportion (> 0.70) and low *cis* proportion (< 0.30). **F.** Boxplot distribution of *cis* proportion for genes DD in the diploid cells for an increasing number of cell types. **G-H.** Boxplots showing the distribution of LOEUF and human-chimpanzee divergence (percent coding identity) respectively, for genes whose interspecies DD can be accounted for by a high, moderate, or low *cis* effect.

To quantify *cis* effects more continuously, we compared the interspecies dispersion effect size between the diploid and allotetraploid lines (Methods). We used this comparison to estimate, for each gene in each cell type, the proportion of diploid DD effect size that is recapitulated in the allotetraploid line between the human and chimpanzee alleles (**Figure 5D**). When the proportion of diploid DD effect size was below 30% or exceeded 70%, we classified the difference in dispersion between the species as arising from interspecies differences in predominantly *trans* or *cis* elements, respectively (Methods; **Supplementary Figure S27; Supplementary Table S7**).

We characterized the differences between *cis*– and *trans*-driving DD genes. We found that when interspecies differences in dispersion are driven predominantly by *trans*-acting elements, the DD effect size between species is larger than when interspecies differences in dispersion are driven predominantly by *cis*-acting elements (*P* < 2 × 10^-16^; **Figure 5E**). However, interspecies differences in dispersion driven predominantly by *trans*-acting elements tend to be more cell-type specific than interspecies DD driven by *cis* (*P* < 5 × 10^-14^; **Figure 5F**). Genes associated with interspecies differences in dispersion driven predominantly by *trans*-acting elements are less tolerant to loss of function mutations than genes where interspecies DD is driven by *cis* (*P* < 0.0003; **Figure 5G**) and have lower coding region divergence between human and chimpanzee (percent ID; *P* < 0.008; **Figure 5H**). These patterns were robust with respect to a range of *trans* and *cis* definitions and cutoffs (**Supplementary Figures S28-S29**) and robust with respect to stricter exclusion criteria of highly expressed genes (**Supplementary Figure S30**).

### Mapping dispersion QTLs

Our comparative analyses suggest that cis-regulatory variation can influence dispersion. We therefore asked whether genetic variants associated with dispersion – *cis* dispersion QTLs – could be mapped across individuals. The available cardiac HDC data from a cohort of 46 individuals (previously mentioned) was originally generated for eQTL mapping and was therefore not designed or powered to detect differences in dispersion, which require substantially more single-cell data. Nevertheless, we reasoned that sufficiently large genetic effects on dispersion might still be detectable.

We therefore performed cis-dQTL mapping separately in each cell type, requiring at least 150 cells per individual for inclusion (Methods**; Supplementary Figure S31**). We tested common variants within 250 kb of each gene’s transcription start site using linear models implemented in MatrixEQTL, with library size included as a covariate, and then jointly analyzed the strongest variant–gene associations across cell types using MASH to leverage sharing of genetic effects across cell types (Methods). Despite the limited power of this dataset for mapping dispersion QTLs, we observed an enrichment of signal for *cis*-regulatory effects (**Figure 6A**). We identified 42 unique dGenes across four cell types at LFSR < 0.1. Importantly, most of these dGenes (32 of the 42) were not detected as eGenes in the same cohort, despite substantially greater power to detect effects on mean expression (**Supplementary Table S8**). For example, we identified a dQTL for *HADHB* in cardiomyocytes in which the less common allele is associated with increased dispersion but no detectable change in mean expression (**Figure 6B**). These results provide additional evidence that genetic variation can contribute to inter-individual differences in expression dispersion independently of effects on mean expression levels.

**Figure 6.**
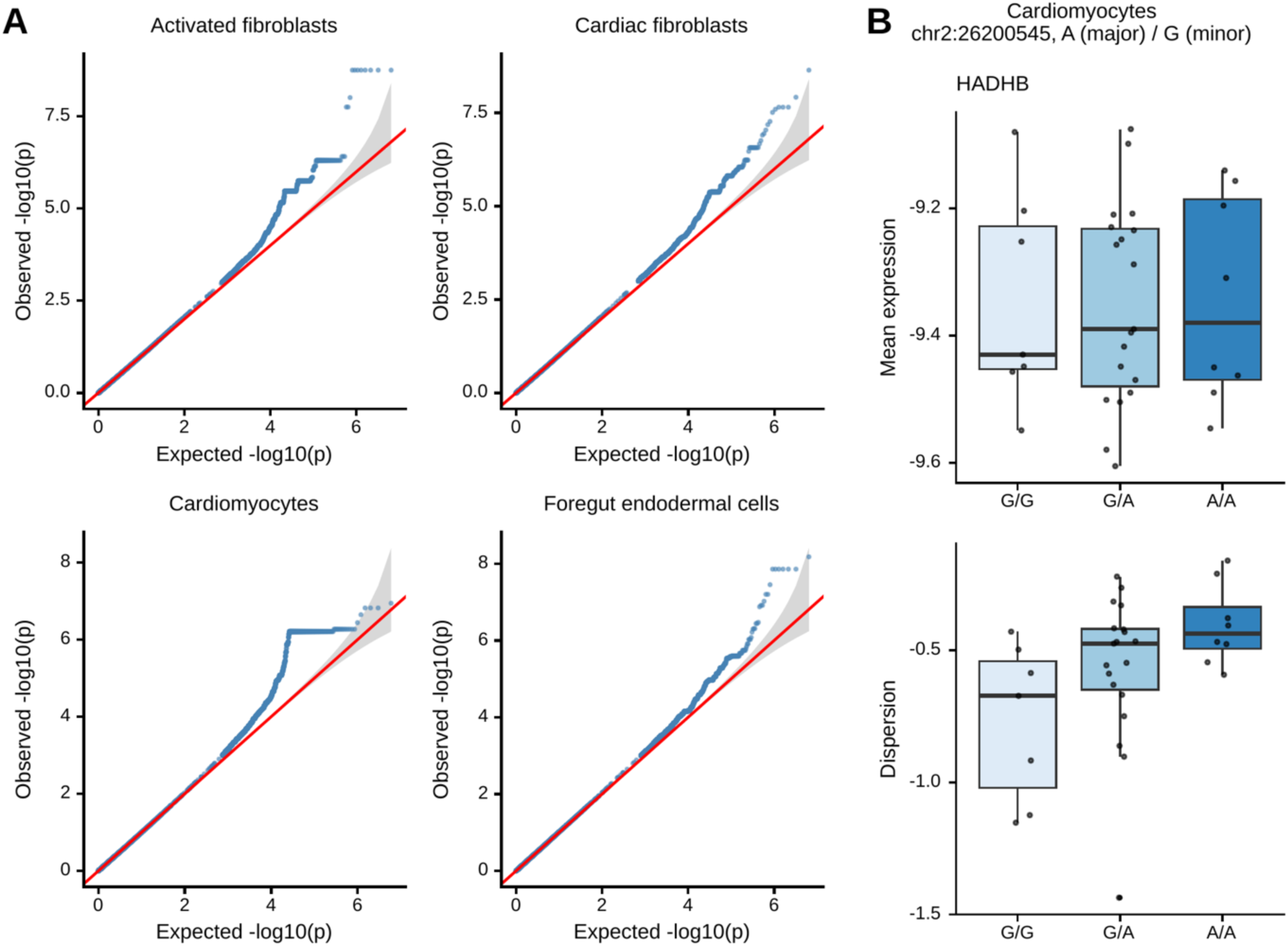
dQTL mapping in the McIntire et al. dataset. **A** Quantile-quantile plots of *cis*-dQTL association p-values for activated fibroblasts, cardiac fibroblasts, cardiomyocytes and foregut endodermal cells. All tests with p < 0.001 are shown, plus a random sample of 3,000 of the remaining tests. **B** Mean expression (top) and dispersion (bottom) of *HADHB* in cardiomyocytes, grouped by genotype (local false sign rate < 0.1).

## Discussion

Gene expression variability has long attracted attention from multiple perspectives. Mechanistic studies established that transcription often occurs in bursts^49^, with promoter switching, polymerase recruitment, and RNA processing kinetics generating cell-to-cell differences in mRNA abundance^6,16,50,51^. In these studies, expression variability was often seen as a byproduct of different transcriptional mechanisms. In parallel, population-level studies showed that gene expression variability across individuals can change following perturbations, such as immune challenge, even when mean expression does not^26,28^. In these studies, variability was generally measured across individuals, not between individual cells.

Here, we frame expression variability, and more specifically, dispersion, as a phenotype that reflects regulatory fidelity. We hypothesized that regulatory dispersion is structured, genetically encoded, and functionally patterned. The evidence supports this view. Dispersion in gene expression shows reproducible, gene-specific structure across cell types. Genes involved in core cellular processes exhibit consistently low dispersion, whereas genes associated with cell-type–specific programs display context-dependent dispersion. Dispersion correlates with regulatory constraint, network connectivity, and promoter and enhancer architecture. Interspecies comparisons further show that a measurable fraction of interspecies difference in expression dispersion is encoded in *cis*-regulatory sequence. Together, these observations indicate that dispersion, or fidelity, is a dimension of gene regulation subject to genetic control.

The structural and functional features associated with dispersion clarify its biological meaning. High-dispersion genes are more frequently differentially expressed across cell types, consistent with greater regulatory plasticity. These genes are enriched for TATA-containing promoters, multiple transcription start sites, longer enhancer landscapes, and increased intron number. Low-dispersion genes are depleted for these features and enriched for housekeeping functions and network connectivity. Dispersion therefore lies along a fidelity–plasticity axis. This notion is consistent with previous observations^41,42^. Genes requiring stable maintenance of expression exhibit high fidelity and narrow distributions. Genes that must remain responsive to distinct regulatory environments exhibit greater structural complexity and broader distributions. Promoter and enhancer architecture appear to shape not only bursting kinetics but also the tolerated breadth of expression variability.

The distinction between mechanistic origin and regulatory function is central. Transcriptional bursting may provide the molecular substrate for variability^52^. The degree to which variability is constrained; however, differs systematically across genes and contexts and is subject to natural selection. Dispersion thus reflects regulatory pressure related to precision. Some genes require tight control across individual cells.

Several caveats merit consideration. Dispersion is estimated in our study within clusters inferred to represent homogeneous cell types or states. Imperfect clustering is generally expected to inflate dispersion as unrecognized heterogeneity would increase rather than decrease the observed variability. Conclusions regarding low dispersion are therefore conservative. To partly mitigate this concern, we evaluated dispersion across multiple clustering resolutions and a reference-based assignment strategy^36^, as well as excluded genes in modules enriched for high dispersion. The principal patterns remain robust.

In the comparative system, *trans*-associated differences in dispersion may capture both genetic and non-genetic components of the cellular environment. The comparative system enables unambiguous identification of *cis* contributions, even in a single allotetraploid line, as sequence divergence between human and chimpanzee chromosomes provides the most plausible explanation for allele-specific expression differences. At the same time, the limited power of this system almost certainly means that we fail to detect some genuine *cis* effects, suggesting our estimates of the contribution of *cis* regulation are conservative. Beyond *cis* effects, trans-associated differences likely reflect a mixture of regulatory divergence and broader context (e.g. measurement error, systematic differences in cell culture microenvironments, etc.).

Nonetheless, the *trans*-acting elements we identified as contributing to interspecies differences in dispersion tend to have larger effect sizes and greater cell-type specificity. This pattern is consistent with previous findings for mean gene expression levels^53–55^ and suggests that a some fraction of these *trans* effects are likely to be real. *Trans* regulatory mutations are expected to be more pleiotropic and therefore more likely to be deleterious. Consequently, the *trans* regulatory mutations that persist are not a random sample, but a selected subset enriched for variants that were advantageous, or in some cases effectively neutral. This survivorship bias predicts that fixed *trans* effects may be relatively large, but confined to fewer contexts, where pleiotropic costs are minimized.

Recognizing dispersion as regulatory fidelity expands how transcriptional phenotypes are conceptualized. Mean expression defines the target level of transcription. Dispersion defines the precision with which that level is achieved across cells. Changes in dispersion can alter the fraction of cells that cross functional thresholds without altering the mean. Such effects are invisible in pseudobulk analyses.

This perspective has direct implications for threshold-dependent phenotypes, including disease onset, drug response, developmental commitment, and gene–environment interactions. Outcomes traditionally attributed to shifts in mean expression may instead reflect changes in regulatory fidelity. A narrow distribution concentrates cells near the mean. A broader distribution increases the probability that some cells cross a functional boundary. Whether increased dispersion enhances or diminishes a phenotype depends on the position of the mean relative to the threshold. If the mean lies below a boundary, greater dispersion increases the chance of crossing it. If the mean lies above it, greater dispersion increases the chance of falling below it. Dispersion is therefore neither inherently beneficial nor detrimental; its effect depends on context.

This framework also helps reconcile prior observations in developmental systems. Studies of transcriptional bursting during differentiation have reported that increased variability can be associated either with enhanced plasticity^56,57^ or with reduced stability of terminal states^58^. When variability is viewed solely as a mechanistic byproduct, these interpretations appear inconsistent. When variability is interpreted as regulated fidelity that modulates threshold crossing, both outcomes become plausible. Dispersion can facilitate transitions when a population approaches a threshold from one side and destabilizes states when a population must remain beyond it.

In summary, gene expression dispersion is structured, genetically controlled, shaped by *cis*– and *trans*-regulatory architecture, and aligned with a fidelity–plasticity continuum. Variability is not simply noise generated by transcriptional bursting. It is a regulated property of gene expression that influences how cellular populations approach and cross functional boundaries.

## Acknowledgment

We thank N. Gonzales for editing and providing comments on the manuscript; members of the Gilad lab and Alexis Battle lab for valuable discussions; University of Chicago Research Computing Center for providing computational resources. This work was supported by NIH grant R35GM131726 to Y.G.

## Data and code availability

The single-cell RNA-seq data have been deposited at GEO under accession GSE326116. The single-cell RNA-seq interspecies unguided HDC data have been deposited at GEO under accession GSE348711. All original code, data, and summary statistics presented in this paper have been deposited on GitHub (https://github.com/awchen55/dispersion).

## Author contributions

Y.G. conceived and designed the study. E.M. collected the eQTL data. B.J. collected the single cell data used to estimate dispersion in cardiac HDCs. K.B. collected single cell data for the interspecies analysis. B.J. and A.C carried out the analysis. B.J. A.C and Y.G wrote the paper. X.H. and Y.G. co-supervised the study.

## Methods

### Experimental model and study design

#### IPSC culture

We maintained iPSC cell lines on Matrigel Growth Factor Reduced (GFR) Basement Membrane (354230, Corning) with Essential 8 medium (A1517001, Thermo Fisher Scientific) and Penicillin-Streptomycin (17–602F, Lonza). Cells were grown at 37°C and 5% CO2 to roughly 80% confluency. Thereafter, we passaged cell cultures with a dissociated reagent (0.5 mM EDTA, 300 mM NaCl in PBS) and seeded cells with 10 uM CEPT (79911KIT, Tocris)

#### Cardiac HDC Model

We differentiated and sampled 3 induced pluripotent stem cell lines from unrelated Yoruba individuals from Ibadan, Nigeria (YRI) into cardiac HDCs from following the procedure developed in the Gilad lab^35^. On the 12^th^ day of culture in the differentiation medium, we collected and dissociated cardiac HDC aggregates as previously described. Following dissociation, we counted cells and recorded viability using Trypan Blue and Countess II (AMQAF1000, Invitrogen). We then mixed cells from each cell line into a single suspension in equal proportions and centrifuged cells at 100 g for 3 minutes. Finally, we resuspended cells in 4°C base heart medium for single-cell RNA seq library preparation.

#### Human-Chimpanzee unguided HDC Model

We used 7 human iPSC lines from unrelated Yoruba individuals and 7 chimpanzee iPSC lines. The chimpanzee lines include 6 previously reported lines^60^ as well as one recently generated line (C50170). We generated this line and verified pluripotency by immunohistochemistry using previously established protocols^60^. Karyotyping performed by WiCell confirmed that the line is karyotypically normal.

We formed unguided HDCs in two batches. The first batch contained 4 human and 3 chimpanzee lines. The second contained 3 human and 4 chimpanzee lines. Each species and batch contained at least one individual of each sex. We also generated unguided HDCs from a single human-chimpanzee allotetraploid line^61,62^ (HLI-25) in a third, separate batch. We followed previously established protocols for generating unguided HDCs^54,63^. On the 21^st^ day of differentiation, samples were dissociated, fixed, and frozen at –80°C for external processing via Scale Biosciences QuantumScale RNA 1M (Product Code 840025).

### Single-cell RNA-sequencing

#### Cardiac HDC Model

With the cells collected from our cardiac HDCs, we performed single-cell RNA sequencing (scRNA-seq) with the 10X Genomics Chromium GEM-X Single Cell 3’ Reagents Kits v4 (Dual Index) (PN-1000215, 10X Genomics). We prepared libraries in a single batch. With 7 lanes of a 10X chip, we recovered roughly 30,000 cells per lane. We sequenced the libraries with the Illumina NovaSeq X at the University of Chicago Functional Genomics Core facility, yielding approximately 87,329 mean reads per cell. We aligned reads to the human reference genome (GRCh38) using CellRanger (v8.0.1) and demultiplexed individuals with Vireo (v0.5.6)^64^. We performed downstream processing analyses in R (v4.4.1) / RStudio (v2024.4.2.764) with Seurat (v5.2.1)^65^. We removed cells classified as doublets or unassigned according to Vireo, and we filtered for cells with at least 4,500 expressed genes. These filtering steps resulted in a total of 38,606 expressed genes and 152,223 cells across the three cell lines.

After filtering, we proceeded to cell type annotation. As noted, the cardiac HDC model was previously developed in the Gilad lab, and cell type annotations were rigorously defined both manually with marker gene expression and against multiple reference datasets^35^. Thus, to annotate our cells, we reference mapped to the annotated data in the previous report^35^. We first log normalized raw counts and selected the 5,000 most variable genes with Seurat’s Variance Stabilizing Transformation method (vst). We then scaled variable features with Seurat’s ScaleData function with default settings. After scaling, we applied principal component analysis (PCA), taking the top 50 principal components, and performed clustering with the Louvain algorithm as implemented in Seurat’s FindClusters function at a resolution of 0.2. With clustered cells and a shared reduction space (PCA) between our dataset and the reference data, we performed the reference mapping procedure in Seurat. Briefly, with the top 50 principal components in both the reference and query datasets, we used FindTransferAnchors to define transfer anchors as cells that are mutual nearest neighbors between the datasets. Then, we used the TransferData function to classify our dataset according to the cell type annotations in the reference^36^. Finally, we applied uniform manifold approximation and projection (UMAP) to the data for ease of visualization.

Given concerns about cell state heterogeneity, in analyses described below we implemented the cell clustering procedure at much finer resolutions (0.5, 1, and 2) and identified a distinct epithelial sub-cluster (**Supplementary Figure S32 & S33**). These cells were removed from the main cluster designated as epithelial cells, and all other analyses performed with this dataset were done after removal of the sub-cluster.

#### Human-Chimpanzee HDC Model

Scale Biosciences sequenced the human and chimpanzee libraries using an Ultima Genomics UG100 with a mean sequencing depth of 35,999 reads per cell. We mapped the provided human and chimpanzee reads to hg38 and panTro6 genomes, respectively, using the Scale Biosciences ScaleRna pipeline (v2.1.0). We used human gene annotations from the 10X Genomics CellRanger 2024 GEX annotation set. For chimpanzee gene annotations we lifted this annotation set to the chimpanzee genome via the Comparative Annotation Toolkit (CAT v2.2.1)^66^. We applied CrossFilt (v0.1.2) ^67^ to remove reads subject to cross-species annotation and alignment bias. We demultiplexed samples via Vireo (v0.5.6).

For the allotetraploid line we separated reads into three classes: human allele-specific reads, chimpanzee allele-specific reads, and reads that could not be confidently assigned to either species. We identified allele-specific (ASE) reads by aligning concatenated human-chimpanzee genome and selecting reads that mapped with high confidence (score > 30) to only one species’ genome. Non-ASE reads were defined as the remaining reads that mapped with high confidence (score >30) to either species alone, but not the concatenated genome. We filtered ASE reads via CrossFilt. Finally, we generated three count matrices: human ASE, chimpanzee ASE, and the total expression defined as the sum of human ASE, chimpanzee ASE, non-ASE count matrices (aligned to the human genome). We used ASE matrices for quantitative comparisons between species, and the total read counts for cell filtering and cell type identification.

After alignment, we performed interspecies analysis in R (v4.4.1) using Seurat (v5.3.0) / SeuratObject (v5.2.0). For the diploid dataset, we performed cell filtering by removing cells classified as doublets or unassigned by Vireo and removed cells with fewer than 1,000 expressed genes; after filtering, 35,032 genes were retained across 1,113,159 cells from 14 unique iPSC lines (7 human, 7 chimpanzee) for downstream analysis. For the allotetraploid dataset, we performed cell filtering by removing cells with fewer than 1,000 expressed genes (no Vireo-based filtering was needed, as this is a single fused hybrid line rather than a pooled multi-donor sample); after filtering, 32,970 genes were retained across 158,290 cells from the single allotetraploid (human-chimpanzee fused hybrid) iPSC line (HLI-25) used for downstream analysis. We annotated cell types using the same pipeline as a previous Gilad lab study that rigorously validated cell type annotations in HDCs^68^. Briefly, cell type annotation was performed at the individual cell level using CelliD^69^ a clustering-free multivariate method that applies Multiple Correspondence Analysis (MCA) to extract per-cell gene signatures. Cells were annotated by testing enrichment of these signatures against a curated reference marker list derived from the human fetal cell atlas^70^ and first-trimester brain atlas^71^, yielding 25 high-confidence cell types. To ensure sufficient power to detect differential dispersion, we restricted our analysis to cell types with at least 1,000 cells per species across diploid lines, represented by at least three individuals each contributing a minimum of 100 cells, and at least 300 cells in the allotetraploid line, yielding 20 cell types for the interspecies analysis (**Supplementary Figures S34–S36**).

### Calculation of mean and dispersion (residual variance)

Memento^28^ separates biological from technical sources of cell-to-cell variability by modeling the observed distribution of single-cell expression counts as a multivariate hypergeometric sampling process, reflecting the fact that transcript capture and sequencing sample molecules without replacement. The method includes a capture efficiency parameter, *q*, which models the probability that a given transcript is correctly captured and observed. Using method-of-moments estimators derived from the first and second moments of the hypergeometric distribution, Memento computes, for each gene, a mean and a variance that is already corrected for sampling-induced (technical) noise via *q*; we refer to this bias-corrected variance as “total variability” and report it in **Supplementary Tables S1-S2**.

Because gene expression variance is also expected to scale with mean expression for biological reasons — more highly expressed genes tend to be intrinsically noisier regardless of measurement process — Memento additionally fits a polynomial regression of log-variance against log-mean across genes and computes the residual from this fit as a mean-corrected measure of variability, which we define as dispersion. We report the percentage of total variability attributable to dispersion (“percent variability that is dispersion”) alongside total variability in **Supplementary Tables S1-S2**.

The explicit formulas Memento uses to estimate mean and variance for cell *c* and gene *g*, respectively:

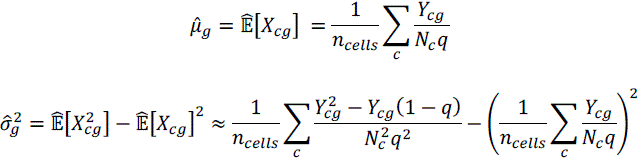

Where μ^_g_ and 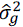 are the estimated mean and variance of the true (pre-sampling) expression of gene *g*. *X* is the true relative mRNA expression and *Y_c_*_g_ is the observed transcript count for gene *g* in cell *c*. *N_c_*is the total transcript count in cell *c*; *n_cells_* is the number of cells; *q* is the capture efficiency parameter, representing the overall probability that a given transcript is captured and sequenced.

We computed gene expression mean, variance, and residual variance (residual variance represents the mean-corrected variance or dispersion/variability) with Memento for each cell type that contained at least 1,000 cells in each individual for the cardiac HDC model. We computed these moments within clusters of cells represented by both cell type and individual. Thus, we made three measurements for each gene in each cell type (3 individuals). To select for genes that are sufficiently expressed for reliable measurement of residual variance, Memento internally filters out genes with a mean raw UMI count less than 0.07 on a per cluster basis. The human cardiac HDC data were generated using fourth-generation 10x GEM-X chemistry, for which we used a capture efficiency parameter of *q*=0.25 because 10x reports higher capture efficiency for this chemistry than for the earlier generations available when Memento was originally developed. Further, to avoid spurious distinctions between cell types driven by power differences, we chose to keep only genes that were expressed above this threshold in all cell type + individual clusters. These filtering steps left us with over 10,000 measured genes.

We applied the same framework in the human-chimpanzee HDC model to estimate mean expression and dispersion. Specifically, estimates were computed for each species, cell type, and individual for cell types with at least 1,000 total cells across individuals and for individuals with at least 100 cells. Estimates were computed for species and cell types with at least 300 cells in the allotetraploid line. The interspecies dataset was generated using the Scale Biosciences QuantumScale RNA platform. A comparative benchmark found that 10x GEM-X detected approximately 24% more genes and 30% more transcripts per cell than Scale^72^, corresponding to Scale recovering approximately 77% of the transcripts and 81% of the genes detected by GEM-X. We therefore used approximately 80% of the cardiac HDC *q* value for the interspecies dataset, resulting in *q*=0.2.

We performed a sensitivity analysis with respect to the choice of *q*. For the cardiac HDC dataset, we repeated all 15 pairwise differential-dispersion comparisons at *q*=0.05, 0.1, 0.15, and 0.2 and compared these results with the baseline *q*=0.25 (**Supplementary Figure S37**). The DD gene sets were highly robust across this range. We also examined the consistently low– and high-dispersion gene sets across *q* values (**Supplementary Figure S38**). 70% of consistently low-dispersion genes and 91% of consistently high-dispersion genes were retained across all *q* values tested, indicating that these classifications are similarly robust to the choice of *q*. For the interspecies dataset, we compared *q*=0.05, 0.1, and 0.15 with the baseline *q*=0.2 and also found that the DD gene sets were highly robust (**Supplementary Figures S39-S40**).

Lastly, we estimated both mean expression and dispersion in the McIntire et al. cHDC cohort for the same 6 cell types as the main cHDC cohort. Given that this cohort was sequenced with the 10X Genomics Chromium Next GEM Single Cell 3’ HT Reagent Kits v3.1 (Dual index, PN-1000348, 10X Genomics), moments with estimated with q=0.2, as this chemistry was previously found to be comparable to Scale in terms of capture efficiency^72^. Cell count thresholds are described in the relevant analyses.

## Analysis using BASiCS

We repeated the classification of low and high dispersion genes using an alternative estimate of dispersion using BASiCS instead of Memento, to confirm that our results can be generalized across different methods to estimate dispersion. Given that BASiCS is known to be computationally intractable in large datasets, we used an updated BASiCS version, BASiCS Divide and Conquer (dBASiCS)^73^ that derives a mean-corrected variance estimate termed residual-over-dispersion, or epsilon. We ran the regression dBASiCS with spike— ins set to FALSE, and we treated individuals as replicates. Further, we down sampled to 1000 cells per individual, and we estimated moments only for genes that were estimated with Memento. For accurate comparisons, we additionally re-estimated moments in the down sampled cell types with Memento. We performed DD analysis using dBASiCS distinct differential testing which call genes differentially dispersed based on a minimum effect size and posterior probability threshold. These parameters were set to default. Results using BASiCS based dispersion estimates are shown in **Supplementary Figures S41.**

## Validation of mean-correction of dispersion estimates

We fit a nonparametric local regression (LOESS) of dispersion against log-mean expression within each sample (species, cell type, and individual; **Supplementary Figures S2-S6**).

Because the LOESS curves still show small deviations from zero at the extreme ends of the expression distribution, we removed the highly expressed ribosomal genes and all genes with log-mean expression > –6 from the analysis. For the interspecies analysis, we additionally removed mitochondrial genes and applied the same log-mean expression cutoff.

We do not mean to imply that dispersion in ribosomal, mitochondrial, or very highly expressed genes is not biologically interesting or should not ultimately be studied. Rather, the primary goal of this paper is to establish that dispersion can be measured as a biological trait and to characterize its structure and regulation independently of mean expression. For that purpose, we considered it more important to work with a set of genes for which we could confidently eliminate the residual relationship between mean and dispersion than to retain every class of gene in the analysis.

Following these exclusions, the LOESS fits are essentially flat across the expression distribution, with only modest deviations at the extreme ends (**Supplementary Figures S2-S6**). Importantly, our primary results are highly robust to these exclusions and thresholds. The consistently low– and high-dispersion gene sets remain nearly unchanged (Jaccard = 0.998 and 0.984, respectively). We also tested a more stringent cutoff in the interspecies analysis by excluding genes with log-mean expression > –7 and found that the inferred patterns of *cis* and *trans* effects on differential dispersion were similarly robust (**Supplementary Figure S30**).

We also directly quantified how much residual mean dependence could contribute to the observed differential-dispersion effect sizes. For each cell type, we fit a LOESS regression of dispersion against average mean expression across the three individuals. For every gene in each pairwise cell-type comparison, we then calculated the dispersion difference expected solely from the gene’s difference in mean expression between the two cell types. For example, for Gene A in a comparison between CT1 and CT2, the expected artifact was calculated as the difference between the LOESS-predicted dispersion at Gene A’s mean expression in CT1 and the corresponding LOESS-predicted value in CT2. Across all differential dispersion tests, the median expected artifact was < 0.01. More generally, the magnitude of the expected mean-related artifact was negligible relative to the observed effect size for dispersion (**Supplementary Figure S42**).

## Gene ontology enrichment analysis

We performed gene ontology enrichment (GO) analysis using the R package clusterProfiler^74^. clusterProfiler computes enrichment tests for a set of genes among GO terms or other biological ontologies based on the hypergeometric distribution. Here, we tested gene sets from pairwise DD tests and rank-based analyses (see below) against GO Biological Process (GO:BP) gene sets within the Molecular Signatures Database^75^ (MSigDB v2025.1.Hs).

We defined background gene sets differently for DD test and rank-based gene sets. For gene sets from DD tests, we were interested in testing low DD genes with respect to all high DE genes. Thus, we defined background genes as high DE genes (FDR < 0.05 and positive log fold change) that overlap with genes in GO:BP gene sets. For rank-based GO enrichments, we defined background genes as those that overlapped between genes in GO:BP gene sets and all measured genes in the ranked gene sets. We corrected for multiple testing with the Benjamini-Hochberg procedure and an FDR cutoff ≤ 0.05 and q-value cutoff ≤ 0.2.

To reduce redundancy in the analysis of low DD and high DE genes cardiomyocytes vs foregut endodermal cells (**Figure 2E**), we additionally hierarchical clustering on GO terms according to their semantic similarity as quantified with the GOSemSim R package^59^. The terms with the lowest BH-adjusted p-values per cluster were used to name the clusters.

In a supplementary analysis, we performed the rank-based GO enrichment test described above for two alternative clustering resolutions (**Supplementary Figure S11**). Thus, rank-based GO enrichments were performed at resolutions 0.2 (cluster resolution for the main cell type annotations), 0.05, and 0.08 as defined in the FindClusters Seurat function (**Supplementary Table S3**)^65^.

### Gene ranking and rank-based analyses

#### Gene Ranking

For all analyses that used rankings of gene expression dispersion, we ranked genes within cell types. There are three measurements per gene per cell type (three individuals for each cell type) in the cardiac HDC data, so we used the median dispersion value per gene across individuals to produce cell type-specific gene rankings.

#### Defining low-dispersion and high-dispersion genes

We considered low– and high-dispersion genes in two contexts for this study: (1) low– and high-dispersion across cell types, and (2) low– and high-dispersion within cell types. Most analyses involved the first context. Here, we binned each cell type-specific dispersion ranking into quintiles. We defined low-dispersion genes as genes in the first quintile (lowest dispersion) across every cell type considered. We defined high-dispersion genes as genes in the fifth quintile (highest dispersion) across every cell type considered.

For low– and high-dispersion genes defined within cell types, we used the lowest ranked 30 percent and highest ranked 30 percent of genes, respectively. In supplementary analyses, we also tested cutoffs at 10 percent and 20 percent (**Supplementary Figure S17**).

In the module inference analyses addressing cell state heterogeneity (**Supplementary Figures S23-S25**), low– and high-dispersion gene sets were defined as the bottom and top 20% of genes by dispersion ranking. Here, ranking was done on the gene universes used for the consensus modules described below (∼5000 genes per cell type).

#### Distribution of low-dispersion genes along mean expression ranking

To confirm that the shared dispersion properties of the consistently low-dispersion and high-dispersion genes were not driven by differences in mean expression, we plotted the distributions of these genes with respect to mean expression. We ranked genes by the median of their mean expression values across cell types, then binned them into quintiles. Within each quintile, we counted the number of low-dispersion and high-dispersion genes, respectively (**Supplementary Figure S10A**). We also equalized the mean expression distributions between the consistently low– and high-dispersion gene sets. We did this by sub-setting the gene set that had the greater number of genes within each bin to match the gene set with fewer (**Supplementary Figure S10B**). We repeated the regulatory and network comparisons described later with the mean-matched gene sets.

### Differential dispersion and differential expression analyses

#### Cardiac HDC Model

We performed differential dispersion analysis between cell types with data from the single batch of cardiac HDCs. We first quantile normalized the residual variance measurements described in the last section. For each sample (cell type and individual) we set cell type as a fixed effect in a standard linear model. Using dreamlet^76^, we fit the model to each gene and defined a contrast matrix representing all pairwise cell type comparisons for hypothesis testing. Finally, we applied empirical Bayes shrinkage and used the Benjamini-Hochberg procedure^77^ to identify differentially dispersed (DD) genes as those with an FDR-adjusted *P* value < 0.05.

We applied the same procedure to test differential mean expression (DE), identifying DE genes with an FDR-adjusted *P* value < 0.05.

#### Comparing power across datasets

We applied the same linear modeling strategy described above in the McIntire et al. cohort. For this data, DD analysis was performed across four cell types: cardiomyocytes, cardiac fibroblasts, activated fibroblasts and mesenchymal cells. These cell types were chosen for two reasons: First, the McIntire et al. dataset was collected over multiple batches, and we wanted to perform the sensitivity analysis within batches to avoid batch effects. Second, the cell types were selected if, within a single batch, they contained more than 1,000 cells per individual.

Once samples (cell type + individual) were selected, we down-sampled to 1,000, 800, 600, 400, and 200 cells per sample. At each step, we estimated dispersion and performed differential dispersion analysis between cell types. As expected, we observed a consistent loss of power to detect differences in dispersion between cell types as the overall number of cells decreased. We then repeated the same analysis in our own cardiac HDC dataset. Again, power consistently dropped with cell count. However, the datasets are not equivalent in one important revealing feature, which reveals the answer to the power question, even when cell counts are similar:

For the cell types tested throughout this study, we captured a median of approximately 35,000 UMIs per cell in our dataset. By comparison, the McIntire et al. cohort captured a more standard ∼9,500 UMIs per cell. Even at the largest cell count step of 1,000 cells per sample, we observed a 4-fold increase in the number of DD genes in our data compared to McIntire et al. Thus, directly addressing the reviewer’s question: we can demonstrate a clear effect of sequencing on the power to detect differential effects which reflects the power to estimate dispersion.

#### Variance partition analysis

We performed variance partition analysis with the variancePartition R package^78^. We tested for the contribution of both biological and technical factors. Following the standard procedure described in the variancePartition documentation, we fit a linear mixed model where cell type was considered as random effects, and cell number, number of features, and percent of mitochondrial reads per sample were considered as fixed effects.

#### Human-Chimpanzee HDC Model

To perform differential dispersion analysis between species in diploid cell lines, we first estimated dispersion using Memento for each sample (species, cell type, and individual). Next, we quantile normalized dispersion and fit a linear model separately for each gene and cell type using dreamlet, with species as a fixed effect. Effect sizes and standard errors from dreamlet were input into MASH^38^, a Bayesian method that models sharing across conditions to improve power and effect size estimates. Significant interspecies DD genes in the diploid line were identified using a local false sign rate (LFSR) < 0.05. We performed differential variability analysis between species in the allotetraploid line using Memento, which is well calibrated to perform hypothesis testing in a two-sample comparison. In an allotetraploid cell, both human and chimpanzee DNA exist in the same *trans* environment. This creates an ideal setting for direct comparisons of dispersion differences without confounding from batch effects. Similarly, effect sizes and standard errors from Memento were input into MASH to improve power to detect DD genes. We identified significant DD genes as genes with an LFSR < 0.05 in both diploid and allotetraploid.

To perform DE analysis between humans and chimpanzees in diploid cells, we generated pseudobulk expression profiles for each sample (defined by species, cell type, and individual) by summing counts across cells. We then used dreamlet to fit a linear model for each gene within each cell type, with species included as a fixed effect. For allotetraploid cells, we performed a paired Wilcoxon signed-rank test comparing, within each cell, the number of reads assigned to human versus chimpanzee alleles. We implemented this test using the row_wilcoxon_paired function from the matrixTests R package^79^. For both analyses, *P* values were adjusted using the Benjamini–Hochberg procedure.

### Dispersion quantitative trait loci (QTL) mapping

#### Power analysis

We knew that the larger cardiac HDC cohort collected by McIntire et al. was underpowered to detect interindividual differences in dispersion. We reasoned that under some cell count threshold, a sample would not contribute informative dispersion estimates for a given genotype group. Therefore, we performed a power analysis to establish a cell count threshold. We sub-sampled the samples included in the DD analysis described above from 1000 cells to 100 in increments of 50 cells. At each step, we repeated the pairwise cell type DD analyses and counted the number of significant DD genes (**Supplementary Figure S31**). We observed that 150 cells per sample was the fewest number of cells with signal to detect DD, and we chose this cell count threshold for dQTL mapping.

#### dQTL mapping

We performed *cis*-dQTL mapping using linear regression models implemented by MatrixEQTL. Mapping was performed separately in each of four cell types: cardiomyocytes, cardiac fibroblasts, activated fibroblasts, and foregute endodermal cells. Each cell type included at least 28 individuals after applying the cell count threshold described above. We used whole-genome sequencing data from the 1000 Genomes Project to map *cis*-dQTLs, removing rare variants with minor allele frequencies < 0.05. We tested variants within a 250kb window of each gene’s transcription start site. We included sex and library size as covariates in the linear models, as well as dispersion principal components (PCs). To determine the number of PCs to include for each cell type, we iteratively mapped dQTLs, successively adding PCs (from 0 to 20) to the linear models at each iteration, and identified the number of PCs that maximized the number of significant (Benjamini-Hochberg FDR < 0.1) dGenes.

After dQTL mapping with MatrixEQTL, we used mashr to address the issue of incomplete power by jointly estimating genetic effects across cell types. We ran mashr using the dQTL summary statistics across all four cell types, including a total of 4,099 genes that were tested in all cell types. Gene-SNP pairs with MAF<0.05 in a given cell type had effects set to 0 and standard errors set to 10^16^ for that cell type, effectively making them uninformative to mashr in that context. To run mashr, we first estimated a residual correlation structure by selecting 9 random SNPs per gene after withholding the gene’s top SNP and finally splitting this set in half for a total of 18,455 random gene-SNP pairs. Next, we estimated data-driven covariance matrices by selecting the top SNP (smallest p-value) per gene across all four cell types. For genes with multiple top hits, we randomly selected one SNP. With this strong set we obtained an empirical covariance matrix, PCA-derived matrices, and matrices computed by empirical Bayes matrix factorization via flashr with non-negative factors and otherwise default settings. The data driven covariance matrices were refined with extreme deconvolution. We fit the mash model to the random set to learn mixture weights on covariance matrices. Finally, we computed posterior statistics for the strong set of 4,099 gene-SNP pairs. For each cell type, gene-SNP pairs with an LFSR<0.1 were designated as significant dQTLs.

### eGene enrichment analysis

#### cardiac HDC collection and eQTL mapping

eQTL data was previously collected in McIntire et al. (unpublished results from cardiac HDCs). Here, we will briefly describe the data collection and eQTL mapping processes.

We differentiated and collected cells from cardiac HDCs from 49 YRI lines using the same cell culture, cardiac HDC differentiation, and collection protocols described above. We sequenced cells from all 49 lines using the 10X Genomics Chromium Next GEM Single Cell 3’ HT Reagent Kits v3.1 (Dual index, PN-1000348, 10X Genomics). We performed alignment, demultiplexing, filtering, and clustering as described above, with some key differences. First, we removed genes expressed in fewer than three cells. Second, we filtered out cells with fewer than 1,500 genes. Following quality control, three individuals were removed, resulting in a final dataset of 46 individuals. We performed cell type annotation as described in McIntire et al.^35^ Finally, for downstream eQTL mapping, we performed pseudobulking within samples (cells represented by cell type and individual). We only kept individuals for pseudobulking in each cell type if they contained at least 15 cells within that cell type. Further, we only kept cell types that contained at least 25 individuals.

We applied linear modeling in MatrixEQTL to map *cis*-eQTLs for each sample. We mapped *cis*-eQTLs with whole-sequencing data from the 1000 Genomes Project, testing SNPs for associations with gene expression within 100kb windows of transcription start sites (TSSs). We excluded SNPs with a minor allele frequency below 0.05.

In a supplementary analysis (**Supplementary Figure S16**) described below, we relied on eGenes identified with MASH. For this set of eGenes, the underlying linear model applied in MatrixEQTL tested SNPs within a 500kb window and included covariates for sex, batch, and cell line passage number at the time of differentiation.

#### Cell type-specific eGene sliding window enrichment analysis

We performed the following analysis first with eGenes identified using MatrixEQTL^39^, then again with eGenes identified after MASH implementation to increase power. We first ranked all measured genes by dispersion within cell types, then, we calculated proportions of eGenes within sliding windows across the dispersion rankings. To compute 95% confidence intervals, for each window we randomly sampled a set of genes equal to the number of eGenes in a given cell type, then calculated the proportion of that gene set represented in the window of ranked genes. We conducted this sampling procedure 50 times for each window.

We restricted the analysis with MatrixEQTL to eGenes with an eQTL discovered at a Benjamini-Hochberg-corrected FDR < 0.05. Further, we tested only four cell types: cardiomyocytes, activated fibroblasts, cardiac fibroblasts, and foregut endodermal cells (an insufficient number of eGenes were detected for this analysis in epithelial and mesenchymal cells). After implementing MASH, a LFSR< 0.05 was used to identify eGenes and all cell types were tested.

#### Global eGene enrichment

The global eGene enrichment analysis differed from the cell type-specific analysis only in the selection of measured genes to rank and eGenes to test. Since we aimed to test for a cell type-agnostic relationship between regulatory variation and dispersion, we limited our analysis to genes with a relatively stable dispersion ranking across cell types, which we identified by computing the median ranking of each gene across four cell types and setting a 15 % threshold that defined how far a gene’s rank could deviate from the median within a cell type. If a gene did not deviate beyond this threshold in any cell type, it was kept for the global enrichment analysis. We then binned these globally ranked genes into quintiles and tested them for enrichment in the eGenes previously identified with MatrixEQTL at an FDR < 0.05, using Fisher’s exact test to quantify enrichment in each quintile. Given the insufficient number of eGenes detected in epithelial and mesenchymal cells, this analysis excluded them both in the ranking of genes and pooling of eGenes.

### Other features relating to the functional significance of dispersion / variability

We compared regulatory characteristics including transcription network complexity and protein interactions between high-dispersion and low-dispersion genes. For analyses on transcription network complexity, we used tissue-specific transcriptome-wide networks (TWNs) and related-tissue TWNs from Saha et al.^40^. We only considered interactions of total expression (TE) levels between genes. For tissue-specific TWNs, we compared the <u>number of TE-TE connections</u> between cell type-specific high-dispersion and low-dispersion genes. For related-tissue TWNs, we compared the <u>ranking of hub genes</u> between high-dispersion and low-dispersion genes across cell types; a lower hub gene ranking indicates more TE-TE connections. Finally, we performed a similar analysis comparing the number of protein interactions per gene between high– and low-dispersion genes, using protein interaction data from the Human Protein Atlas^80–83^.

### Robustness of dispersion estimates to clustering resolution

In the cHDC dataset, we repeated clustering at high resolutions of 0.5, 1 and 2. For comparisons with the full dataset, sub-clusters were kept if they contained at least 700 cells in at least 2 individuals. Mean expression and dispersion were estimated for these sub-clusters as described previously.

### Cell state heterogeneity

#### Co-expression network analysis

Single-cell co-expression network analysis was performed separately on each sample with CS-CORE^84^. Within each sample, the top 5000 genes were selected for network inference according to the method’s depth-normalized mean estimate. CS-CORE was otherwise run with default parameters. Gene-gene correlations with a Benjamini-Hochberg FDR > 0.05 and absolute effect size < 0.25 had effect sizes set to 0 before module inference.

#### Module inference

Module inference was performed with the WGCNA R package^85^, guided by the CS-CORE vignette. First, a soft power threshold was set per sample by iterating from 1-20 with the pickSoftThreshold.fromSimilarity function. Soft thresholds were chosen for each sample by selecting the lowest threshold that reached a scale-free fit R^2^ greater than or equal to 0.85. If this value was never reached, the soft power that gave the best scale-free fit was used (**Supplementary Figure S23A**). Finally, topological overlap matrices (TOM) were computed for each sample using default parameters.

To obtain a single network per cell type, TOMs were combined across the three individuals into consensus modules. For consensus module inference, we filtered genes down to the union genes across individuals. Within cell types, we then calibrated individual TOMs with WGCNA’s single-quantile method. TOMs were combined per element by taking the median value across individuals. Finally, modules were inferred by performing hierarchical clustering on the dissimilarity matrices (1 – TOM) and cutting the trees into module with the cutreeDynamic function in the dynamicTreeCut R package, as recommended in WGCNA documentation.

### Cross-cohort correlation analyses

We tested the reproducibility of dispersion estimates by correlating dispersion profiles in the main cHDC cohort to the McIntire et al. cohort (**Supplementary Figure S8**). Dispersion was estimated in the McIntire et al. cohort for samples with at least 1000 cells. Dispersion profiles were first correlated between the cohorts within cell types. To generate the cell-type-mismatched null expectation (**Supplementary Figure S9**), for each cell type in the main cHDC cohort, all five mismatched cell types in the McIntire et al. cohort were correlated against it and the mean correlation was reported. The null 95% confidence intervals were bootstrapped by sampling genes with replacement in each mismatched pairing, and recomputing the per sample correlation. In each iteration, the rho values across each pair are again averaged. We did this for 2000 iterations. The 95% percent confidence intervals for the matched cell type comparisons relied on the same sampling strategy. A Welch’s t test was used to test for differences between the matched cell type pairs and the null expectation.

### Estimates of *cis* proportion

We estimated the proportion of *cis* and *trans* contributions to gene expression variability divergence by comparing interspecies DD effect sizes of genes in the allotetraploid line to the corresponding effect size in the diploid lines. Taking the absolute value of the log2 fold change between humans and chimpanzees, we calculated the *cis* proportion as abs(allotetraploid.logFC)/(abs(allotetraploid.logFC) + abs(diploid.logFC − allotetraploid.logFC)). This is an adaptation of a procedure used by several previous studies to estimate *cis/trans* contribution to mean expression levels^54,61,62^. We classified a gene as *cis* within a cell type if the estimated *cis* proportion is greater than 70%; if less than 30%, we classified it as *trans*; otherwise, the genes were classified as *neither* (**Supplementary Figure S27**). For each gene, we averaged the estimated *cis* proportion across the cell types in which it was DD in diploid. Genes with a mean *cis* proportion greater than 70% were classified as cis, those below 30% as trans, and all others as neither. Analyses using different cutoffs yielded robust results. (**Supplementary Table S7; Supplementary Figures S28-S29).**

### Characterization of regulatory architecture and constraint

We compared genic features including intron counts, transcription start site (TSS) counts, and per gene cumulative enhancer lengths between genes that were high-dispersion and genes that were low-dispersion across cell types. For intron counts, we used the Ensembl BioMart^86^ and counted introns according to the Matched Annotation from NCBI and EMBL-EBI (MANE) select transcript^87^ for each gene. We calculated TSS counts per gene according to the promoter peaks identified in Cap Analysis of Gene Expression (CAGE) experiments and compiled in the FANTOM5 database^88^. We determined per-gene enhancer lengths by calculating the mean cumulative enhancer length across 131 tissues and cell types as predicted by an Activity-by-contact (ABC) model^89^ (ABC score > 0.05). For testing enrichments of the TATA sequence in gene promoters, we relied on annotations in the Eukaryotic Promoter Database^90^. We obtained the percent coding identity between humans and chimpanzees from BioMart^91^, using chimpanzee as the reference. To characterize the intolerance of interspecies DD genes to loss of function mutations, we obtained loss-of-function observed/expected upper bound fraction (LOEUF) scores from a study of 141,456 human exomes and used scores according to the MANE select transcript with option = ‘true’ for each gene^92^.

### Correlations and *P* values

Correlations reported in the main text were computed using Spearman’s rank-order correlation tests implemented using the R function *corr.test* with option method = ‘s’. To compare effect sizes of *cis* and *trans* effects, a *P* value was calculated using a Kolmogorov-Smirnov test with the R function *ks.test*. For all analyses comparing differences between distributions, we computed *P* values using the two-sided Mann–Whitney *U* test with the R function *wilcox.test*, unless otherwise noted for specific analyses. To test for enrichments in the eGene and TATA enrichment analyses, we used Fisher’s exact test.

## References

1. Umans, B. D., Battle, A. & Gilad, Y. Where Are the Disease-Associated eQTLs? Trends Genet. 37, 109–124 (2021).

2. Ward, M. C. & Gilad, Y. Cracking the regulatory code. Nature 550, 190–191 (2017).

3. Aguet, F. et al. Genetic effects on gene expression across human tissues. Nature 550, 204–213 (2017).

4. Aguet, F. & Ardlie, K. G. Tissue Specificity of Gene Expression. Curr. Genet. Med. Rep. 4, 163–169 (2016).

5. Jovic, D. et al. Single-cell RNA sequencing technologies and applications: A brief overview. Clin. Transl. Med. 12, e694 (2022).

6. Zhang, Q., et al. Transcriptional bursting dynamics in gene expression. Front. Genet. 15, (2024).

7. Kim, J. K. & Marioni, J. C. Inferring the kinetics of stochastic gene expression from single-cell RNA-sequencing data. Genome Biol. 14, R7 (2013).

8. Bax, D. J. C. et al. Gene expression noise in development: genome-wide dynamics. Trends Genet. 0, (2026).

9. L. Lun, A. T., Bach, K. & Marioni, J. C. Pooling across cells to normalize single-cell RNA sequencing data with many zero counts. Genome Biol. 17, 75 (2016).

10. Lun, A. T. L. & Marioni, J. C. Overcoming confounding plate effects in differential expression analyses of single-cell RNA-seq data. Biostat. Oxf. Engl. 18, 451–464 (2017).

11. Crowell, H. L. et al. muscat detects subpopulation-specific state transitions from multi-sample multi-condition single-cell transcriptomics data. Nat. Commun. 11, 6077 (2020).

12. Buettner, F. et al. Computational analysis of cell-to-cell heterogeneity in single-cell RNA-sequencing data reveals hidden subpopulations of cells. Nat. Biotechnol. 33, 155–160 (2015).

13. Lieberman, B. et al. Toward uncharted territory of cellular heterogeneity: advances and applications of single-cell RNA-seq. J. Transl. Genet. Genomics 5, 1–21 (2021).

14. Li, T. et al. An overview of computational methods in single-cell transcriptomic cell type annotation. Brief. Bioinform. 26, bbaf207 (2025).

15. Vieth, B., Parekh, S., Ziegenhain, C., Enard, W. & Hellmann, I. A systematic evaluation of single cell RNA-seq analysis pipelines. Nat. Commun. 10, 4667 (2019).

16. Raj, A., Peskin, C. S., Tranchina, D., Vargas, D. Y. & Tyagi, S. Stochastic mRNA Synthesis in Mammalian Cells. PLoS Biol. 4, e309 (2006).

17. Graf, T. & Stadtfeld, M. Heterogeneity of embryonic and adult stem cells. Cell Stem Cell 3, 480–483 (2008).

18. Altschuler, S. J. & Wu, L. F. Cellular heterogeneity: when do differences make a difference? Cell 141, 559–563 (2010).

19. Mohammed, H. et al. Single-Cell Landscape of Transcriptional Heterogeneity and Cell Fate Decisions during Mouse Early Gastrulation. Cell Rep. 20, 1215–1228 (2017).

20. Shi, J. et al. Dynamic transcriptional symmetry-breaking in pre-implantation mammalian embryo development revealed by single-cell RNA-seq. Development 142, 3468–3477 (2015).

21. Mahat, D. B. et al. Single-cell nascent RNA sequencing unveils coordinated global transcription. Nature 631, 216–223 (2024).

22. Ramalingam, V., Natarajan, M., Johnston, J. & Zeitlinger, J. TATA and paused promoters active in differentiated tissues have distinct expression characteristics. Mol. Syst. Biol. 17, e9866 (2021).

23. Qiu, Q., et al. Massively parallel and time-resolved RNA sequencing in single cells with scNT-seq. Nat. Methods 17, 991–1001 (2020).

24. Shi, G. Genome-wide variance quantitative trait locus analysis suggests small interaction effects in blood pressure traits. Sci. Rep. 12, 12649 (2022).

25. Wolf, S. et al. Characterizing the landscape of gene expression variance in humans. PLOS Genet. 19, e1010833 (2023).

26. Blischak, J. D. et al. Predicting susceptibility to tuberculosis based on gene expression profiling in dendritic cells. Sci. Rep. 7, 5702 (2017).

27. Blischak, J. D., Tailleux, L., Mitrano, A., Barreiro, L. B. & Gilad, Y. Mycobacterial infection induces a specific human innate immune response. Sci. Rep. 5, 16882 (2015).

28. Kim, M. C. et al. Method of moments framework for differential expression analysis of single-cell RNA sequencing data. Cell 187, 6393–6410.e16 (2024).

29. Ren, G. et al. CTCF-Mediated Enhancer-Promoter Interaction Is a Critical Regulator of Cell-to-Cell Variation of Gene Expression. Mol. Cell 67, 1049–1058.e6 (2017).

30. Ali, M. Z., Choubey, S., Das, D. & Brewster, R. C. Probing Mechanisms of Transcription Elongation Through Cell-to-Cell Variability of RNA Polymerase. Biophys. J. 118, 1769–1781 (2020).

31. Sanchez, A., Garcia, H. G., Jones, D., Phillips, R. & Kondev, J. Effect of Promoter Architecture on the Cell-to-Cell Variability in Gene Expression. PLOS Comput. Biol. 7, e1001100 (2011).

32. Liu, T., Zhang, J. & Zhou, T. Effect of Interaction between Chromatin Loops on Cell-to-Cell Variability in Gene Expression. PLOS Comput. Biol. 12, e1004917 (2016).

33. Frontiers | Incomplete Penetrance and Variable Expressivity: From Clinical Studies to Population Cohorts. https://www.frontiersin.org/journals/genetics/articles/10.3389/fgene.2022.920390/full.

34. Raj, A., Rifkin, S. A., Andersen, E. & van Oudenaarden, A. Variability in gene expression underlies incomplete penetrance. Nature 463, 913–918 (2010).

35. McIntire, E., Barr, K. A., Gonzales, N. M., Allen, O. L. & Gilad, Y. Guided Differentiation of Pluripotent Stem Cells into Heterogeneously Differentiating Cultures of Cardiac Cells. BioRxiv Prepr. Serv. Biol. 2023.07.21.550072 (2025) doi:10.1101/2023.07.21.550072.

36. Stuart, T. et al. Comprehensive Integration of Single-Cell Data. Cell 177, 1888–1902.e21 (2019).

37. Kim, M. C. et al. Method of moments framework for differential expression analysis of single-cell RNA sequencing data. Cell 187, 6393–6410.e16 (2024).

38. Urbut, S. M., Wang, G., Carbonetto, P. & Stephens, M. Flexible statistical methods for estimating and testing effects in genomic studies with multiple conditions. Nat. Genet. 51, 187–195 (2019).

39. Shabalin, A. A. Matrix eQTL: ultra fast eQTL analysis via large matrix operations. Bioinformatics 28, 1353–1358 (2012).

40. Saha, A. et al. Co-expression networks reveal the tissue-specific regulation of transcription and splicing. Genome Res. 27, 1843–1858 (2017).

41. Blake, W. J. et al. Phenotypic Consequences of Promoter-Mediated Transcriptional Noise. Mol. Cell 24, 853–865 (2006).

42. Lehner, B. Conflict between Noise and Plasticity in Yeast. PLOS Genet. 6, e1001185 (2010).

43. Sun, M. & Zhang, J. Allele-specific single-cell RNA sequencing reveals different architectures of intrinsic and extrinsic gene expression noises. Nucleic Acids Res. 48, 533–547 (2020).

44. Wills, Q. F., et al. Single-cell gene expression analysis reveals genetic associations masked in whole-tissue experiments. Nat. Biotechnol. 31, 748–752 (2013).

45. Grün, D., Kester, L. & van Oudenaarden, A. Validation of noise models for single-cell transcriptomics. Nat. Methods 11, 637–640 (2014).

46. Molenaar, T. M. & van Leeuwen, F. SETD2: from chromatin modifier to multipronged regulator of the genome and beyond. Cell. Mol. Life Sci. 79, 346 (2022).

47. Rupp, M. et al. FOXO3-mediated chemo-protection in high-stage neuroblastoma depends on wild-type TP53 and SESN3. Oncogene 36, 6190–6203 (2017).

48. Wu, L., Sun, T., Kobayashi, K., Gao, P. & Griffin, J. D. Identification of a Family of Mastermind-Like Transcriptional Coactivators for Mammalian Notch Receptors. Mol. Cell. Biol. 22, 7688–7700 (2002).

49. Blake, W. J., KAErn, M., Cantor, C. R. & Collins, J. J. Noise in eukaryotic gene expression. Nature 422, 633–637 (2003).

50. Herzog, V. A. et al. Thiol-linked alkylation of RNA to assess expression dynamics. Nat. Methods 14, 1198–1204 (2017).

51. Verhagen, P. G. A. & Hansen, M. M. K. Exploring the Central Dogma Through the Lens of Gene Expression Noise. J. Mol. Biol. 438, 169202 (2026).

52. Leyes Porello, E. A., Trudeau, R. T. & Lim, B. Transcriptional bursting: stochasticity in deterministic development. Development 150, dev201546 (2023).

53. Starr, A. L. et al. Disentangling cell-intrinsic and cell-extrinsic factors underlying evolution. Cell Genomics 5, 100891 (2025).

54. Barr, K. A., Rhodes, K. L. & Gilad, Y. The relationship between regulatory changes in cis and trans and the evolution of gene expression in humans and chimpanzees. Genome Biol. 24, 207 (2023).

55. Chen, M. et al. Cell-type-specific eQTLs underlie the genetic architecture of complex traits. Nature 1–8 (2026) doi:10.1038/s41586-026-10577-6.

56. Goolam, M. et al. Heterogeneity in Oct4 and Sox2 Targets Biases Cell Fate in 4-Cell Mouse Embryos. Cell 165, 61–74 (2016).

57. Perera, M. et al. Transcriptional heterogeneity and cell cycle regulation as central determinants of Primitive Endoderm priming. eLife 11, e78967 (2022).

58. Desai, R. V. et al. A DNA repair pathway can regulate transcriptional noise to promote cell fate transitions. Science 373, eabc6506 (2021).

59. Yu, G. et al. GOSemSim: an R package for measuring semantic similarity among GO terms and gene products. Bioinformatics 26, 976–978 (2010).

60. Gallego Romero, I., et al. A panel of induced pluripotent stem cells from chimpanzees: a resource for comparative functional genomics. eLife 4, e07103 (2015).

61. Agoglia, R. M. et al. Primate cell fusion disentangles gene regulatory divergence in neurodevelopment. Nature 592, 421–427 (2021).

62. Gokhman, D. et al. Human–chimpanzee fused cells reveal cis-regulatory divergence underlying skeletal evolution. Nat. Genet. 53, 467–476 (2021).

63. Rhodes, K. et al. Human embryoid bodies as a novel system for genomic studies of functionally diverse cell types. eLife 11, e71361 (2022).

64. Huang, Y., McCarthy, D. J. & Stegle, O. Vireo: Bayesian demultiplexing of pooled single-cell RNA-seq data without genotype reference. Genome Biol. 20, 273 (2019).

65. Hao, Y. et al. Dictionary learning for integrative, multimodal and scalable single-cell analysis. Nat. Biotechnol. 42, 293–304 (2024).

66. Fiddes, I. T. et al. Comparative Annotation Toolkit (CAT)—simultaneous clade and personal genome annotation. Genome Res. 28, 1029–1038 (2018).

67. Barr, K. A. & Gilad, Y. CrossFilt: a cross-species filtering tool that eliminates alignment bias in comparative genomics studies of primates. Genome Biol. 27, 187 (2026).

68. Lin, W. et al. Disease-associated loci share properties with response eQTLs under common environmental exposures. Cell Genomics 0, (2026).

69. Cortal, A., Martignetti, L., Six, E. & Rausell, A. Gene signature extraction and cell identity recognition at the single-cell level with Cell-ID. Nat. Biotechnol. 39, 1095–1102 (2021).

70. Cao, J. et al. A human cell atlas of fetal gene expression. Science 370, eaba7721 (2020).

71. Braun, E. et al. Comprehensive cell atlas of the first-trimester developing human brain. Science 382, eadf1226 (2023).

72. Elz, A. E., et al. Evaluating the practical aspects and performance of commercial single-cell RNA sequencing technologies. NAR Genomics Bioinforma. 8, lqaf215 (2026).

73. O’Callaghan, A. & Vallejos, C. A. Scalable inference of transcriptional variability with BASiCS. J. Theor. Biol. 611, 112157 (2025).

74. Yu, G., Wang, L.-G., Han, Y. & He, Q.-Y. clusterProfiler: an R Package for Comparing Biological Themes Among Gene Clusters. OMICS J. Integr. Biol. 16, 284–287 (2012).

75. Liberzon, A. et al. Molecular signatures database (MSigDB) 3.0. Bioinformatics 27, 1739–1740 (2011).

76. Hoffman, G. E. et al. Efficient differential expression analysis of large-scale single cell transcriptomics data using dreamlet.

77. Benjamini, Y. & Hochberg, Y. Controlling the False Discovery Rate: A Practical and Powerful Approach to Multiple Testing. J. R. Stat. Soc. Ser. B Methodol. 57, 289–300 (1995).

78. Hoffman, G. variancePartition: Quantifying and interpreting drivers of variation in multilevel gene expression experiments.

79. Koncevičius, K. matrixTests: Fast Statistical Hypothesis Tests on Rows and Columns of Matrices. (2025).

80. Uhlén, M. et al. Tissue-based map of the human proteome. Science 347, 1260419 (2015).

81. Yu, N. Y.-L. et al. Complementing tissue characterization by integrating transcriptome profiling from the Human Protein Atlas and from the FANTOM5 consortium. Nucleic Acids Res. 43, 6787–6798 (2015).

82. Lindskog, C. The Human Protein Atlas – an important resource for basic and clinical research. Expert Rev. Proteomics 13, 627–629 (2016).

83. Thul, P. J. & Lindskog, C. The human protein atlas: A spatial map of the human proteome. Protein Sci. Publ. Protein Soc. 27, 233–244 (2018).

84. Su, C. et al. Cell-type-specific co-expression inference from single cell RNA-sequencing data. Nat. Commun. 14, 4846 (2023).

85. Langfelder, P. & Horvath, S. WGCNA: an R package for weighted correlation network analysis. BMC Bioinformatics 9, 559 (2008).

86. Dyer, S. C. et al. Ensembl 2025. Nucleic Acids Res. 53, D948–D957 (2025).

87. Morales, J. et al. A joint NCBI and EMBL-EBI transcript set for clinical genomics and research. Nature 604, 310–315 (2022).

88. Kawaji, H., Kasukawa, T., Forrest, A., Carninci, P. & Hayashizaki, Y. The FANTOM5 collection, a data series underpinning mammalian transcriptome atlases in diverse cell types. Sci. Data 4, 170113 (2017).

89. Nasser, J. et al. Genome-wide enhancer maps link risk variants to disease genes. Nature 593, 238–243 (2021).

90. The Eukaryotic Promoter Database EPD – PubMed. https://pubmed-ncbi-nlm-nih-gov.proxy.uchicago.edu/9399872/.

91. Kinsella, R. J. et al. Ensembl BioMarts: a hub for data retrieval across taxonomic space. Database J. Biol. Databases Curation 2011, bar030 (2011).

92. Karczewski, K. J. et al. The mutational constraint spectrum quantified from variation in 141,456 humans. Nature 581, 434–443 (2020).

