## Supplementary Figures and Tables for "Beyond the mean: genetic control of gene expression fidelity and dispersion"

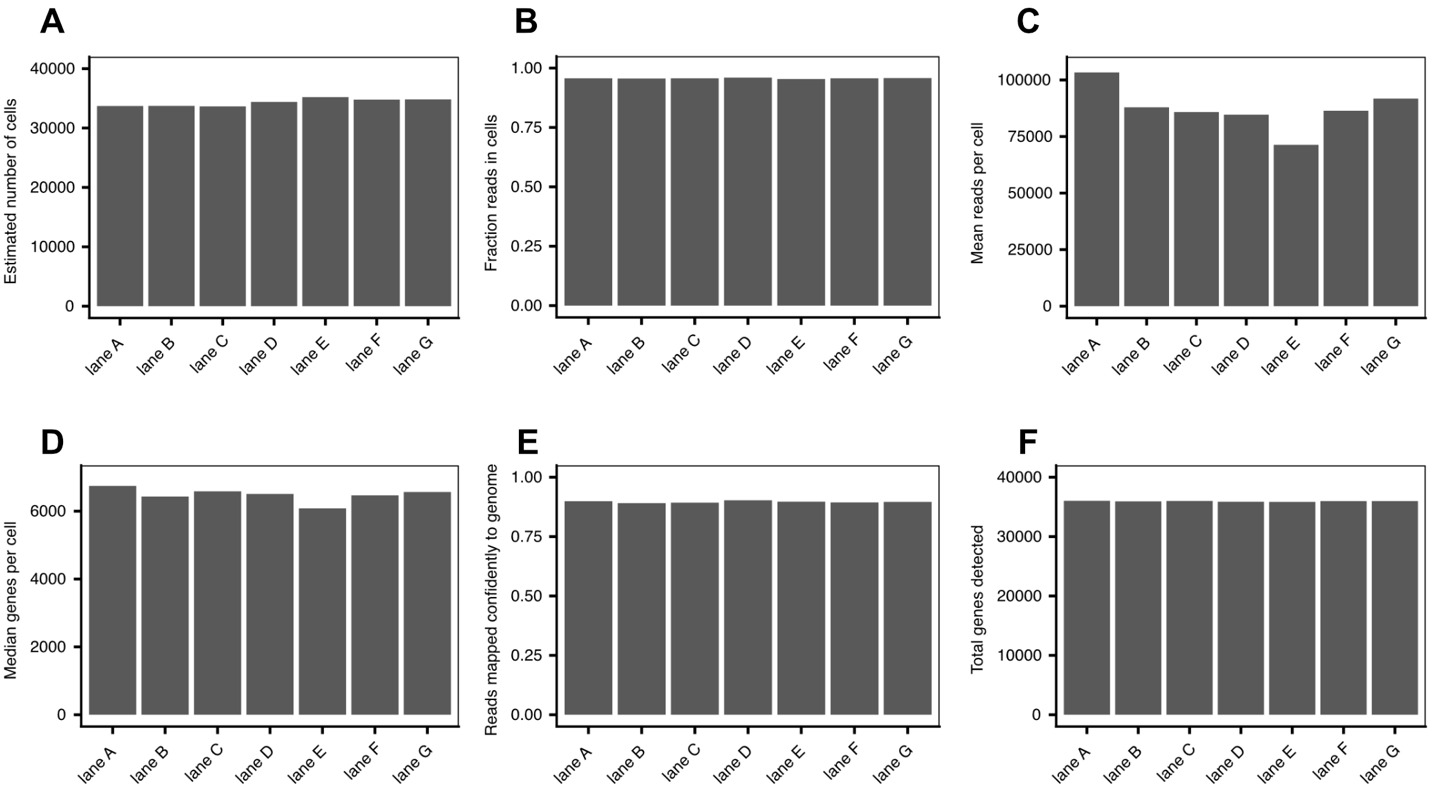

**Figure S1.** Quality control metrics for each single cell RNA-seq library. **A** Estimated number of cells per lane. **B** Proportion of reads in cells per lane. **C** Mean reads per cell per lane. **D** Median number of detected genes per cell per lane. **E** Proportion of genes mapped confidently to the genome per lane. **F** Total number of genes detected per lane.

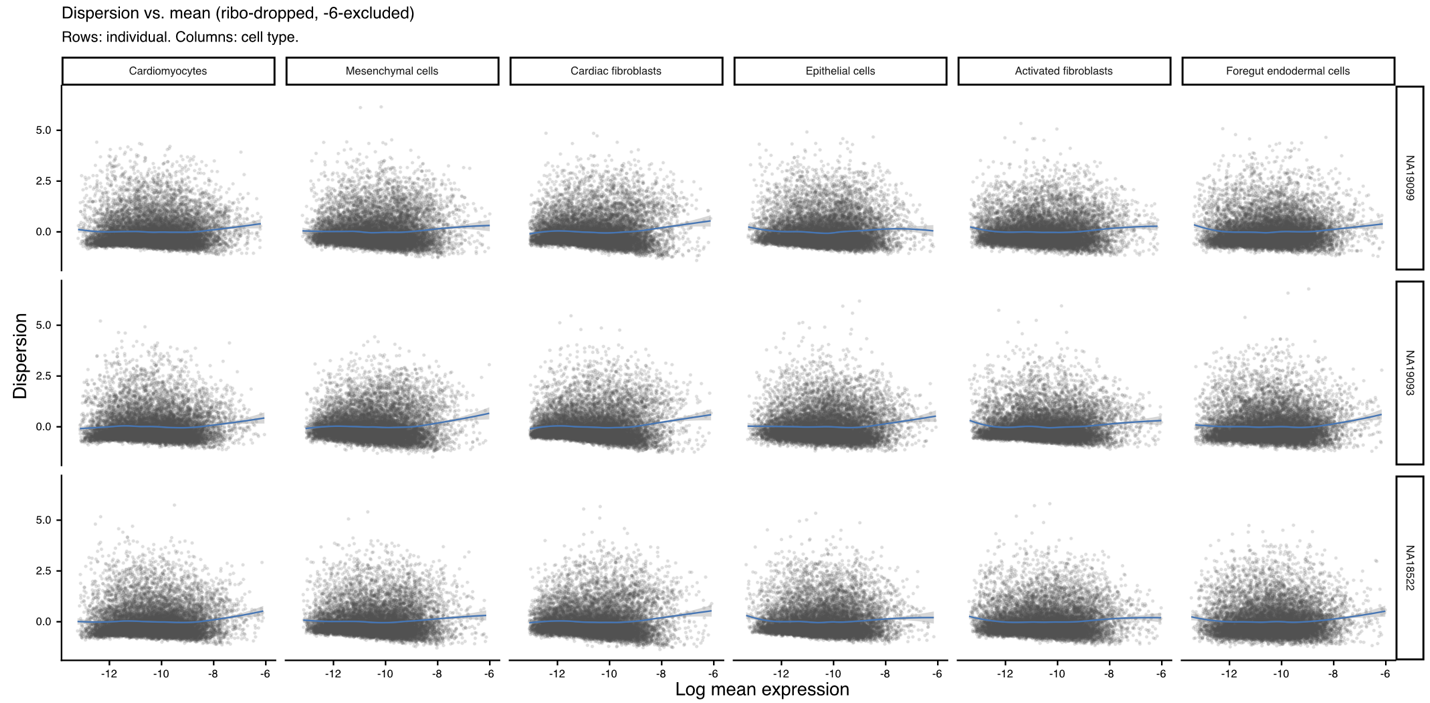

**Figure S2.** Per-individual scatter of log-dispersion (y) vs. log-mean expression (x) for the 6 cardiac cell types (columns) x 3 individuals (rows), on the gene set retained after dropping ribosomal genes and applying a > -6 mean-expression exclusion. Blue line/band = per-panel LOESS fit.

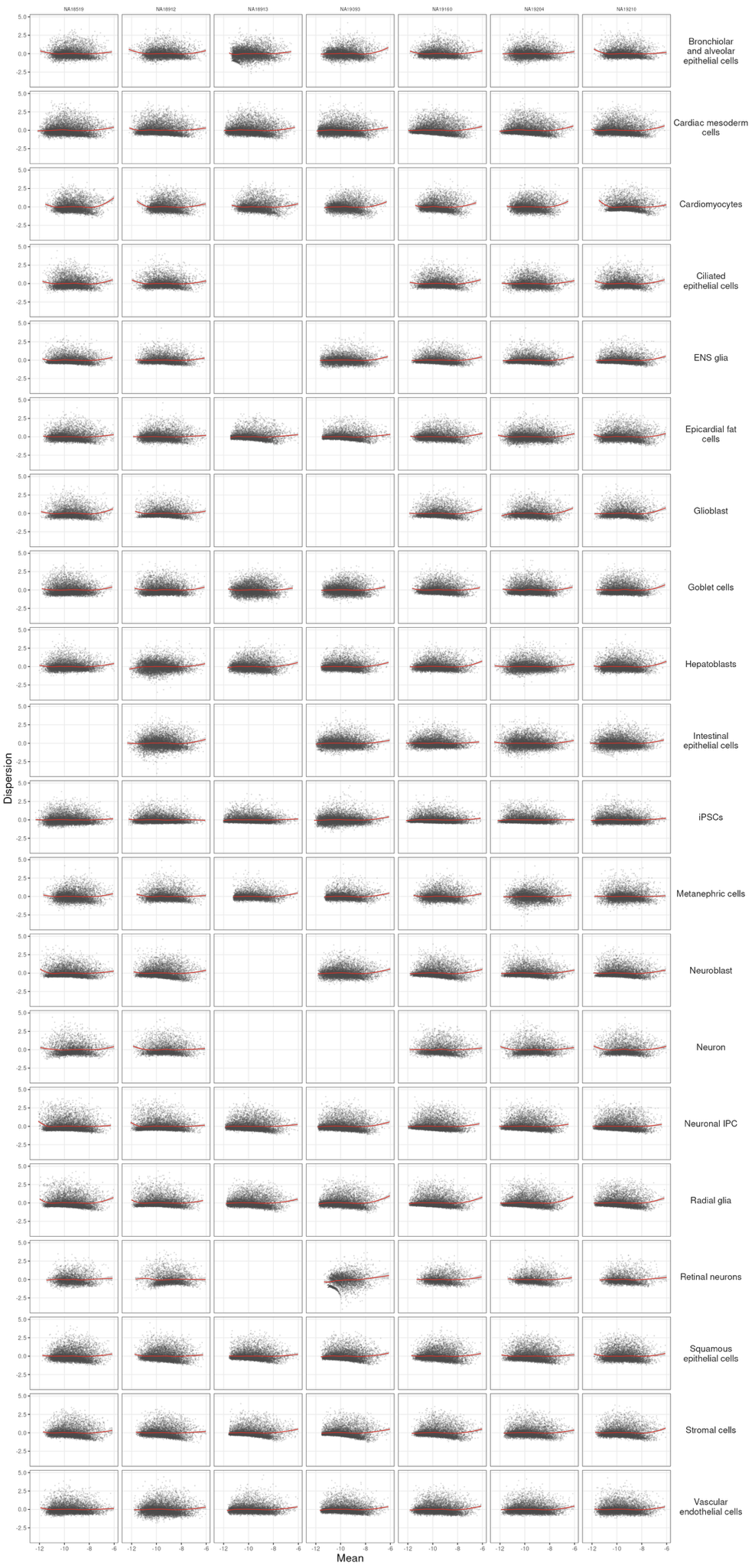

**Figure S3.** Per-individual scatter of log-dispersion (y) vs. log-mean expression (x) for the 20 cell types (rows) x 7 individuals (columns) for diploid human cells, on the gene set retained after dropping ribosomal and mitochondrial genes and applying a > -6 mean-expression exclusion. Red line/band = per-panel LOESS fit.

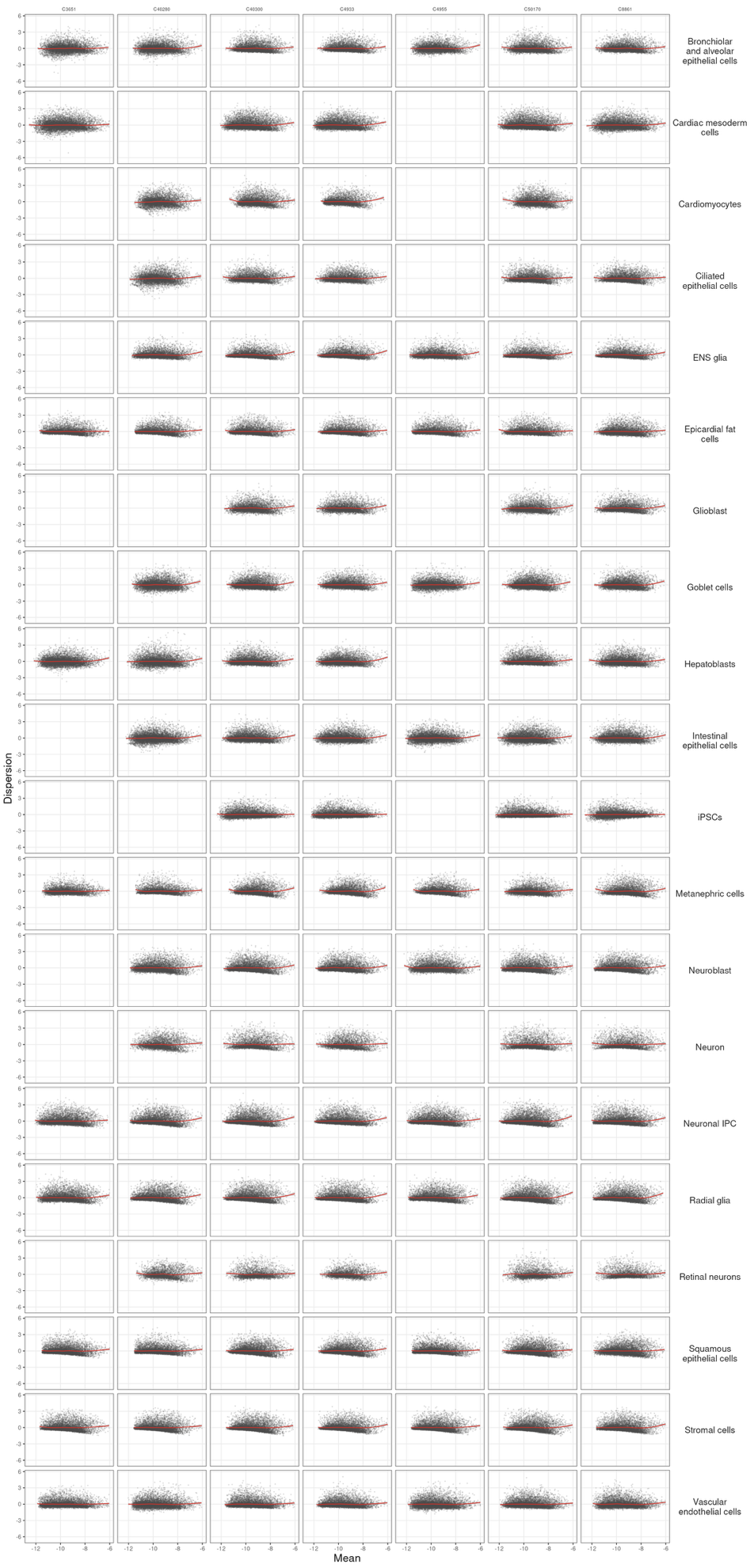

**Figure S4.** Per-individual scatter of log-dispersion (y) vs. log-mean expression (x) for the 20 cell types (rows) x 7 individuals (columns) for diploid chimpanzee cells, on the gene set retained after dropping ribosomal and mitochondrial genes and applying a > -6 mean-expression exclusion. Red line/band = per-panel LOESS fit.

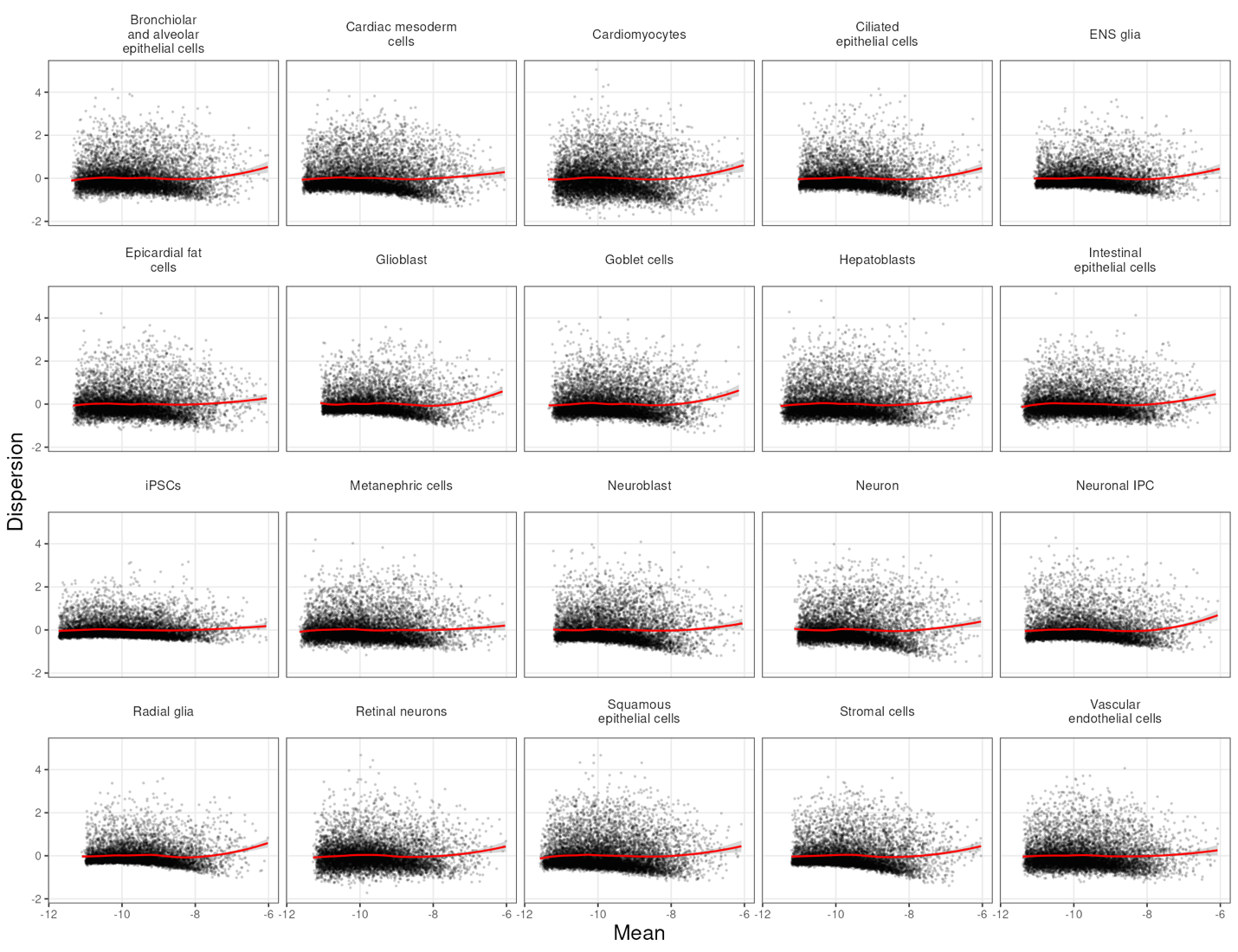

**Figure S5.** Scatter of log-dispersion (y) vs. log-mean expression (x) for the 20 cell types in allotetraploid cells mapped to a human reference after dropping ribosomal and mitochondrial genes and applying a > -6 mean-expression exclusion. Red line/band = per-panel LOESS fit.

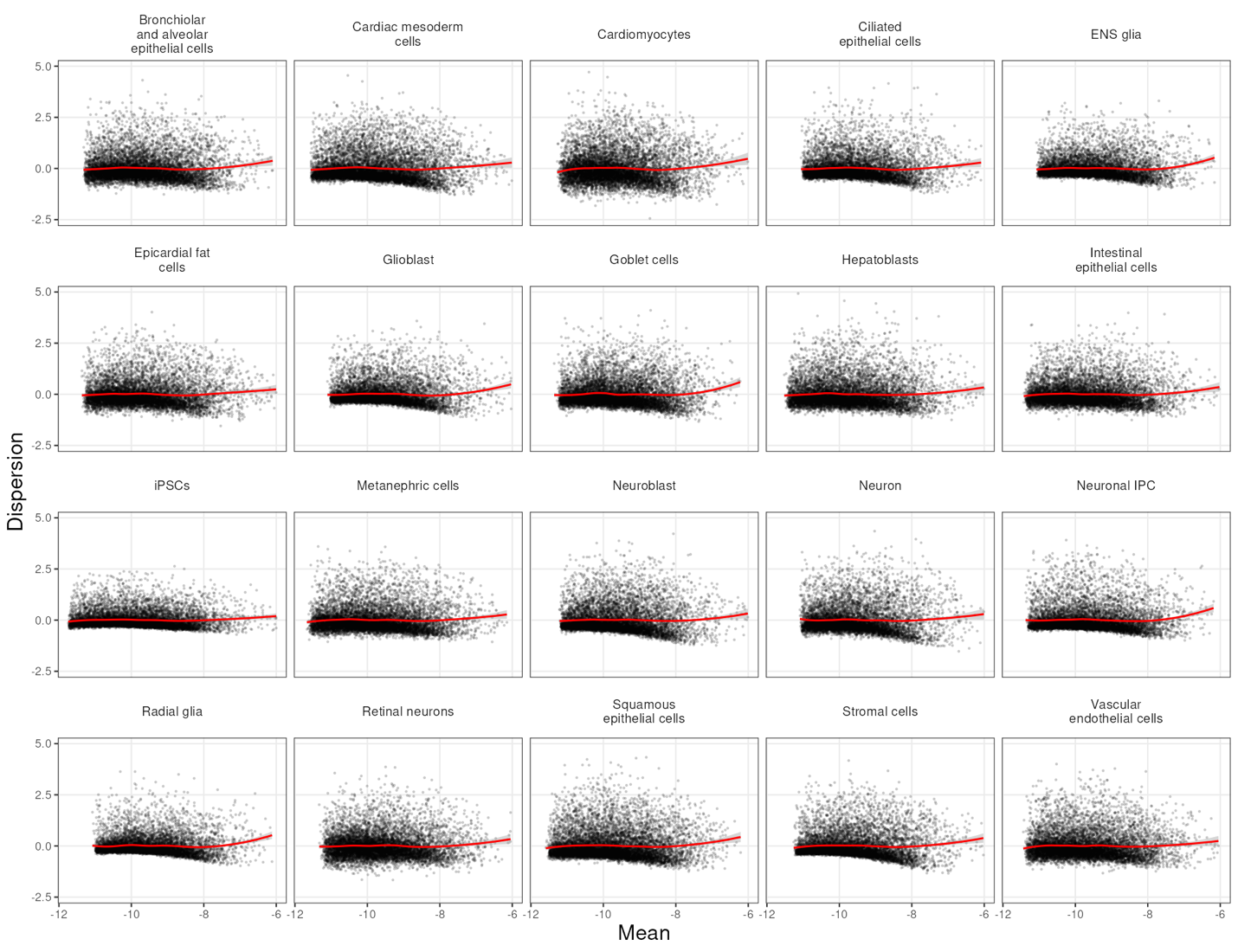

**Figure S6.** Scatter of log-dispersion (y) vs. log-mean expression (x) for the 20 cell types in allotetraploid cells mapped to a chimpanzee reference after dropping ribosomal and mitochondrial genes and applying a > -6 mean-expression exclusion. Red line/band = per-panel LOESS fit.

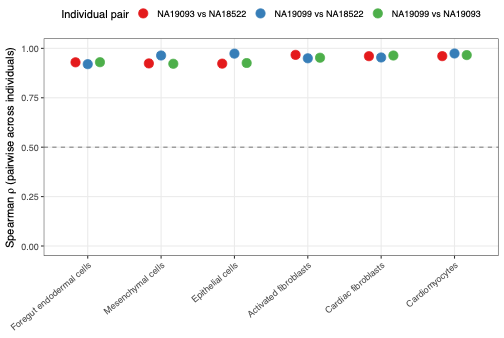

**Figure S7.** Reproducibility of dispersion estimates across individuals and cell types. For the 6 cardiac cell types in the cHDC data, the Spearman correlation of dispersion estimates for every individual pair.

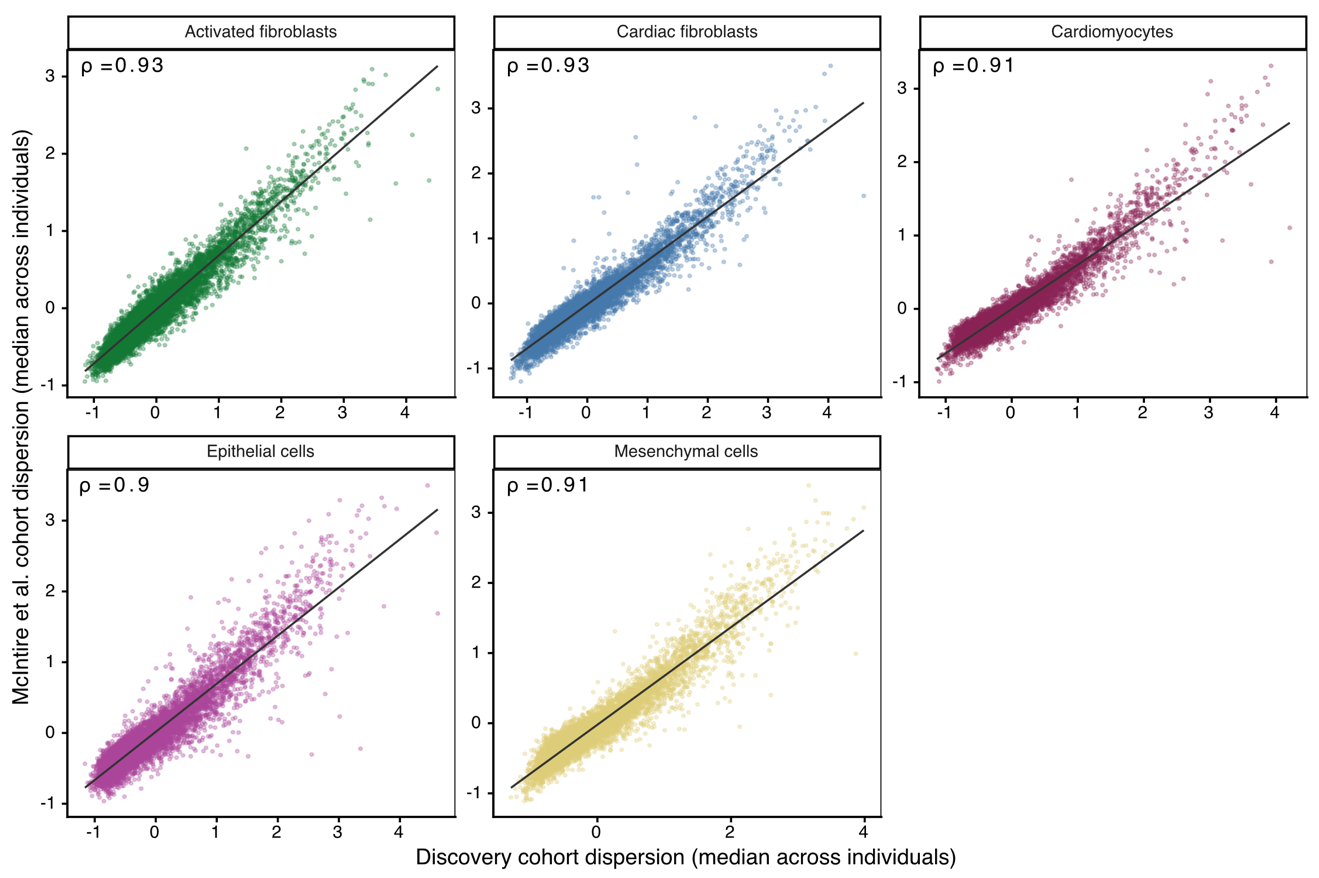

**Figure S8.** Per-cell-type scatter of per-gene dispersion (median across individuals) in this study's discovery cHDC cohort (x) vs. McIntire et al.’s independent cHDC cohort (n=46) (y), at a 1,000-cells-per-individual threshold. Spearman rho = 0.91-0.93 across all 5 shared cell types.

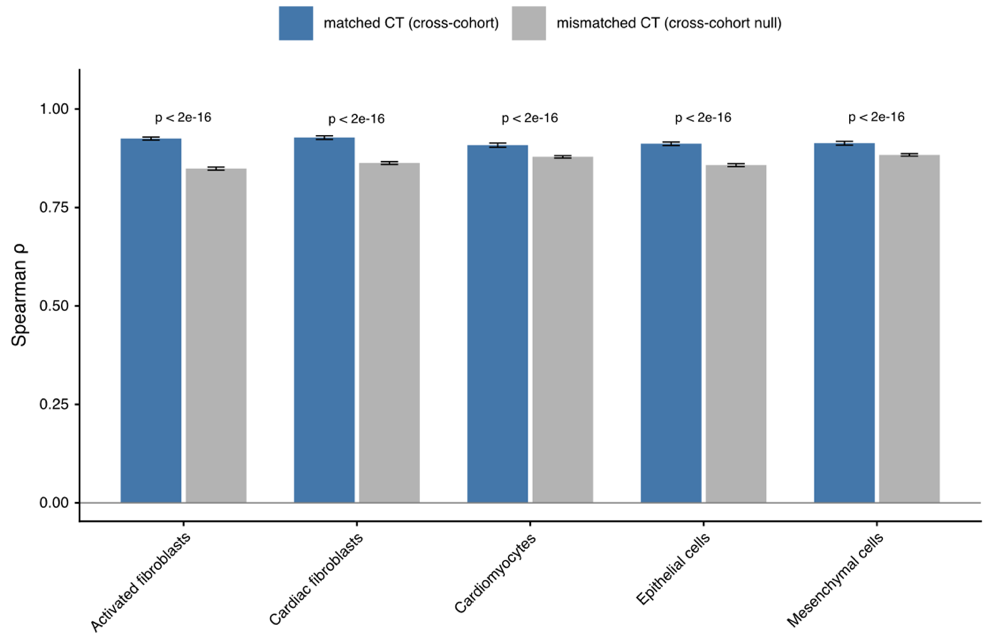

**Figure S9.** Cross-cohort (n=3 cHDC cohort and n=46 cHDC McIntire et al. cohort) Spearman correlation for each cell type's own dispersion estimate ("matched CT", blue) compared to a null built by correlating against a different cell type's estimate in the other cohort ("mismatched CT", gray). Matched-cell-type correlations (~0.90-0.93) significantly exceed the mismatched null (~0.84-0.88) in every cell type (*P* < 2 $\times$ 10^-16^).

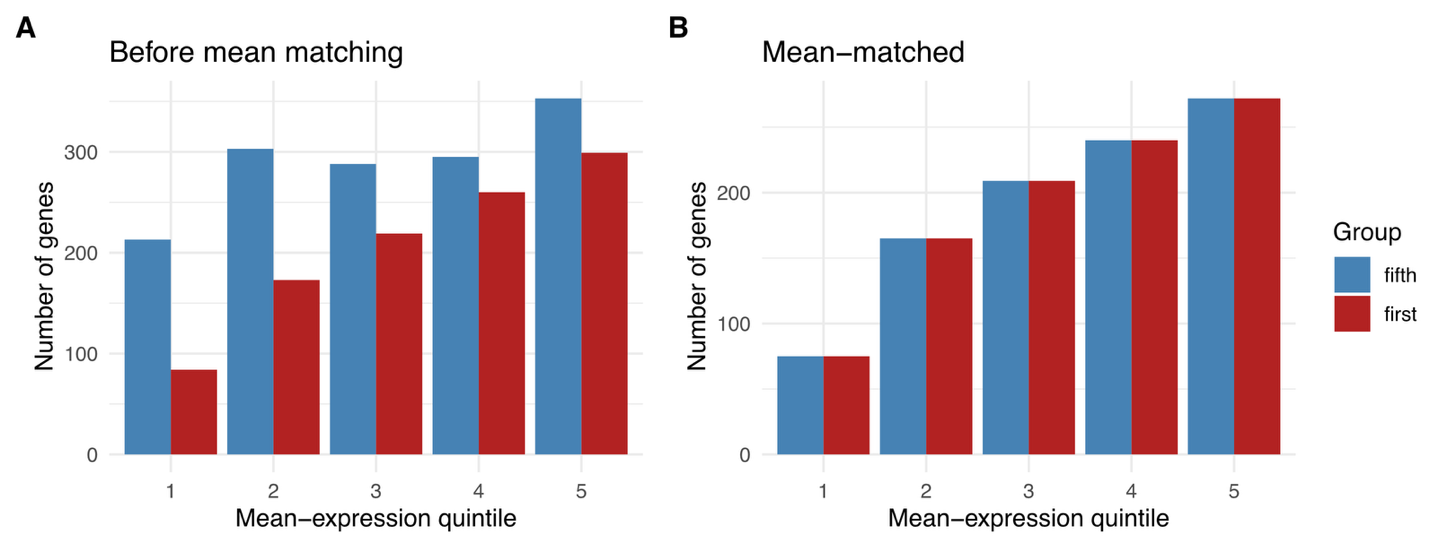

**Figure S10.** Distribution of high and low-dispersion genes across mean expression quintiles. **A.** True distributions. **B.** Distributions after matching mean-expression quintile bins.

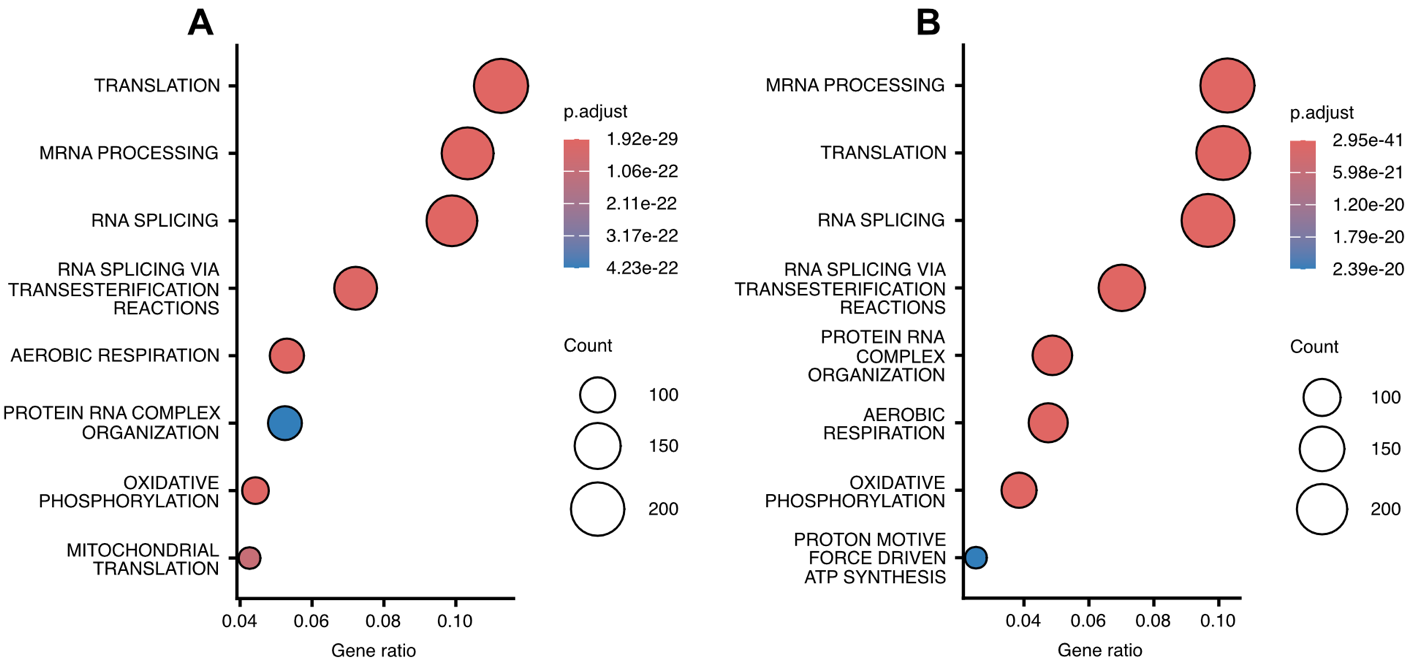

**Figure S11.** Gene ontology (GO) enrichments of low-dispersion genes in alternative clustering resolutions. **A** GO enrichment for genes shared in the first quintile of ranked dispersion across clusters at a resolution of 0.05 (Methods). **B** GO enrichment for genes shared in the first quintile of ranked dispersion across clusters at a resolution of 0.08 (Methods).

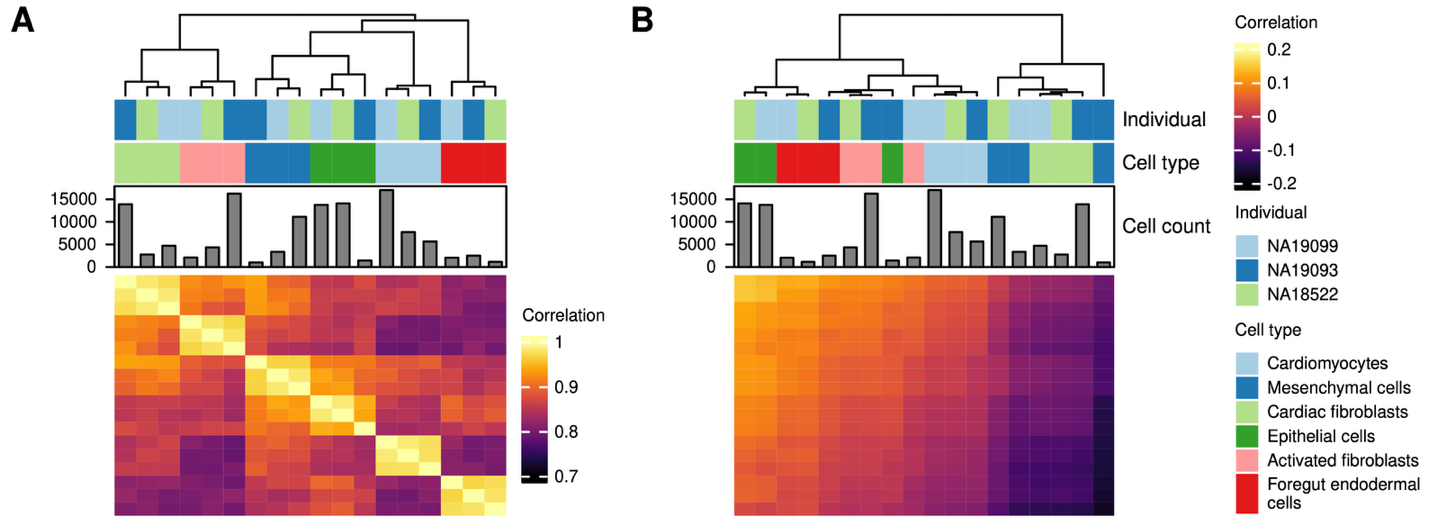

**Figure S12.** Clustering samples by mean expression and mean vs dispersion. **A** Correlation structure of mean expression profiles across samples. **B** Correlation structure of mean expression vs dispersion profiles for each sample.

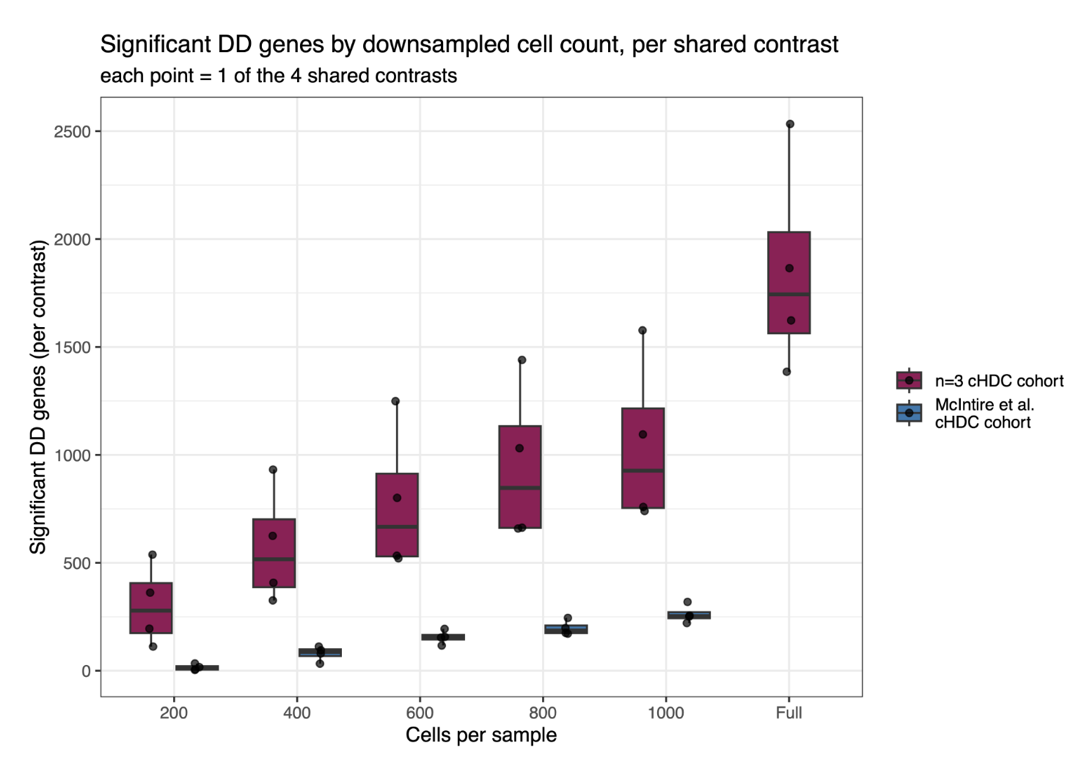

**Figure S13.** Number of significant DD genes (adj.P.Val ≤ 0.05) as a function of downsampled cells-per-individual-per-cell-type, comparing this study's n=3 cHDC cohort (maroon) to McIntire et al.'s independent cHDC cohort (blue), plus n=3 cHDC cohort at full-depth value. Each dot represents a different pairwise cell type comparison.

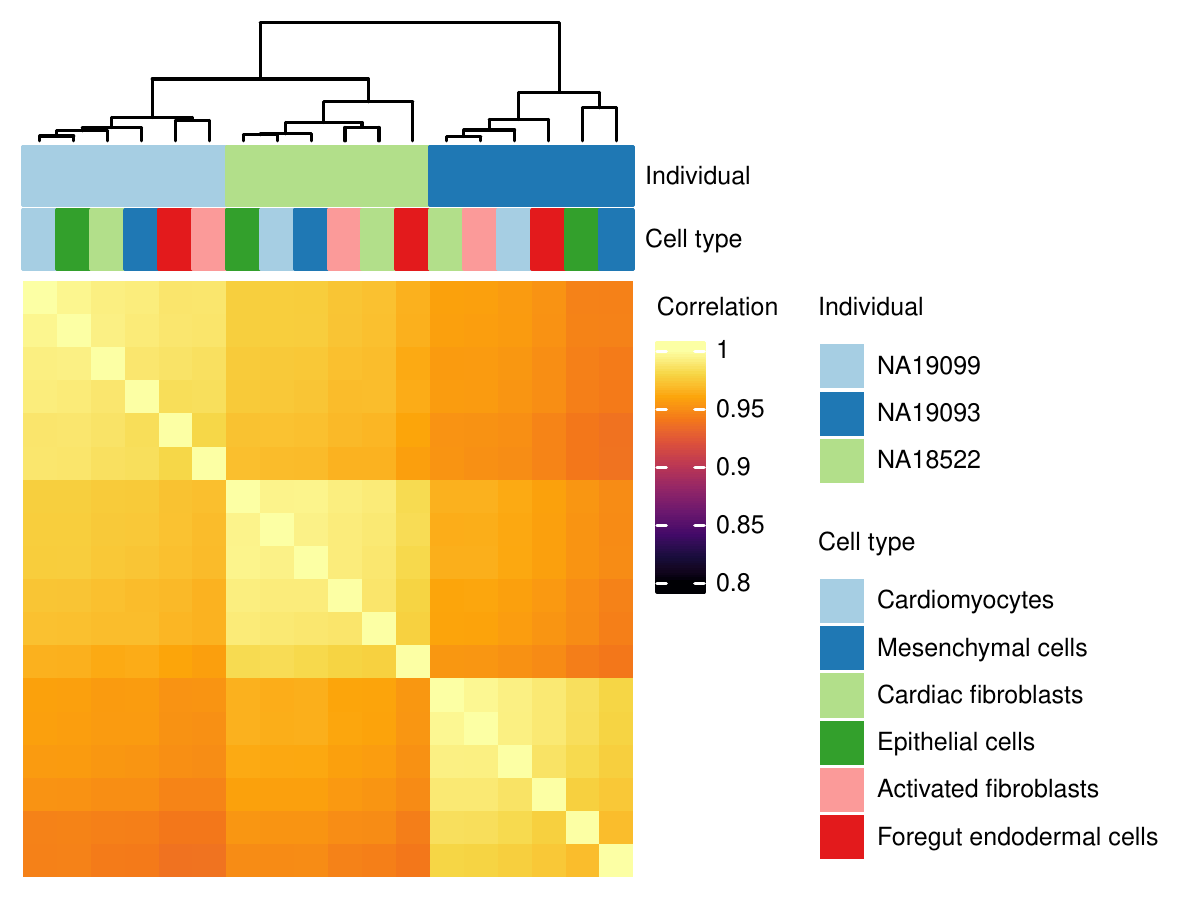

**Figure S14.** Correlation structure of dispersion profiles across samples following random permutation of cell type labels.

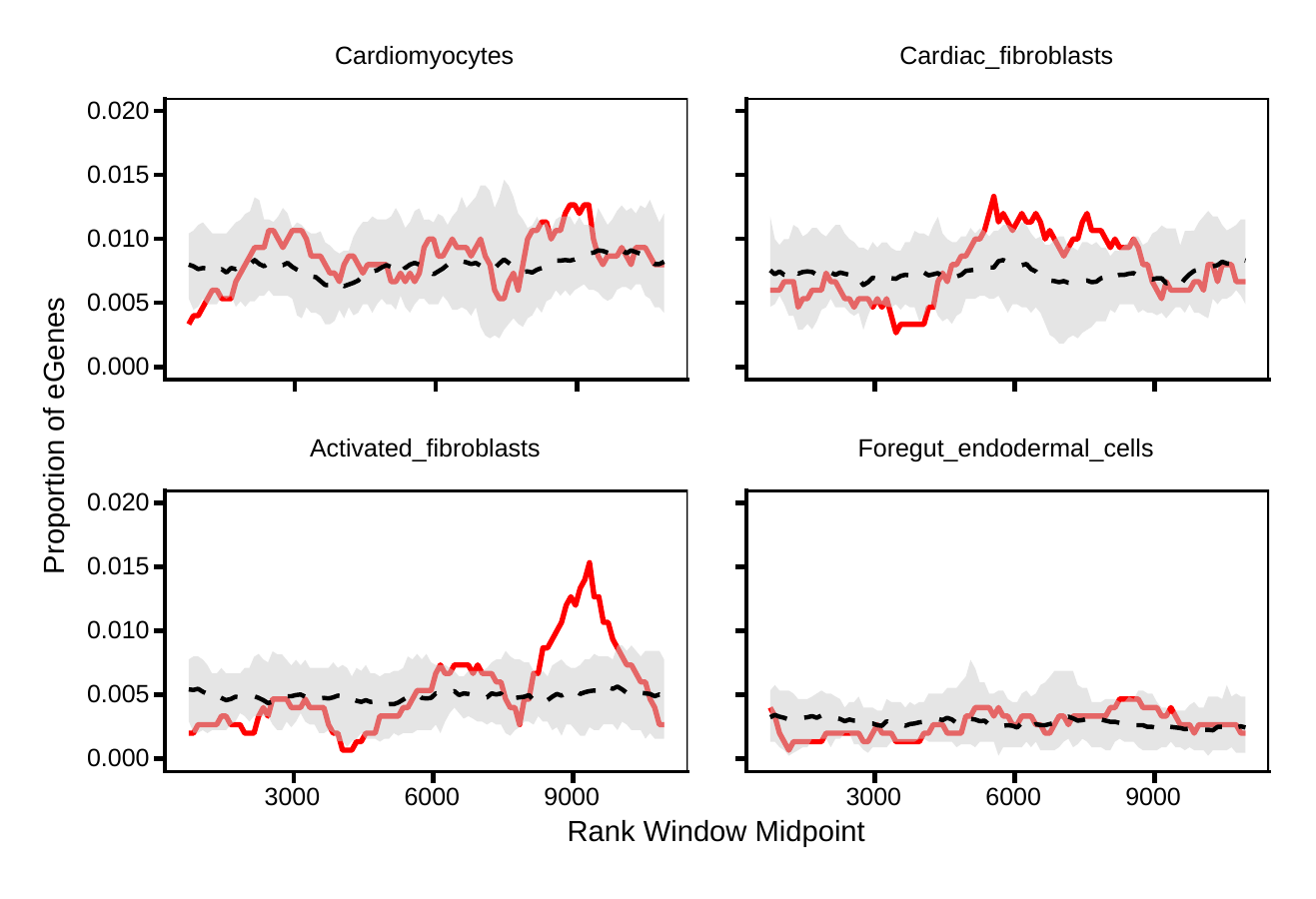

**Figure S15.** Enrichments of eGenes discovered with MatrixEQTL in sliding windows of genes ranked by dispersion in each cell types. The numbers per cell type are small – a couple dozen eGenes.

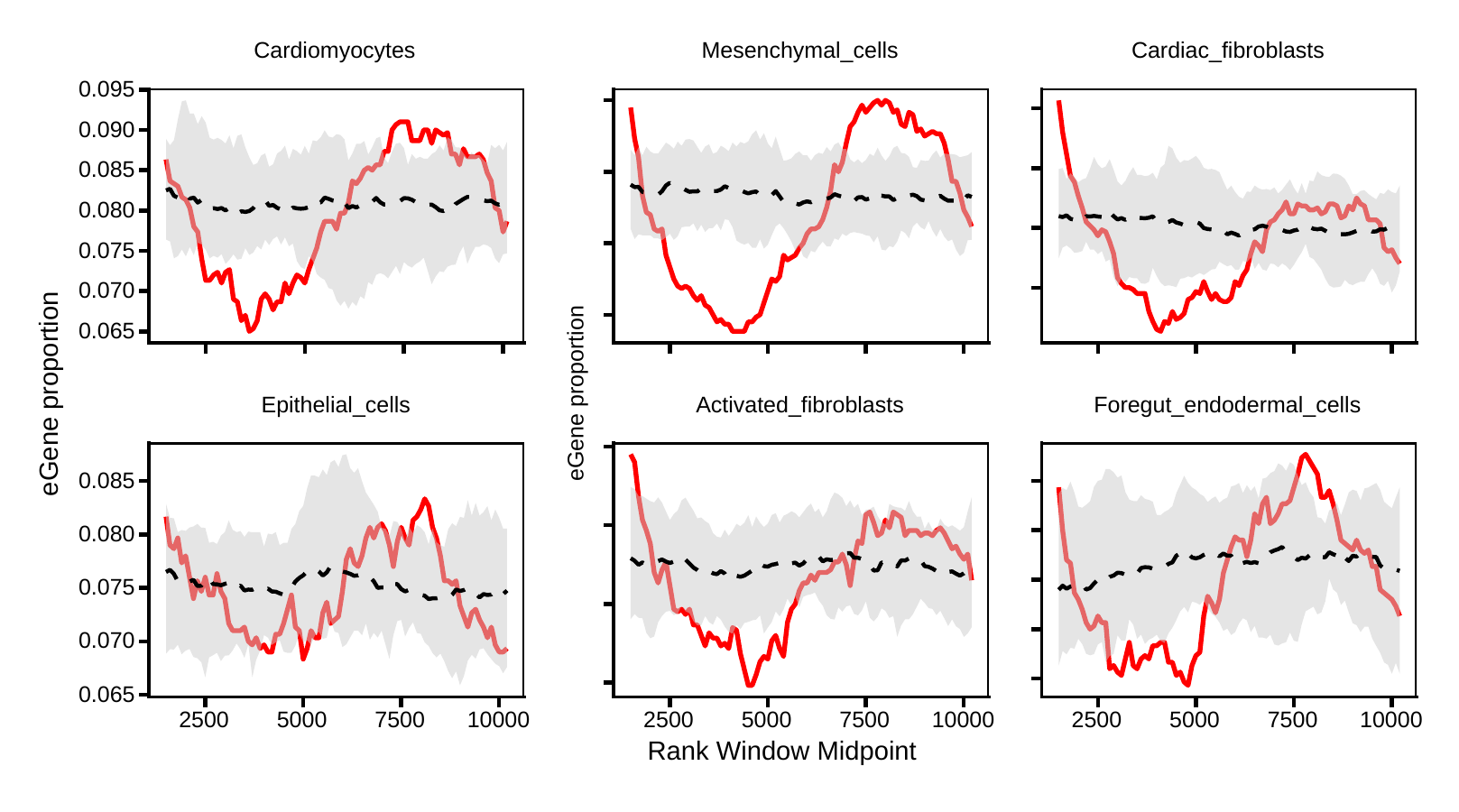

**Figure S16.** Enrichments of eGenes discovered with MASH in sliding windows of genes ranked by dispersion within cell types. MASH is conservative and results in considerable power to detect eQTLs. At the limits, where dispersion is very small or very large, the variance impacts power, which results in the enrichment and depletion of eGenes at the very tails.

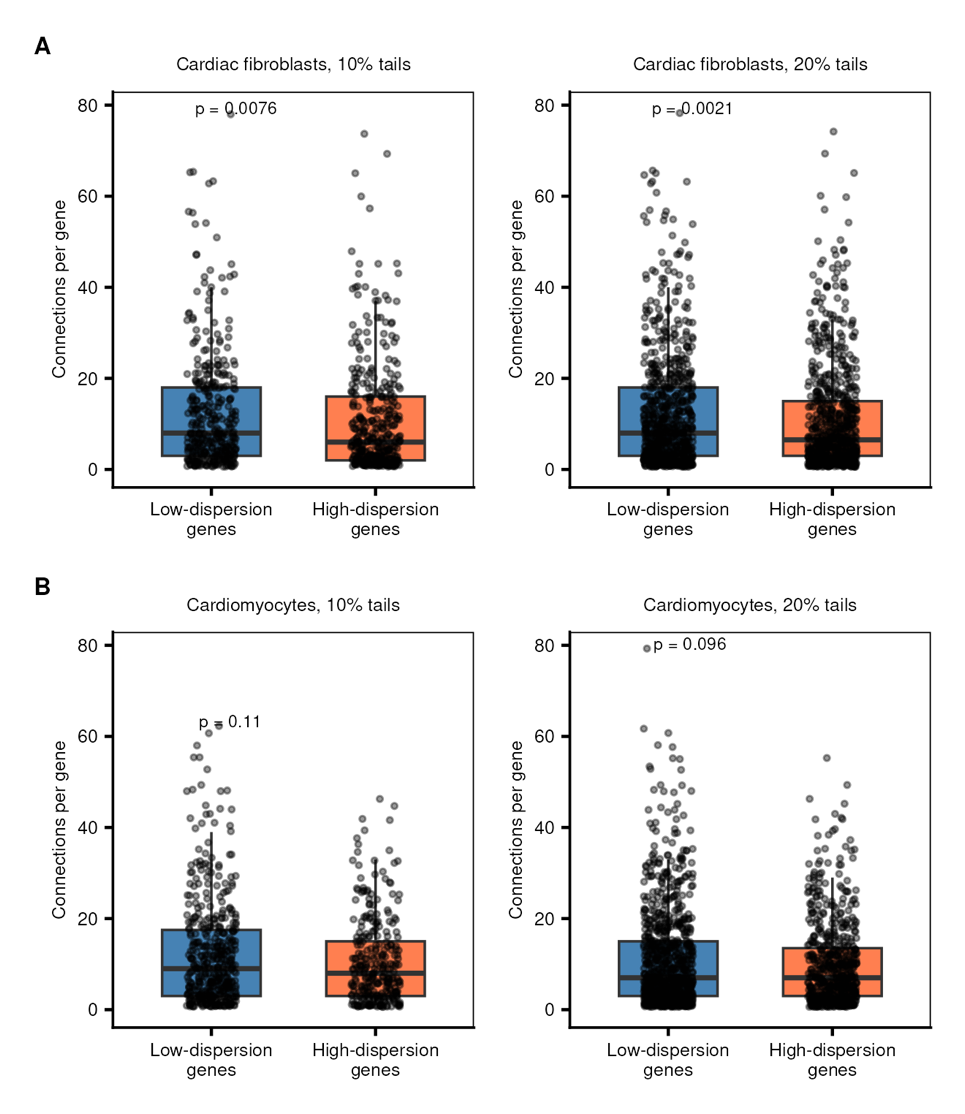

**Figure S17.** Robustness in comparison of per gene connections in cell type-specific co-expression networks. Multiple cutoffs for defining low and high-dispersion genes in cell type-specific dispersion rankings were used to test the robustness of the observation of greater connectivity in low-dispersion genes relative to high-dispersion genes. **A** Comparison of the number of connections per gene in the low-dispersion versus high-dispersion sets in cardiac fibroblasts. **B** Comparison of the number of connections per gene in the low-dispersion versus high-dispersion sets in cardiomyocytes. **Left panels:** Gene sets defined as the 10% tails of the cardiac fibroblast-specific ranking. **Right panels:** Genes sets defined as the 20% tails.

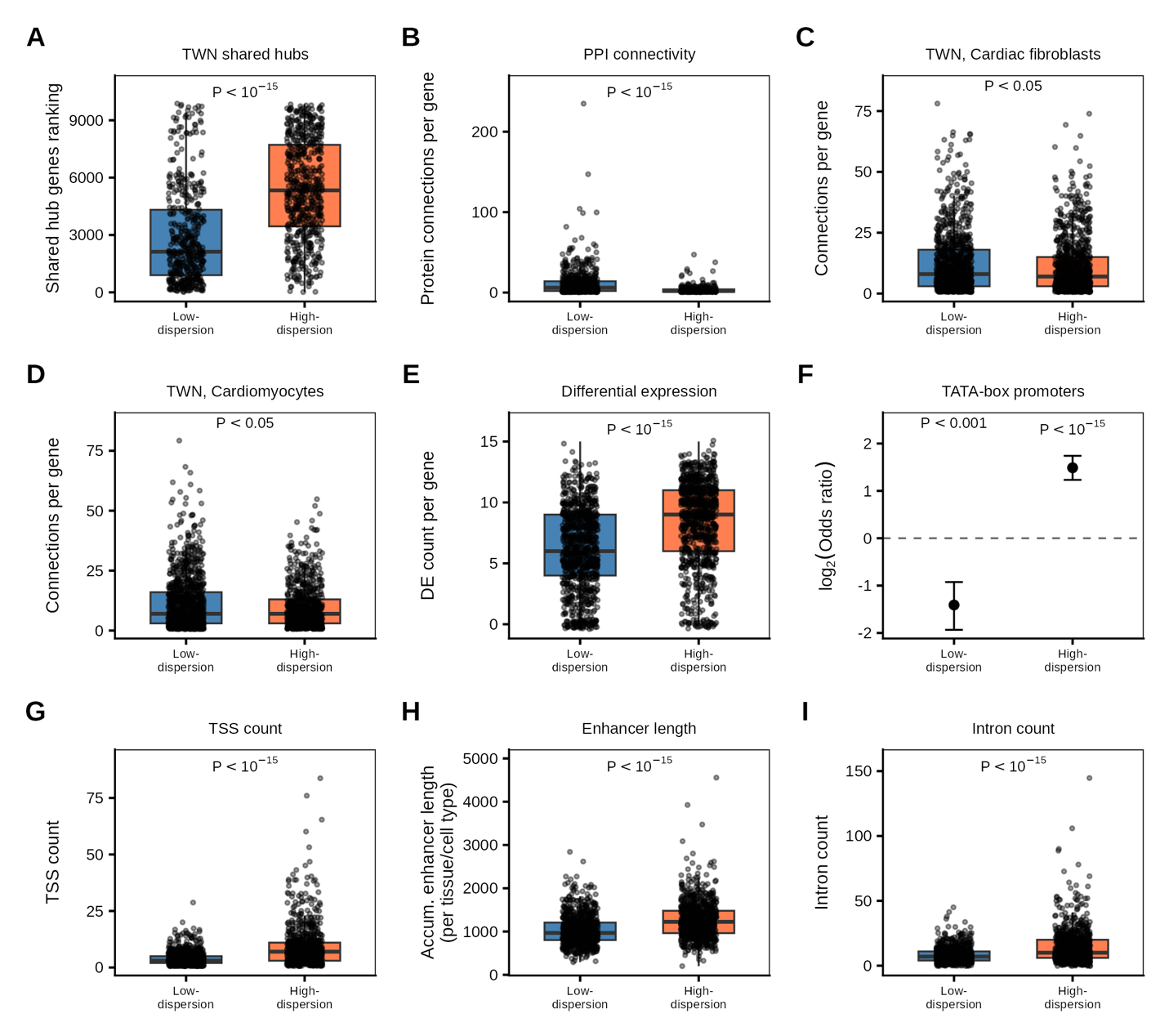

**Figure S18.** Mean-expression-matched replications of the manuscript's Fig 3B-3D & Fig 4, re-tested after equalizing the mean-expression distributions of the consistently-low- and consistently-high-dispersion gene sets. **A** TWN shared-hub rank (Saha et al. 2017 network). **B** Protein-protein interaction (PPI) connectivity. **C** Per-cell-type TWN connectivity for Cardiac fibroblasts and, **D** Cardiomyocytes. **E** Differential-expression (DE) count. **F** TATA-box motif enrichment. **G-I** Comparison of the number of transcription start sites, cumulative enhancer lengths, and number of introns, between low-dispersion and high-dispersion genes, respectively.

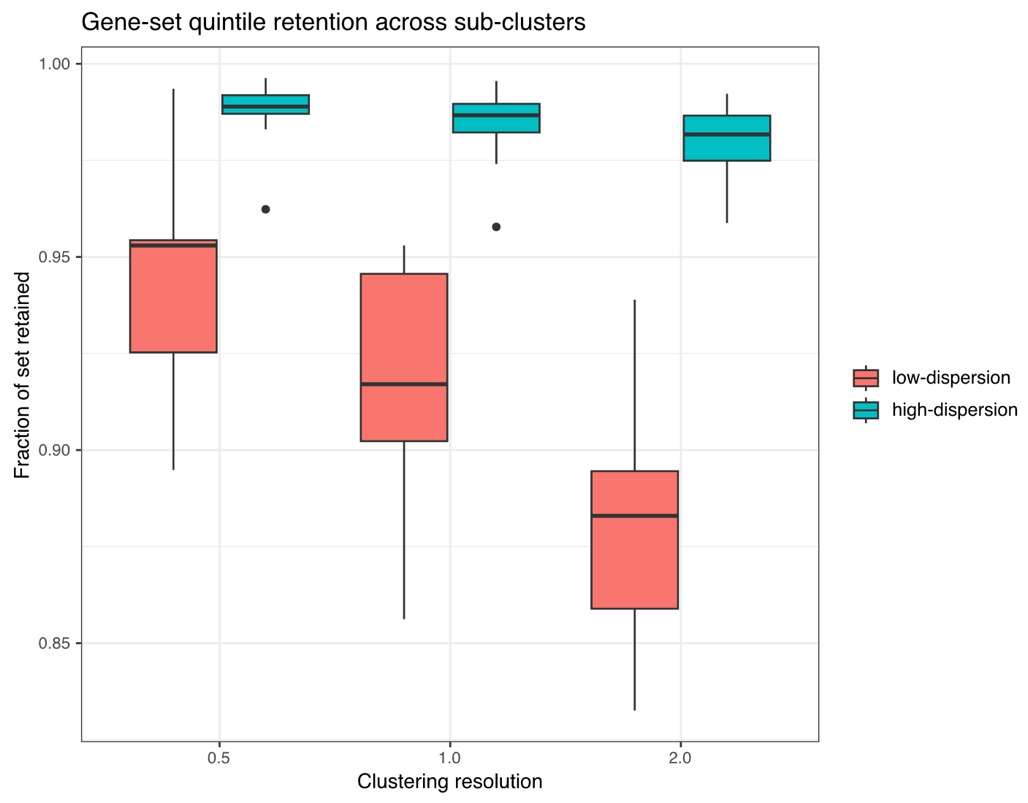

**Figure S19.** For each sub-cluster in Figure 17, the fraction of consistently-low (red) and consistently-high (teal) dispersion gene sets that remain in the same quintile when genes are re-ranked using that sub-cluster's own dispersion estimates, by clustering resolution.

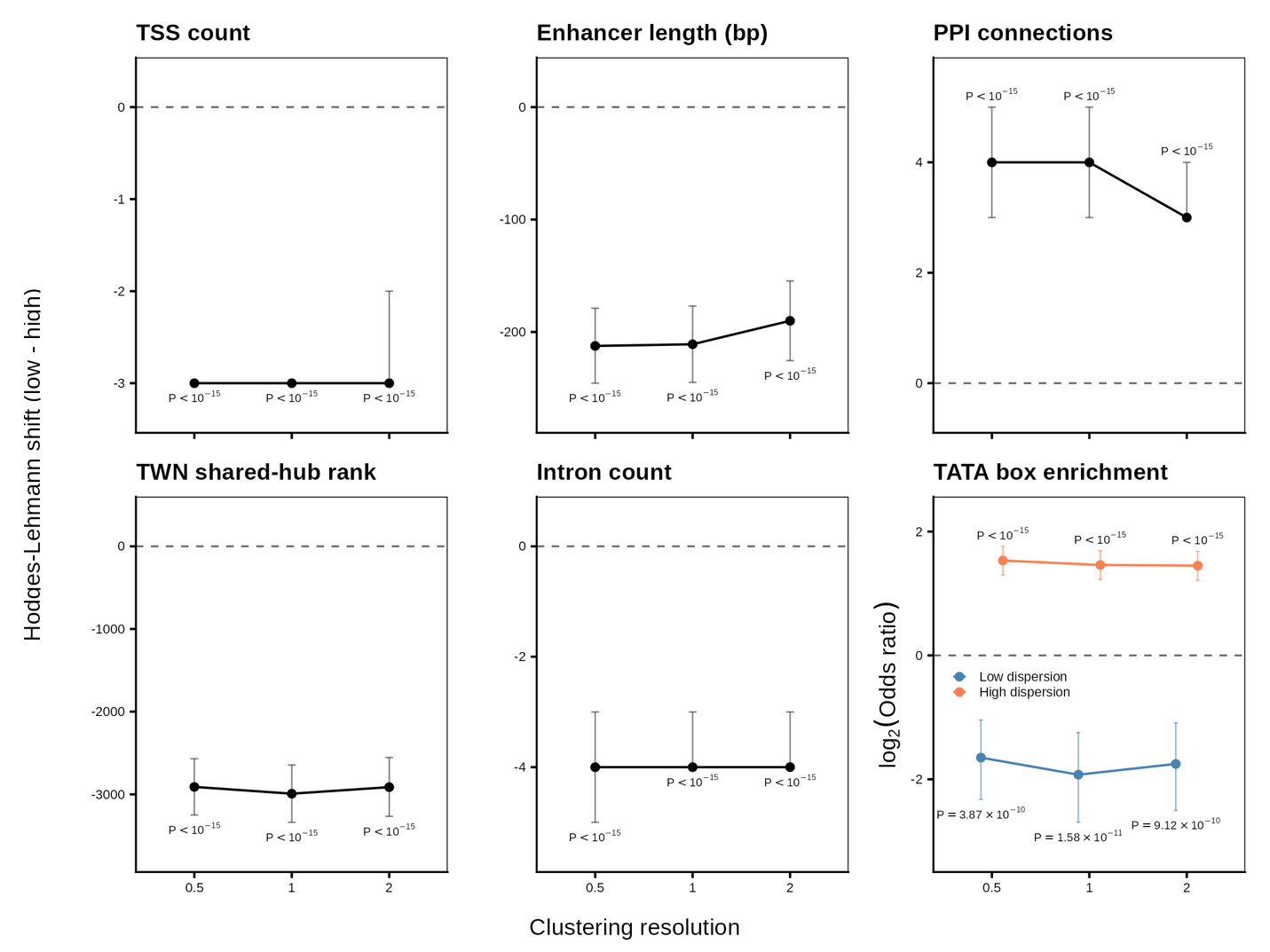

**Figure S20**. Regulatory and network feature comparisons between low- and high-dispersion genes for each clustering resolution. Hodges-Lehmann shift represents the relative value of the high dispersion genes with respect to the corresponding low dispersion genes.

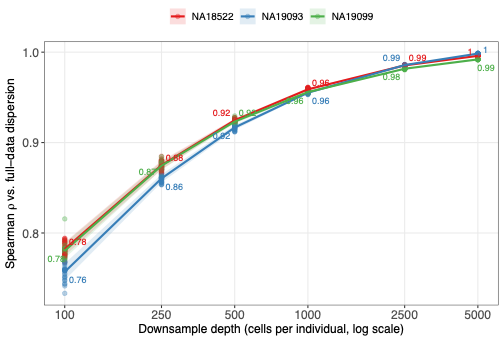

**Figure S21.** Robustness of dispersion estimates across cell count. For the largest cell type, Cardiomyocytes (30,423 cells pooled across 3 individuals — NA19099: 17,064; NA18522: 7,715; NA19093: 5,644), each individual was downsampled to 100, 250, 500, 1,000, 2,500, and 5,000 cells (30 repeated subsample replicates per individual at each depth), and dispersion was re-estimated. Spearman correlation is shown between each replicate's dispersion estimate and that individual's full-data (no downsampling) dispersion.

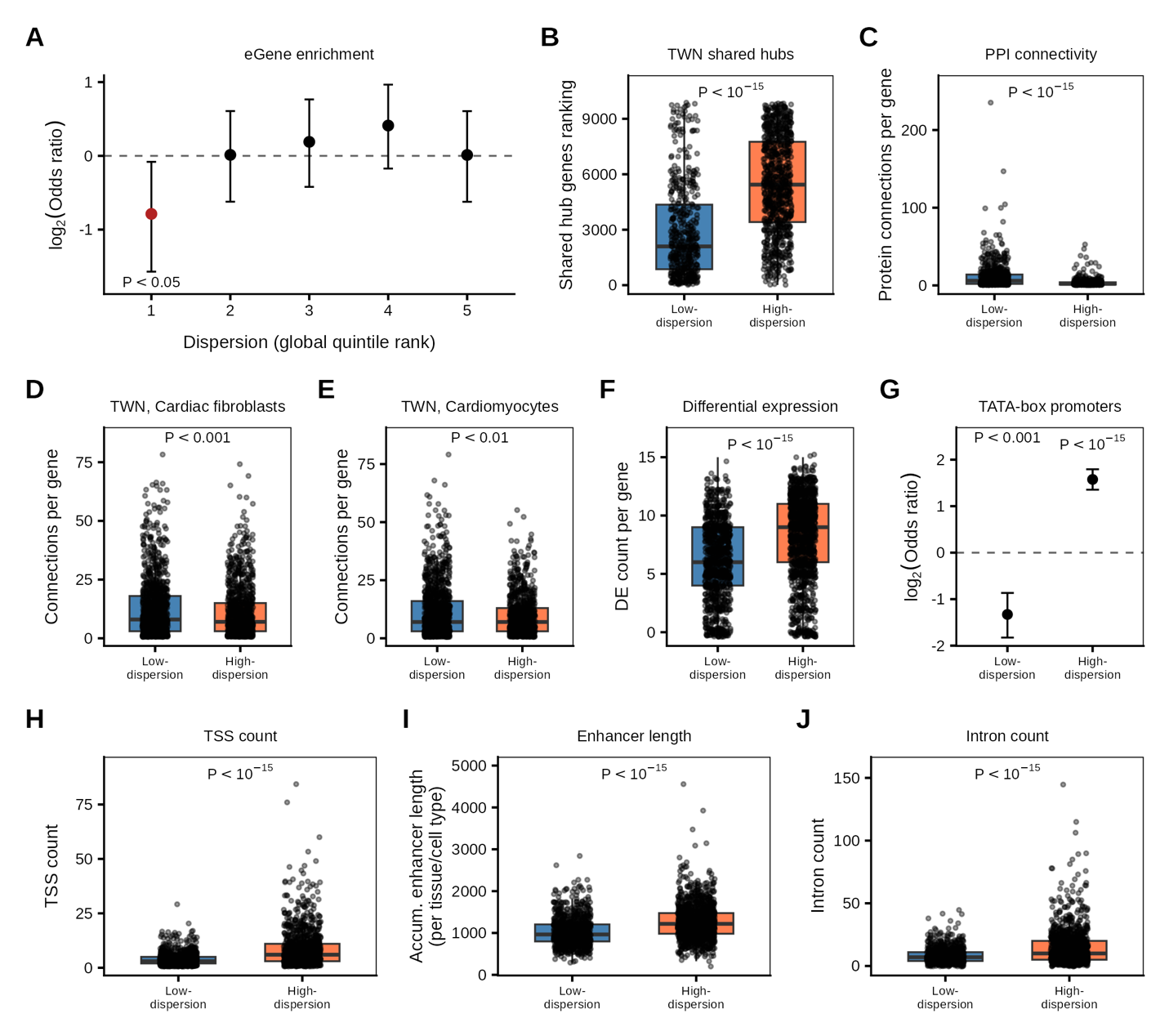

**Figure S22.** Replications of the manuscript's Fig 3 & Fig 4, re-tested after removing known cell-state markers. **A** Enrichment of eGenes in every quintile of the global dispersion ranking. **B** TWN shared-hub rank (Saha et al. 2017 network). **C** Protein-protein interaction (PPI) connectivity. **D** Per-cell-type TWN connectivity for Cardiac fibroblasts and, **E** Cardiomyocytes. **F** Differential-expression (DE) count. **G** TATA-box motif enrichment. **H-I** Comparison of the number of transcription start sites, cumulative enhancer lengths, and number of introns, between low-dispersion and high-dispersion genes, respectively.

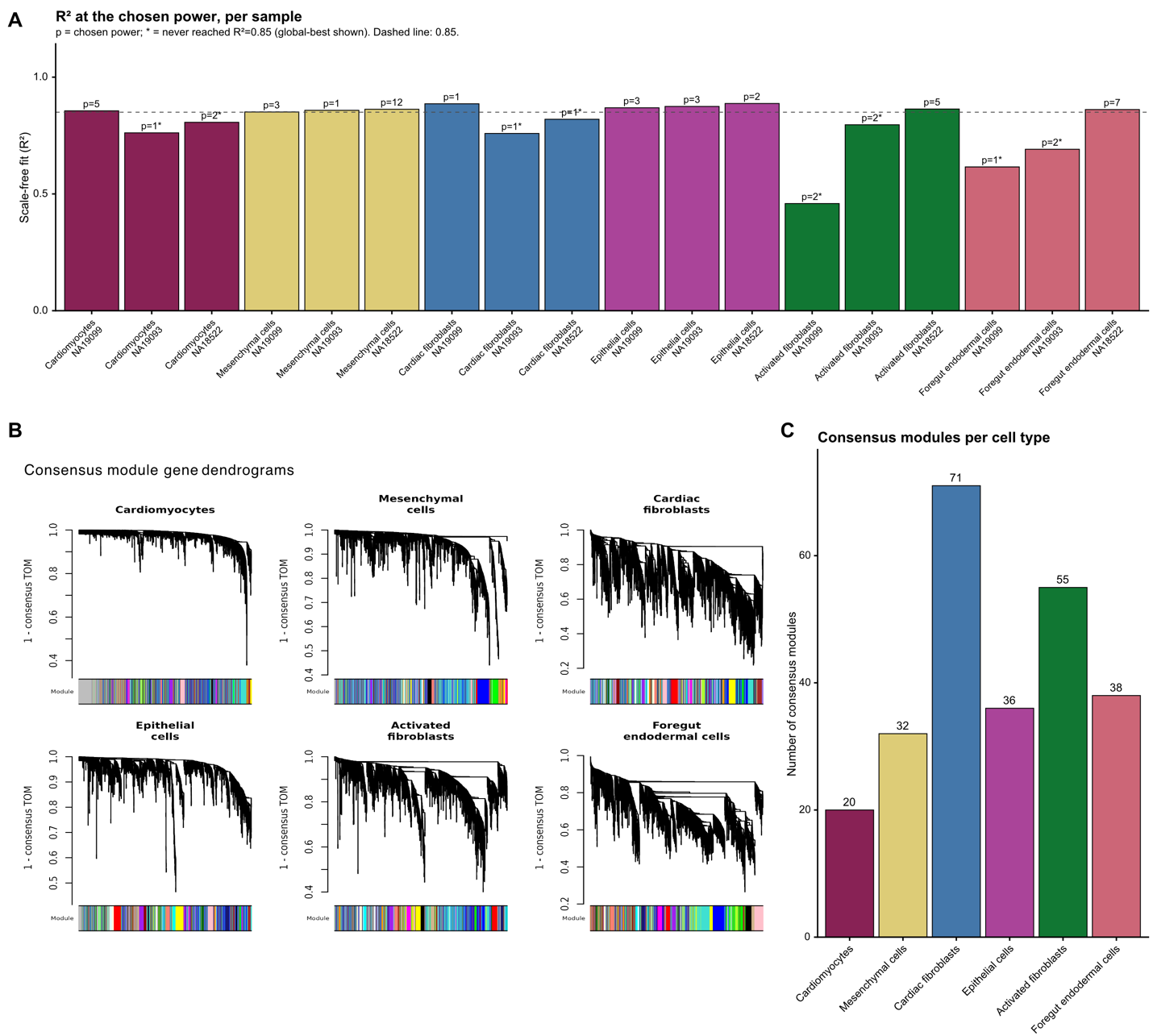

**Figure S23.** Inferred per-sample and consensus modules. **A** Scale-free topology fit (R²) at the chosen soft-thresholding power. Co-expression values below 0.25 were set to zero. The power was chosen as the lowest power that reached R² ≥ 0.85 (dashed line). An asterisk marks samples that never reached R² = 0.85 over the tested powers (1–20). For those, the power giving the best fit in the sweep is shown. Bars are colored by cell type. **B** Gene dendrograms used to call consensus modules in each cell type. Each tree is an average-linkage hierarchical clustering of 1 − consensus topological overlap (TOM). The color bar beneath each tree shows module assignment, and grey marks unassigned genes. **C** Number of consensus modules per cell type.

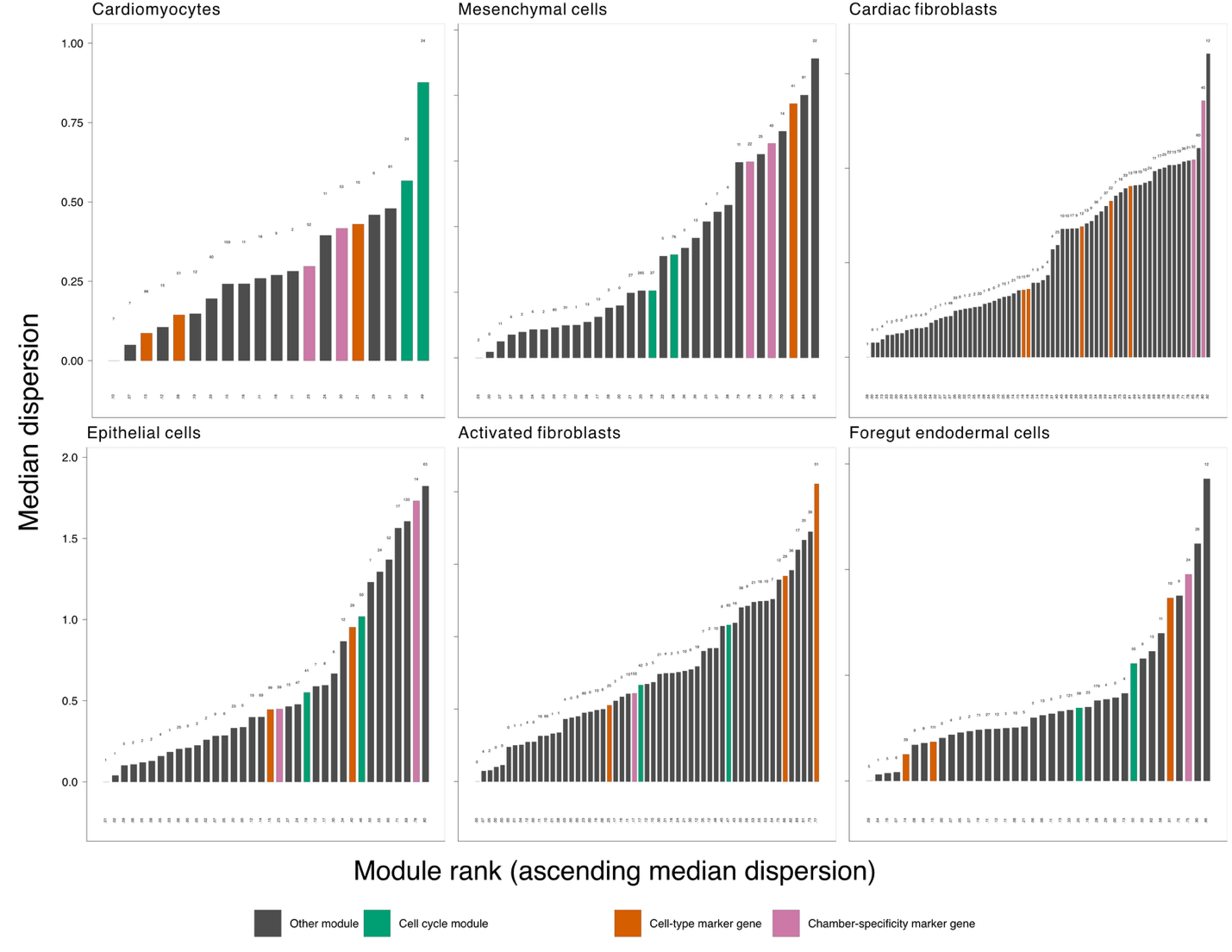

**Figure S24.** Consensus modules ranked by median dispersion of constituent genes. Modules containing cell state markers including cell type-specific, cell cycle, and chamber specificity are colored consistently across cell types. Above each bar is the number of high-dispersion genes with that module. Below each bar is the proportion of that module that is high-dispersion genes.

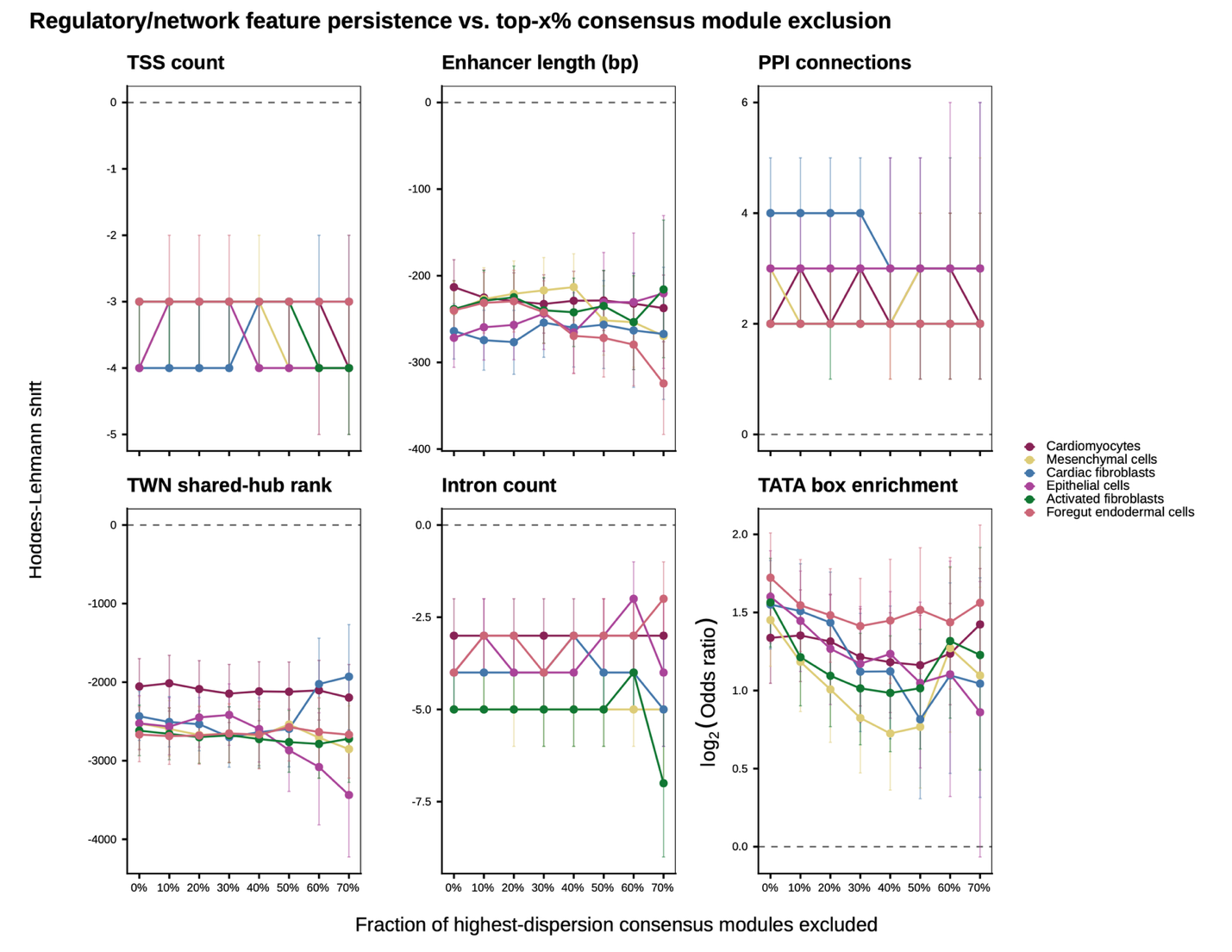

**Figure S25.** Robustness of regulatory and network signatures within the high-dispersion genes. High-dispersion was defined per cell type using the ~5000 gene universe representing each set of consensus modules. Hodges-Lehmann shift represents the relative value of the high dispersion genes with respect to the corresponding low dispersion genes.

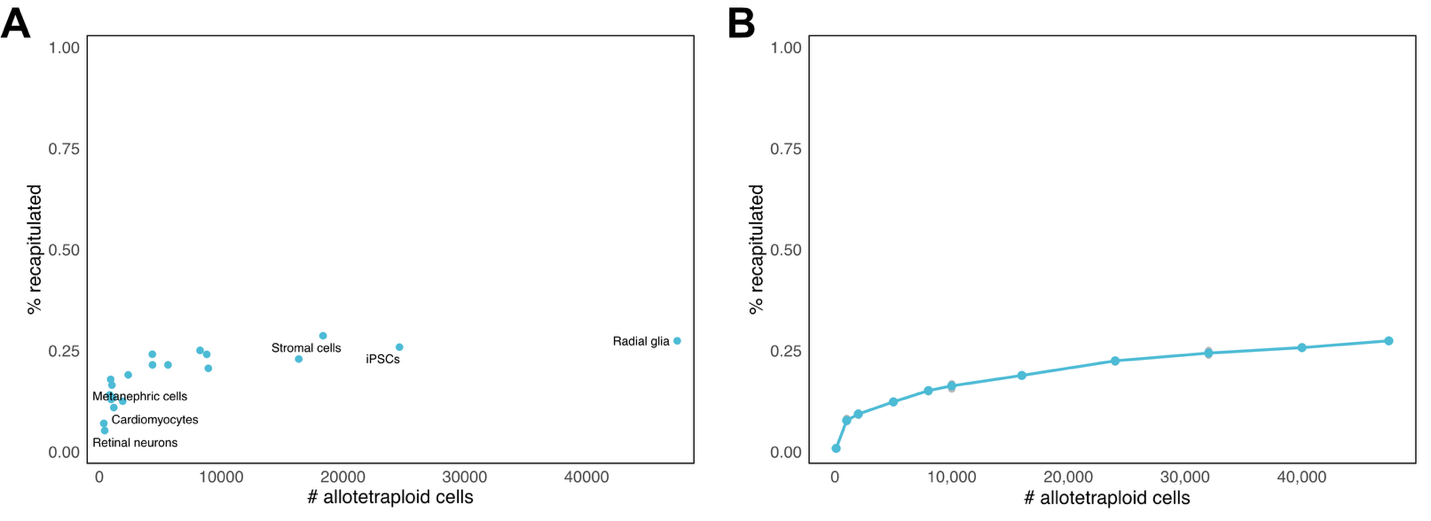
**Figure S26.** Recapitulation rate of diploid DD genes (% recapitulated with matching effect-size sign in the allotetraploid) as a function of allotetraploid cell number. **A** Each dot represents the recapitulation rate for cell types in the interspecies data. **B** For the largest cell type, radial glia (47,437 cells), the allotetraploid cell counts generated from downsampling between 100 to 40,000 cells with 10 independent random-subsample replicates. SD and 95% CIs provided in Table S7.

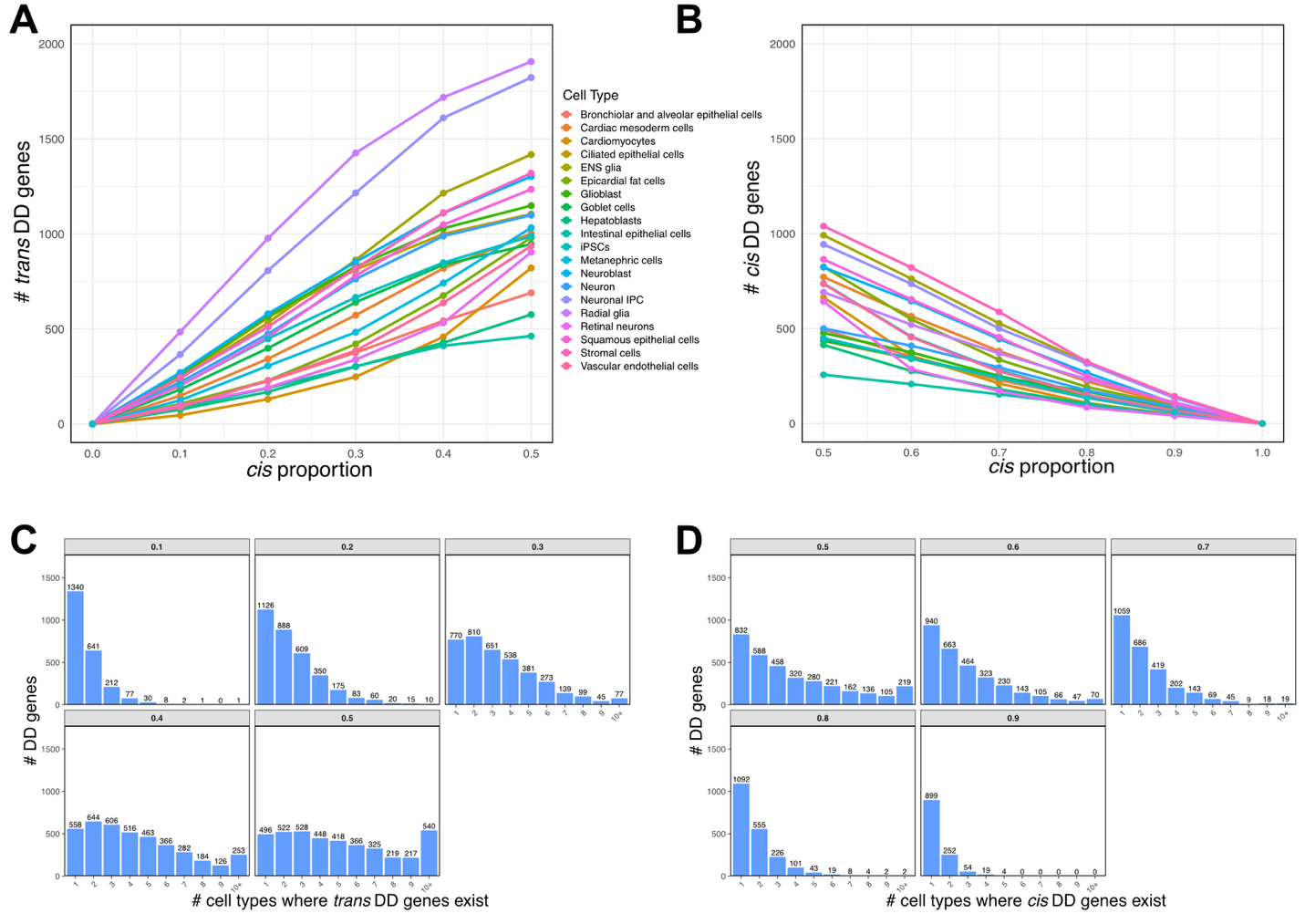

**Figure S27.** Number of *trans* and *cis* DD genes based on *cis* proportion cutoffs. **A-B** Among identified significant DD genes within a cell type (LFSR < 0.05; recapitulated in both diploid and allotetraploid cells), we classify *trans* or *cis* DD genes based on their estimated *cis* proportion. We vary the *cis* proportion threshold, defining trans DD genes as those with *cis* proportion below the cutoff and cis DD genes as those with *cis* proportion above the cutoff. **C-D** For each *cis* proportion cutoff, we quantify the number of DD genes that exist in multiple cell types, separately for trans and cis DD genes.

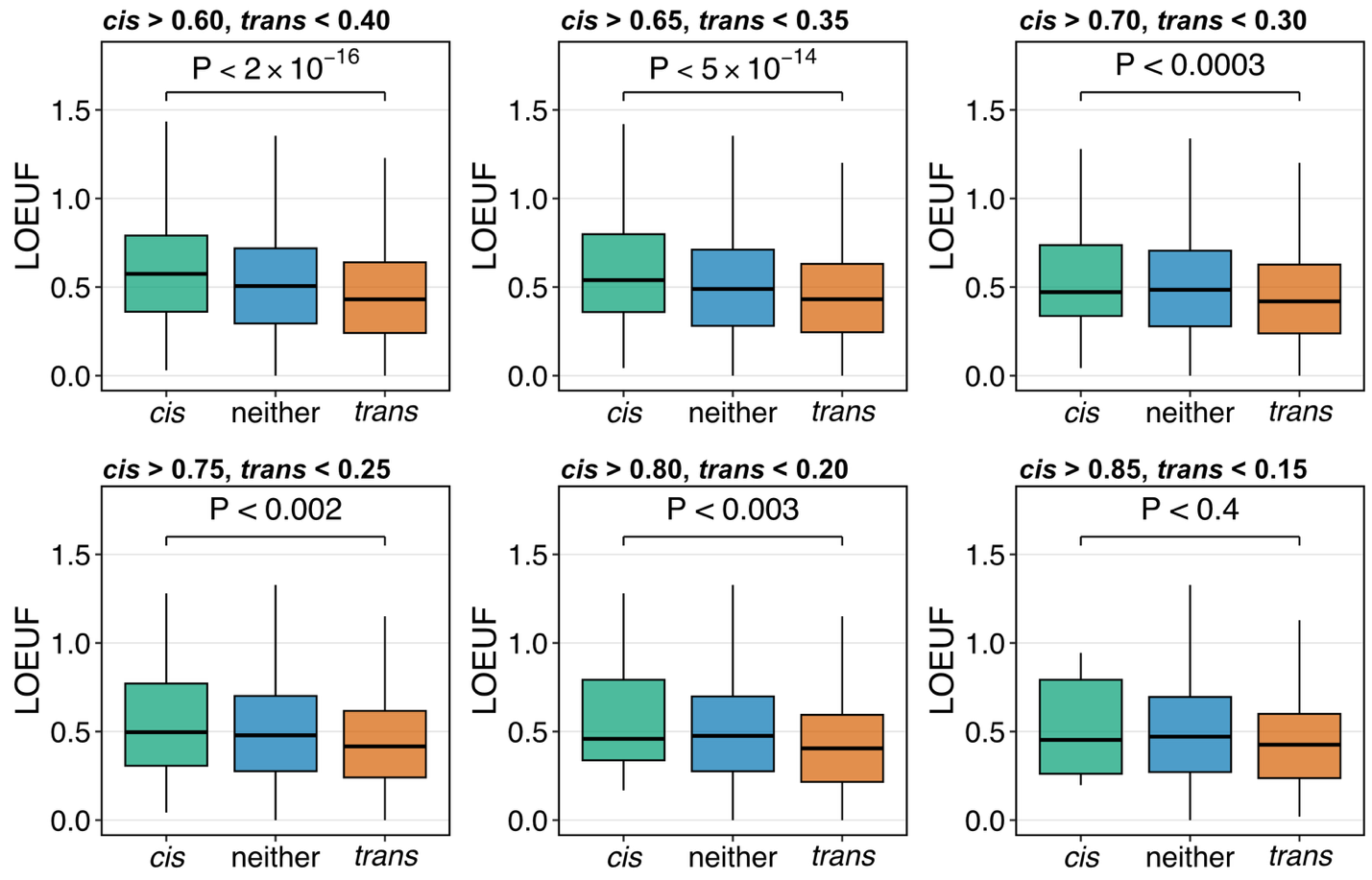

**Figure S28.** LOEUF scores based on different mean *cis* proportion cutoffs of cell types DD to define *cis* and *trans* DD genes, across cell types.

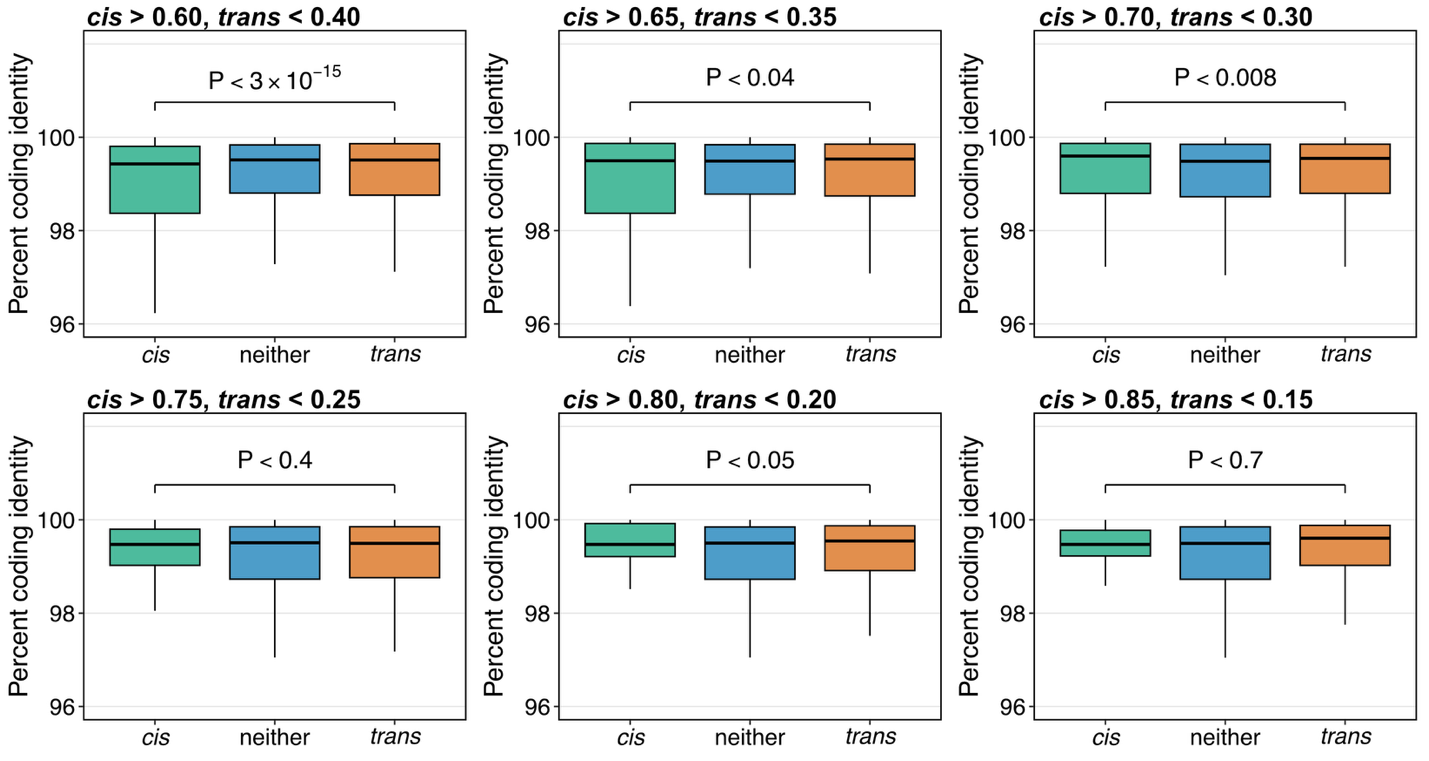
**Figure S29.** Percent coding identity (human-chimpanzee divergence) based on different mean *cis* proportion cutoffs of cell types DD to define *cis* and *trans* DD genes, across cell types.

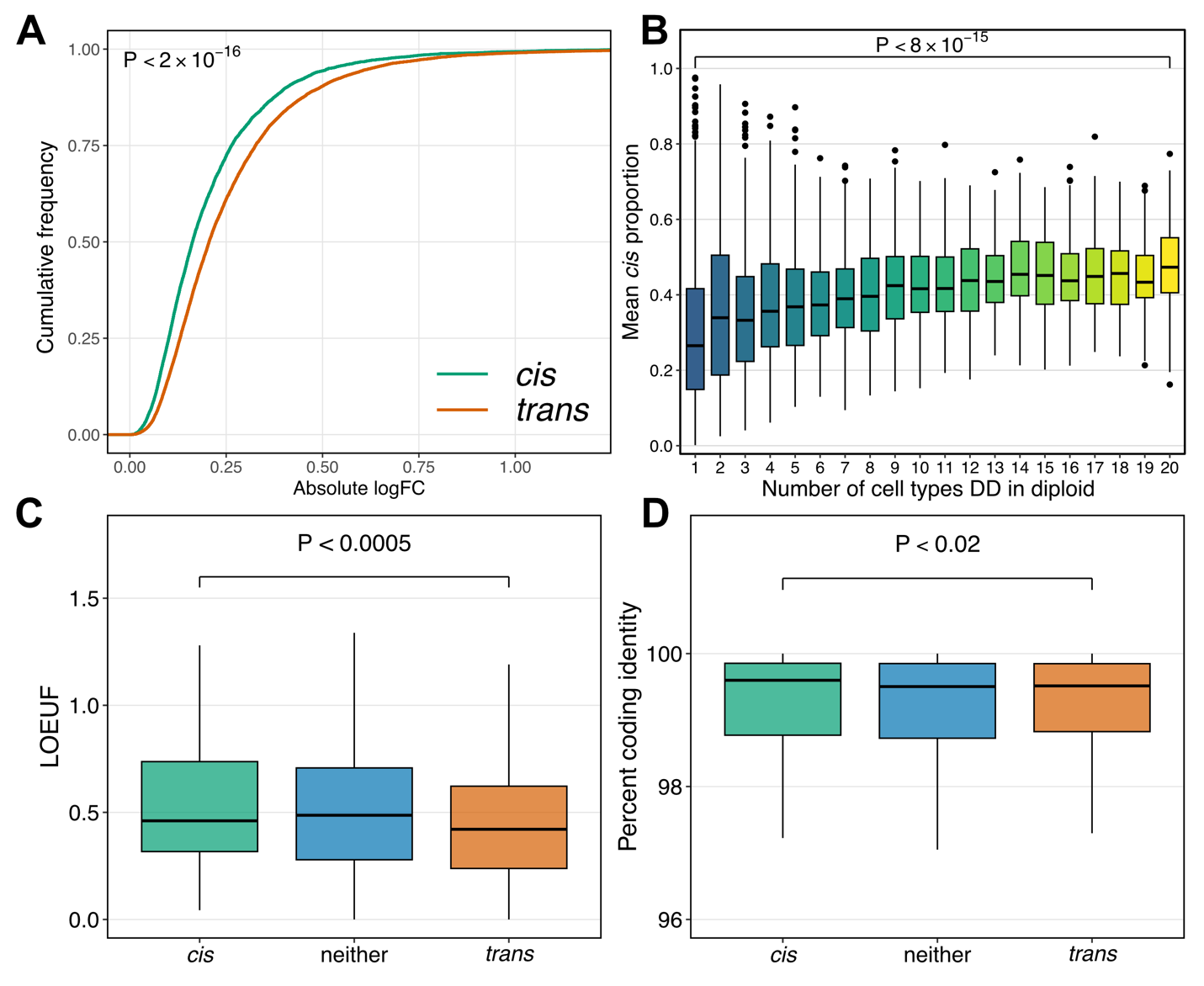

**Figure S30. Patterns of *cis* and *trans* DD effects using a stricter log-mean cutoff of -7 (exclude genes with log-mean > -7) A.** Empirical cumulative distributions of effect size for all cell types in diploid cells for tests with high cis proportion (> 0.70) and low cis proportion (< 0.30). **B.** Boxplot distribution of cis proportion for genes DD in the diploid cells for an increasing number of cell types. **C-D.** Boxplots showing the distribution of LOEUF and human-chimpanzee divergence (percent coding identity) respectively, for genes whose interspecies DD can be accounted for by a high, moderate, or low cis effect.

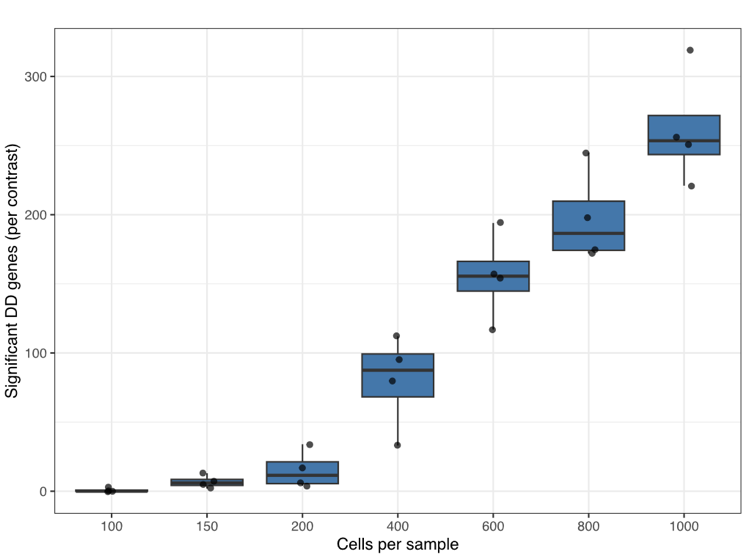

**Figure S31.** Significant differential-dispersion (DD) genes detected in McIntire et al’s cHDC data (batches B2+B6; DD analysis done in each batch separately) across downsampled cell-count thresholds (100–1000 cells per individual per cell type). Each box summarizes the 4 shared cell-type contrasts (adj. P < 0.05, BH-corrected); points show the per-contrast counts.

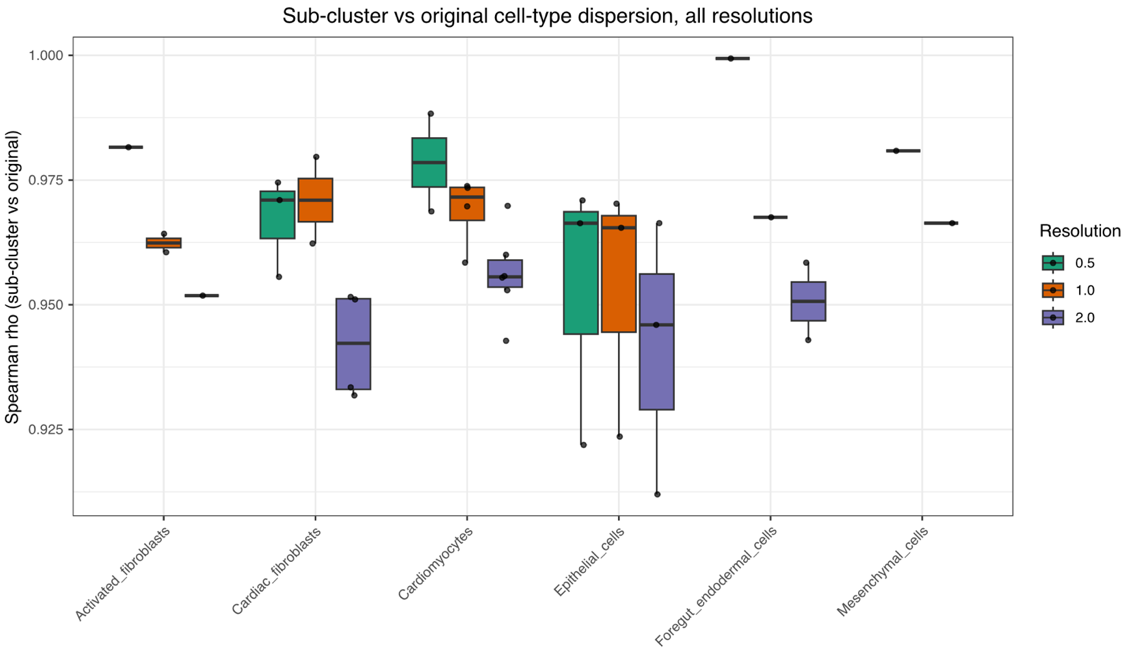

**Figure S32.** Each of the manuscript's 6 published cell types was sub-clustered at 3 Seurat resolutions (0.5/1.0/2.0; sub-clusters kept only if ≥ 2 individuals clear a 700-cell threshold), dispersion re-estimated per sub-cluster with Memento, and Spearman-correlated back against that cell type's original, published cell-type-level dispersion. Sub-cluster-vs-original rho is consistently high (0.91-1.00) across every cell type and resolution.

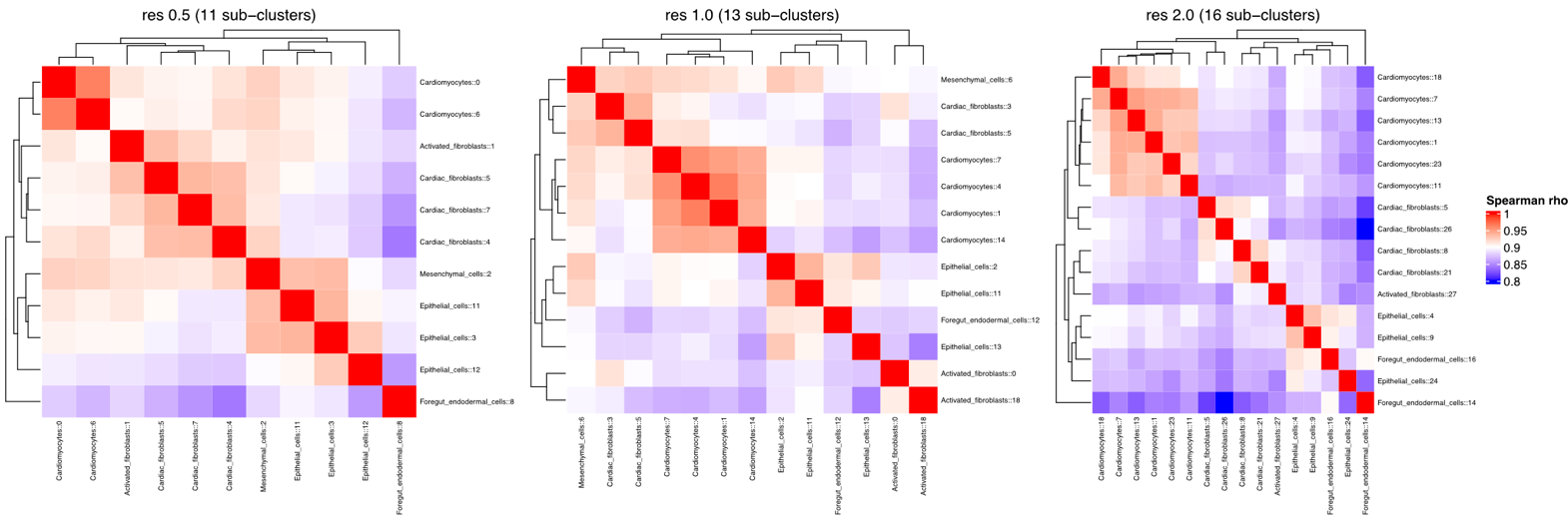

**Figure S33**. Spearman correlation heatmaps of pairwise sub-cluster dispersion estimates (all sub-clusters across all 6 cell types, one panel per clustering resolution: 0.5/1.0/2.0), hierarchically clustered on both axes.

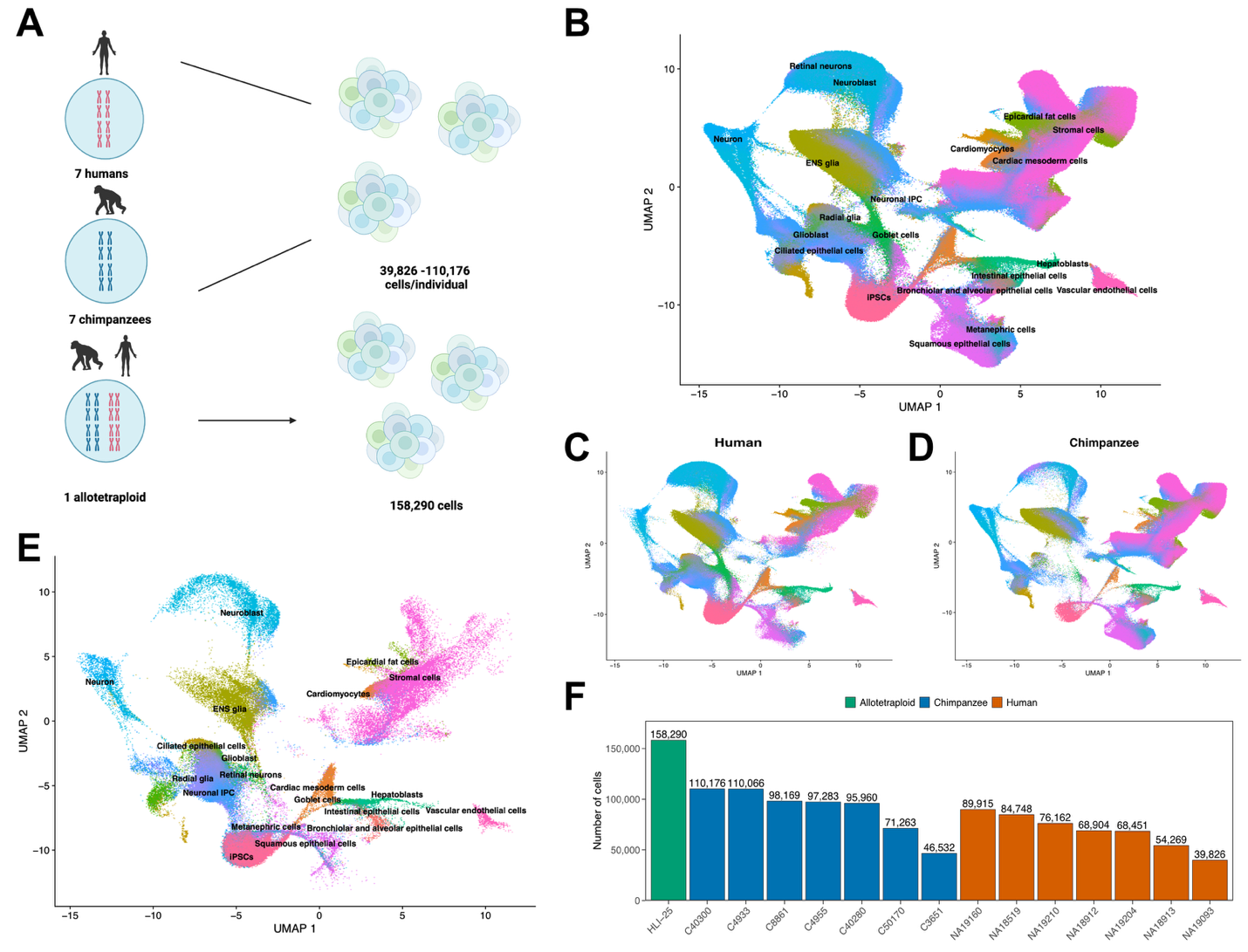

**Figure S34.** Interspecies study design and dataset. **A** Study design: scRNA-seq data were generated from HDCs derived from iPSC lines from 7 human individuals, 7 chimpanzee individuals, and 1 allotetraploid line. **B** UMAP projection of all diploid human and diploid chimpanzee single cells after quality control filtering and reference-based annotation. **C-D** UMAP projections restricted to diploid human (**C**) or diploid chimpanzee (**D**) cells, respectively. **E** UMAP of allotetraploid cells, annotated using the human reference, projected onto the embedding shown in (**B**). **F** Number of single cells recovered per individual after quality control and preprocessing.

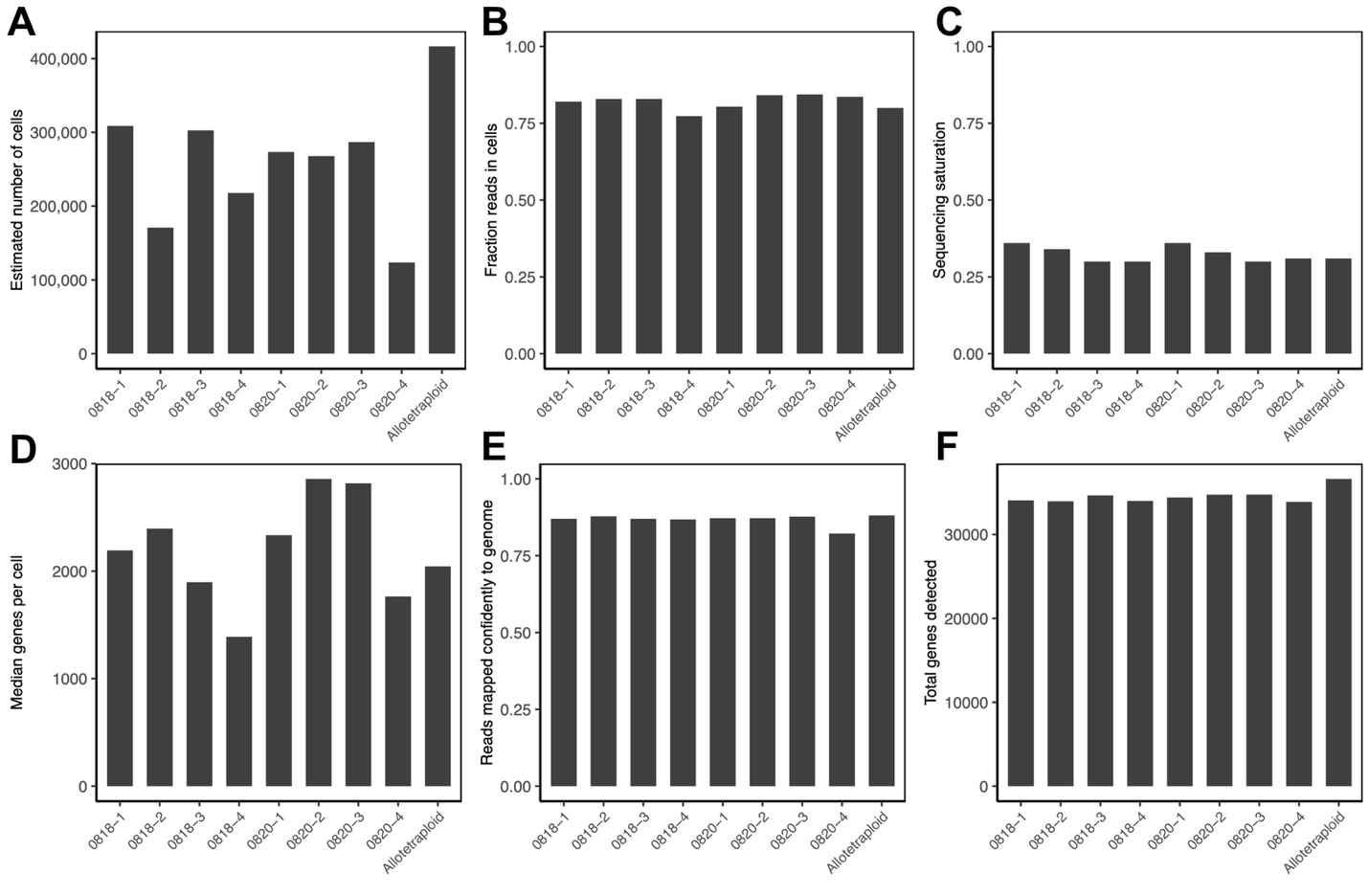

**Figure S35.** Quality control metrics for interspecies dataset. Each bar is one ScaleBio pool (9 pools: 4 chimpanzee, 4 human, 1 allotetraploid). Pools Sample-0818-1/2 and Sample-0820-1/2 are diploid chimpanzee; Sample-0818-3/4 and Sample-0820-3/4 are diploid human. Allotetraploid values shown are for humanized data. **A** Estimated number of cells. **B** Proportion of reads in cell per pool. **C** Sequencing saturation. **D** Median number of detected genes per cell per pool. **E** Proportion of genes mapped confidently to the genome per pool. **F** Total number of genes detected per pool.

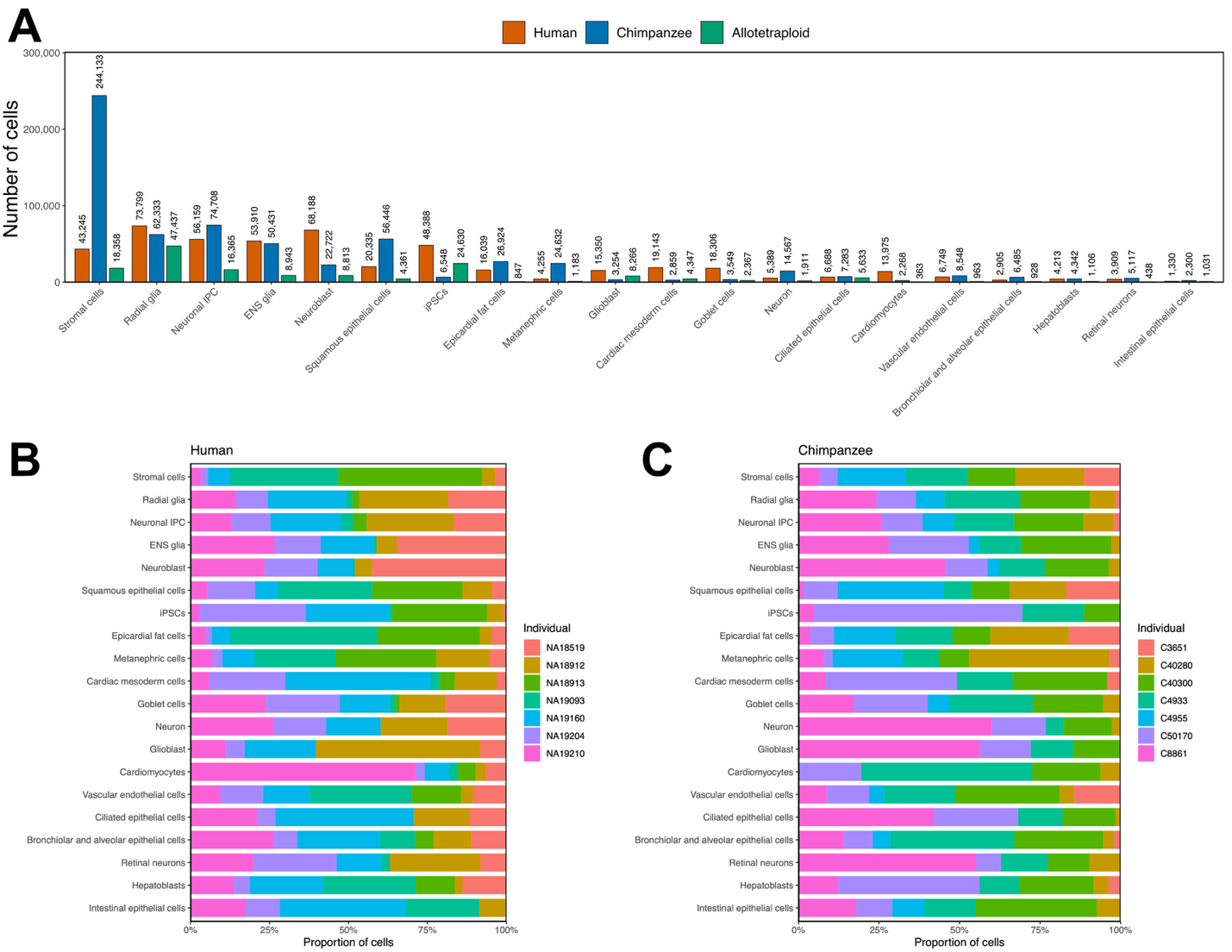

**Figure S36.** Interspecies dataset by cell type. **A** The number of diploid human, diploid chimpanzee, and allotetraploid single cells by cell type, respectively. **B** The proportion of diploid human cells by individual within each cell type. **C** The proportion of diploid chimpanzee cells by individual within each cell type.

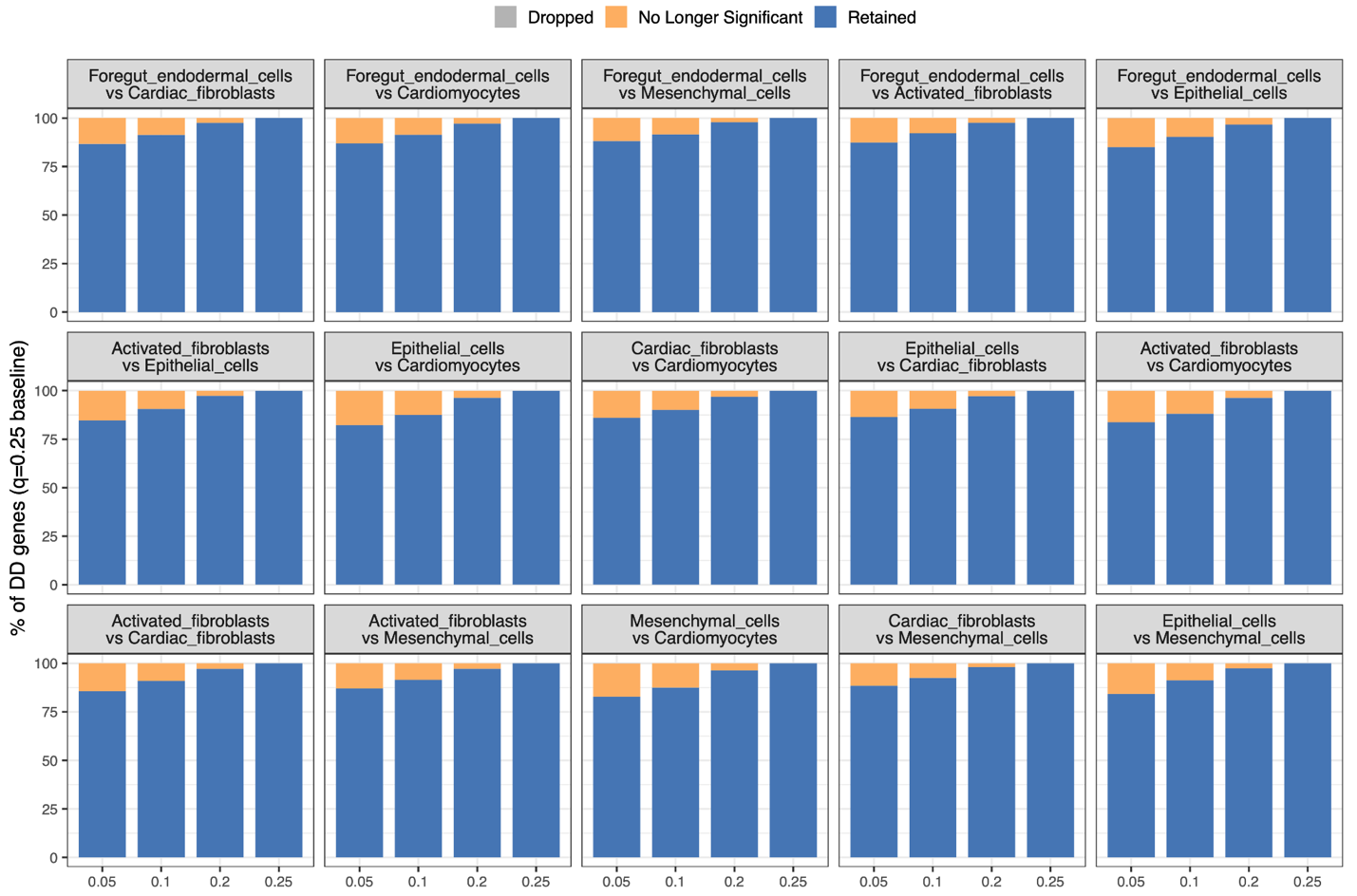

**Figure S37**. For each of the 15 pairwise cell-type DD contrasts, DD genes at a baseline *q*=0.25 are defined as retained, no longer significant, or lost from the gene universe as *q* is reduced to 0.05: Retention stays above ~85% even at the lowest *q*, and losses are almost entirely to non-significance rather than genes dropping from the universe.

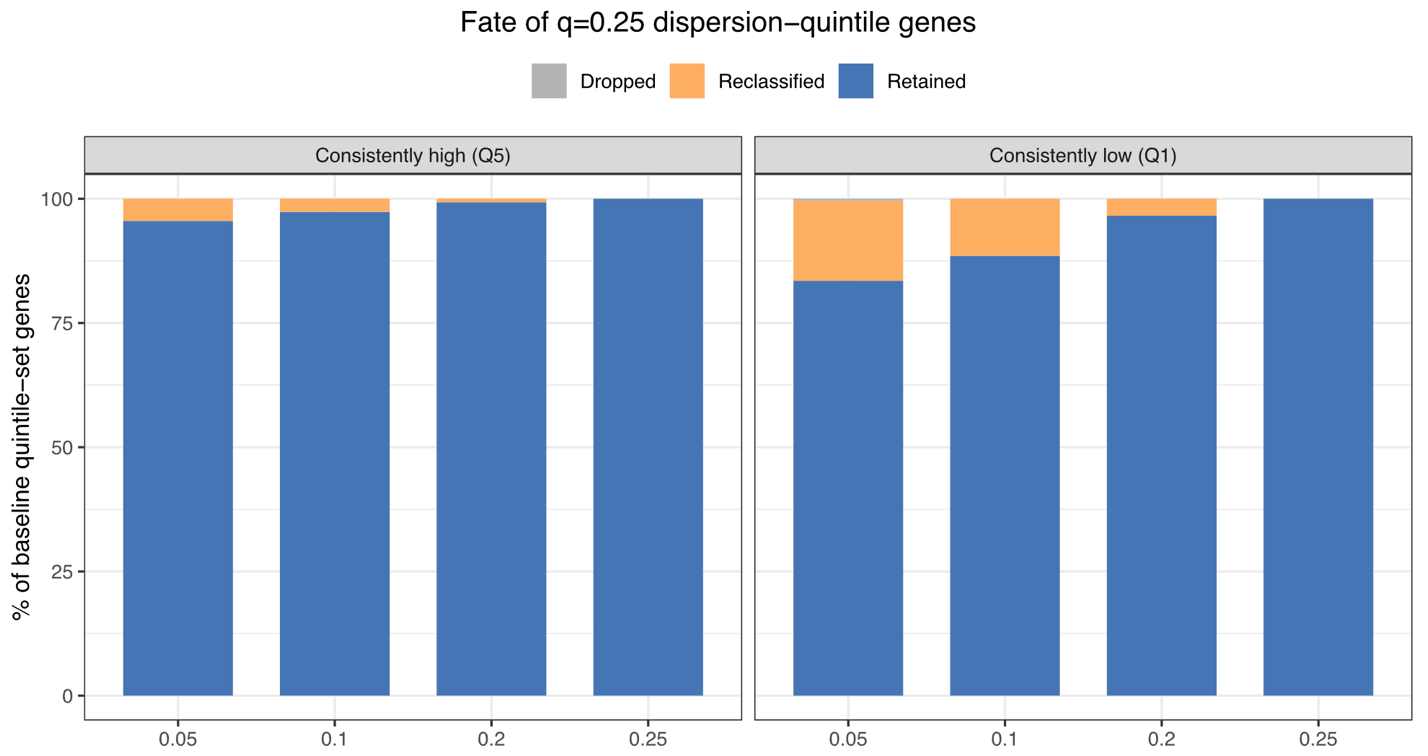

**Figure S38***.* Consistently low and high-dispersion gene sets at the baseline *q*=0.25 are defined as retained, reclassified or dropped from the gene universe as q decreases to 0.05. Both sets of genes are highly stable (>80% retained across *q* values).

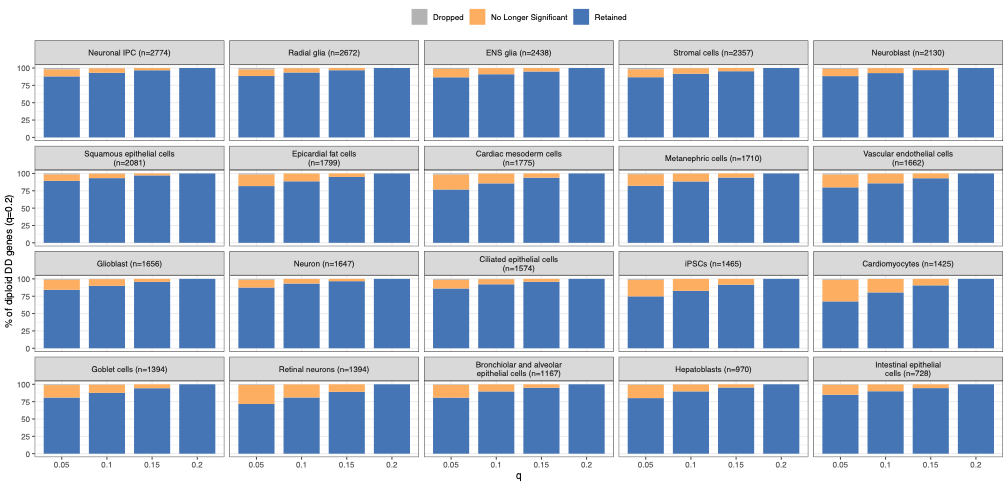

**Figure S39.** Sensitivity of *q* threshold on diploid DD genes in interspecies data. For each lower *q* value tested (*q* ∈ {0.05, 0.1, 0.15}), the percentage of diploid DD genes (LFSR < 0.05) identified at the baseline *q* = 0.2 that are retained, lose significance, or are dropped/unmeasured is shown. Blue indicates retained DD genes, yellow indicates loss of significance, and grey indicates genes dropped or not measured. Cell types are ordered by the number of significant DD genes at baseline.

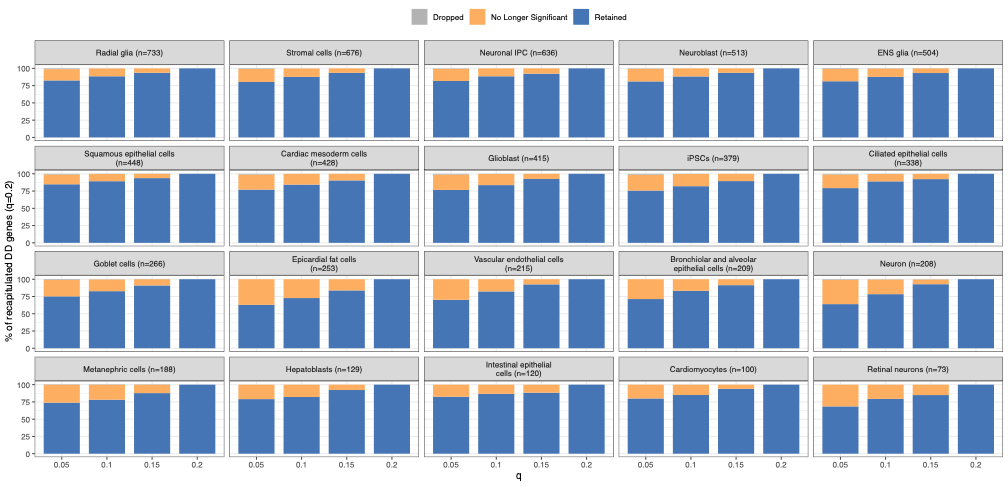

**Figure S40.** Sensitivity of *q* threshold on recapitulated DD genes in interspecies data. For each lower *q* value tested (*q* ∈ {0.05, 0.1, 0.15}), the percentage of recapitulated DD genes (LFSR < 0.05 in diploid and allotetraploid settings with matching effect size sign) identified at the baseline *q* = 0.2 that are retained, lose significance, or are dropped/unmeasured is shown. Blue indicates retained DD genes, yellow indicates loss of significance, and grey indicates genes dropped or not measured. Cell types are ordered by the number of significant DD genes at baseline.

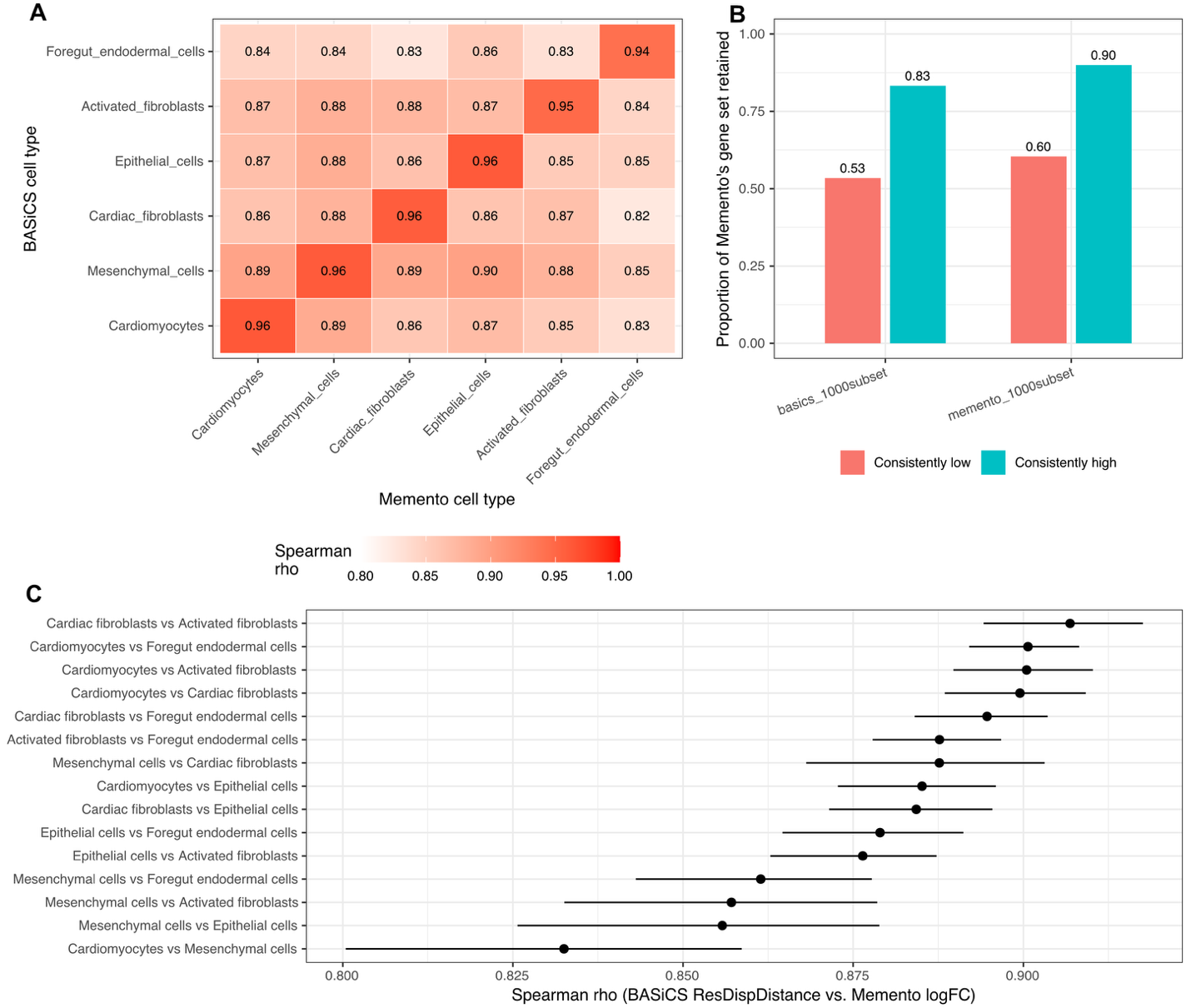

**Figure S41**. Cross-method validation of Memento's dispersion estimates. **A** Spearman correlation between BASiCS's epsilon (residual over-dispersion) and Memento's residual variance, cell-count-matched to 1,000 cells/individual. The matched cell type is each BASiCS estimate's best Memento correlate. **B** Proportion of Memento's consistently low- (red) and high-dispersion (teal) gene sets retained as the same class under each alternative method/condition. **C** Correlation in DD effect sizes between BASiCS and Memento across all 15 pairwise cell-type comparisons. For each pair of cell types, points show the Spearman correlation between BASiCS's difference in residual over-dispersion (epsilon) and Memento's DD log fold change, computed for gene called significant from Memento. Horizontal bars show bootstrap 95% confidence intervals (2,000 resamples of genes with replacement).Figure S27

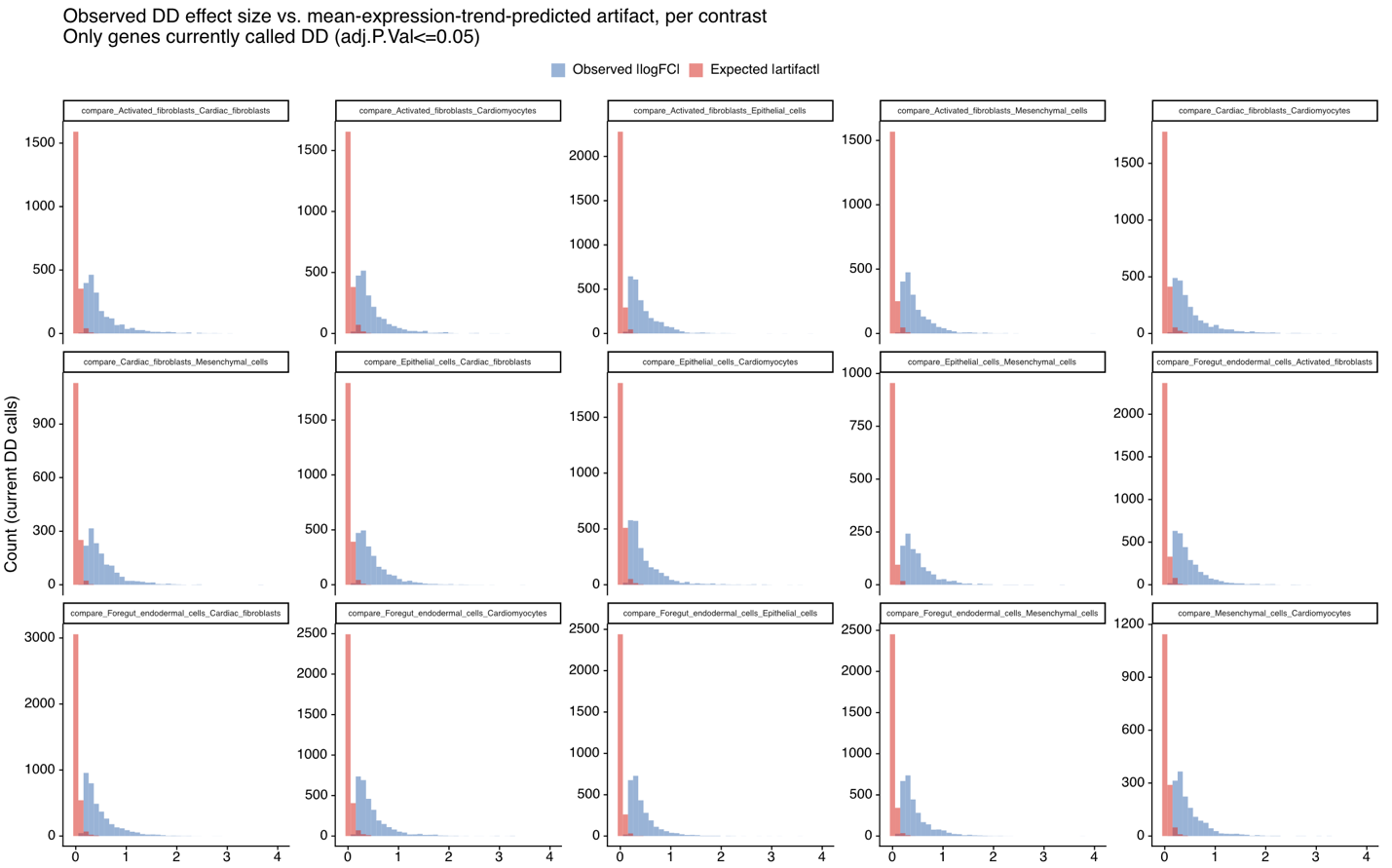

**Figure S42.**  For each of the 15 pairwise cell-type contrasts, overlaid histograms of observed absolute effect sizes (blue) vs. the effect size the mean-expression LOESS trend alone would predict for that gene pair ("expected artifact", red), restricted to genes currently called DD (adj.P.Val ≤ 0.05).

**Supplementary Table S1 (provided as a separate file).** Per gene estimates of log-dispersion and log-mean per sample (indicated by donor and cell type annotation in the column names) from the cardiac HDC data as estimated by Memento. Provide log.total.variability, log.mean.component.of.variance, and the percent contribution to a gene’s total variability due to dispersion and the mean effect, respectively.

**Supplementary Table S2 (provided as a separate file).** Per gene estimates of log-dispersion and log-mean per sample (indicated by donor, species, and cell type annotation in the column names) from the interspecies HDC data as estimated by Memento. Provides log.total.variability, log.mean.component.of.variance, and the percent contribution to a gene’s total variability due to dispersion and the mean effect, respectively.

**Supplementary Table S3 (provided as a separate file).** Gene ontology enrichment. The first sheet contains enrichment results that were identified as low DD and high DE in each cell type comparison (indicated in the “contrasts” column). The second sheet contains enrichments for low-dispersion genes across cell types. The “cluster_resolution” column indicates at what resolution clustering was performed and dispersion was estimated to define low-dispersion genes. In both sheets, all other columns are standard outputs from clusterProfiler.

**Supplementary Table S4 (provided as a separate file).** All DE and DD results for both cardiac HDC and interspecies datasets (per gene effect sizes, standard errors, significance results). Detailed descriptions of each field provided in the file.

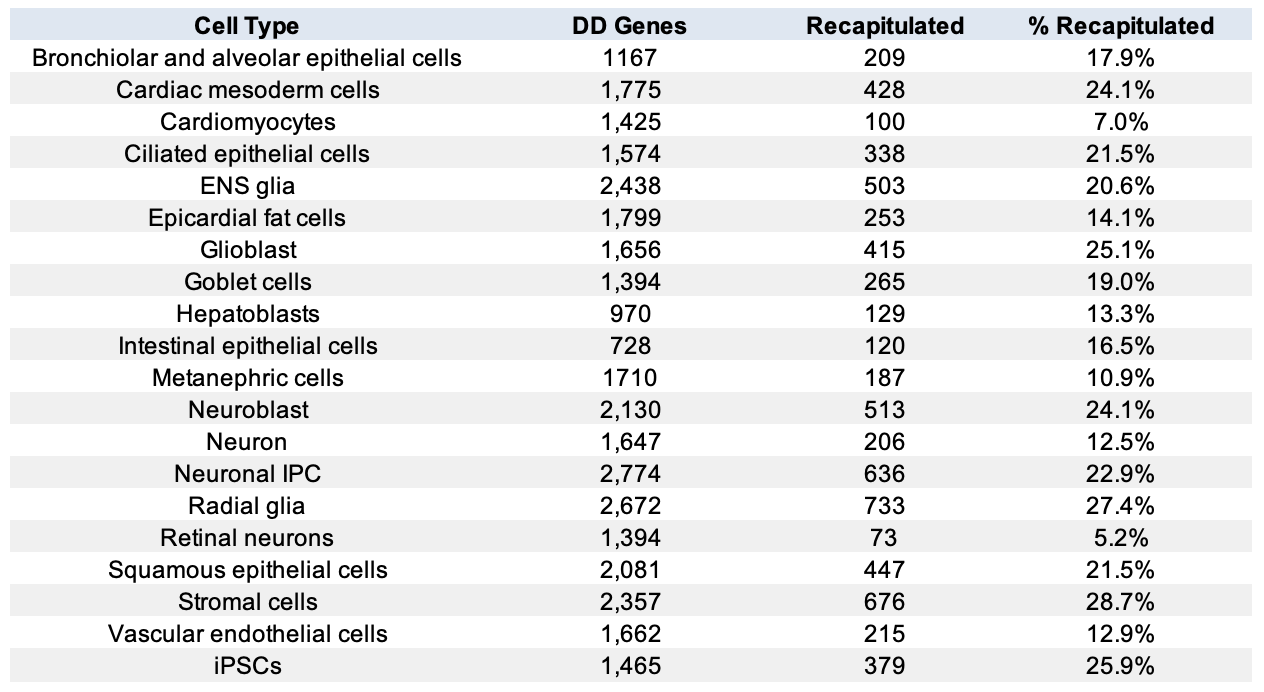

**Supplementary Table S5.** Number of genes per cell type in the diploid lines with significant species differences in dispersion (LFSR < 0.05). Number and percent of DD genes that are recapitulated (namely, regulated in *cis*) in the allotetraploid line.

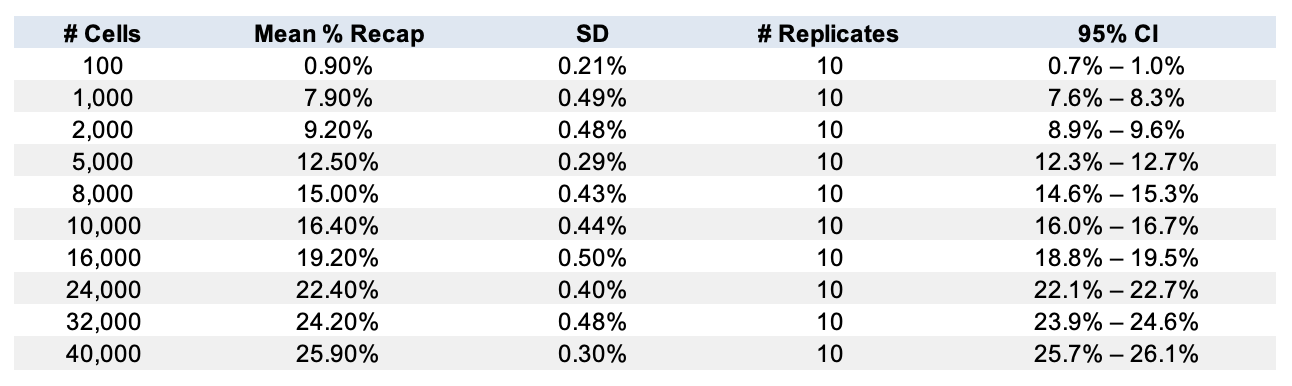

**Supplementary Table S6.** Summary table of downsampling of allotetraploid cells to assess recapitulation rate (Figure S26).

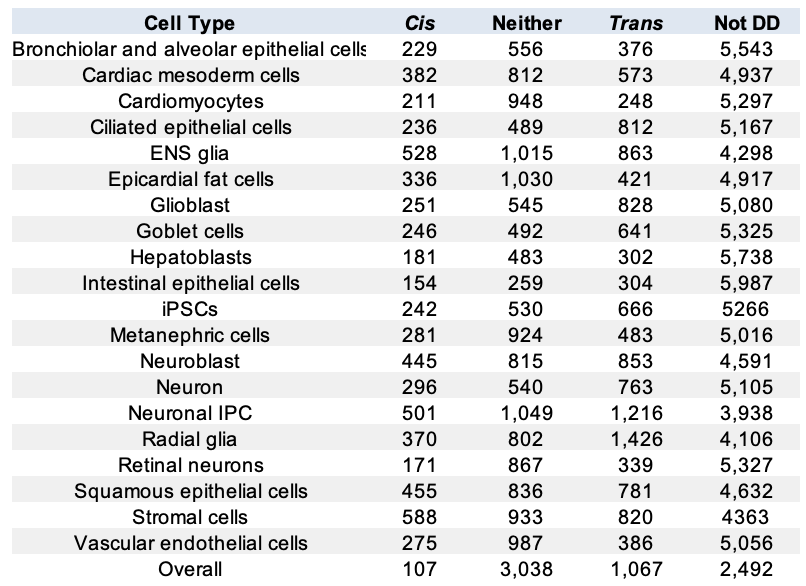

**Supplementary Table S7.** Number of DD genes that are classified as *cis*, *trans*, or neither (see Methods) per cell type. DD genes are classified as *cis* if the estimated cis proportion exceeds 0.7, as *trans* if it is below 0.3, and as *neither* otherwise. In the ‘overall’ analysis, which considered all cell type, ‘mixed’ refers to genes whose DD was accounted for by *cis* in at least one cell type and *trans* in at least one other cell type. The total number of genes per cell type (*cis* + *trans* + neither) equals the number of DD genes observed in the diploid lines (‘DD genes’ in Table S4).

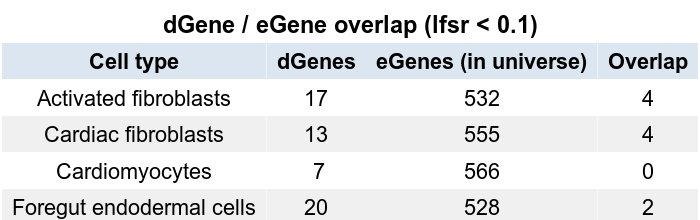

**Supplementary Table S8.** Overlap between dispersion-QTL genes (dGenes, lfsr < 0.1) and mean-expression-QTL genes (eGenes; McIntire et al’s independent eQTL mapping analysis), within the shared tested-gene universe, per cell type.
